# Multi-level broad-yet-sparse input organization of LC-NE neurons revealed by multiplexed whole-brain EM reconstruction

**DOI:** 10.1101/2025.06.12.659365

**Authors:** Fu-Ning Li, Jia-Zheng Liu, Can Shi, Jing-Bin Yuan, Ya-Nan Lv, Jing Liu, Li-Na Zhang, Lin-Lin Li, Li-Jun Shen, Xi Chen, Hao Zhai, Yan-Chao Zhang, Rong-Kun Tao, Han-Yang Hu, Fang-Xu Zhou, Tong Xin, Bo-Hao Chen, Hao-Ran Chen, Sheng Chang, Hong-Tu Ma, Hai-Yang Yan, Jia-Liang Zu, Jin-Yue Guo, Zheng-Huan Fang, Xiao-Hui Dong, Li-Mei Lin, Xing-Hui Zhao, Chen-Ying Qian, Yan-Song Wang, Shi-Rong Jin, Chang-Mei Zhang, Pei-Shan Xiang, Yong-Xin Yang, Yu Qian, Yu-Chen Gong, Xu-Fei Du, Qi-Wei Xie, Hua Han, Jiu-Lin Du

## Abstract

Diverse neuromodulatory systems confer functional flexibility upon the hardwired sensorimotor pathways of the brain^1–6^. Among these, the evolutionarily conserved locus coeruleus (LC)-norepinephrinergic (NE) system integrates and broadcasts information globally^7–9^. While previous studies have primarily focused on its axonal outputs and effects on neural processing, the organizational logic of its synaptic inputs, enabling it to sense global brain states and in turn shape its activity dynamics for appropriate neuromodulation^10,11^, remains poorly explored. To address this, we mapped the synaptic input architecture of individual LC-NE neurons by establishing Fish-X, a whole-brain microscale reconstruction of larval zebrafish with neuron-type annotations. This dataset encompasses the retina, brain and anterior spinal cord, capturing >240,000 cells and >25 million synapses. Monoaminergic (including NE, dopaminergic, and serotonergic), hypocretinergic, and glycinergic neurons were resolved by multiplexed subcellular APEX2 labeling, while glutamatergic and GABAergic identities were inferred via morphology comparison with a zebrafish mesoscopic atlas^12,13^. Compared with other neuronal populations, LC-NE neurons display distinct perisomatic features and high dendritic indegrees. Reconstruction of near-complete dendritic inputs to individual LC-NE neurons reveals their broad yet sparse synaptic convergence across the brain. These synaptic inputs are not randomly distributed but instead organized according to sensory/motor modality, excitatory/inhibitory identity, and synaptic strength. Individual LC-NE neurons share common inputs, a feature conserved within and across monoaminergic systems, suggesting a co-innervation mechanism for coordinated neuromodulation. Thus, our study uncovers multi-level principles governing the spatial organization of LC-NE neurons’ broad-yet-sparse inputs and provides a pivotal resource for deciphering the microscale architecture of neuromodulatory systems.

---

The brain is organized across scales, from precisely wired synapses, through morphologically and functionally heterogeneous neurons, to specific neural circuits and brain-wide networks^14–16^. A simplified framework for deconstructing its organization is the concept of a bi-pathway architecture, in which inter-connected sensorimotor and neuromodulatory pathways function complementarily^1–6^. Neuromodulatory systems orchestrate sensorimotor circuits, shaping their state, dynamics and output without directly driving behavioral outputs. While previous studies mainly focused on their axonal outputs and influence on sensorimotor processing^7,10,17–21^, a critical question remains: how do these systems, through their synaptic inputs, perceive external environmental changes as well as internal brain states and in turn generate dynamic activities enabling spatiotemporally specific modulation^1–3,5,6,10,11^? To address this issue, it is essential to elucidate the spatial organization of synaptic inputs onto neuromodulatory neurons, including cellular origin, functional modality, and excitatory (E)/inhibitory (I) identity.

Microscale connectomics based on electron microscopy (EM) is a powerful approach for synaptic input mapping^22–27^, yielding wiring diagrams for the central nervous system (CNS) of small invertebrates (including *Caenorhabditis elegans* and *Drosophila melanogaster*)^28–33^ and cubic millimeter-scale reconstructions of the mouse and human cortices^34–38^. The larval zebrafish (*Danio rerio*) with its manageable brain scale has emerged as a tractable vertebrate model for whole-brain reconstruction^39^. Previous studies have provided foundational datasets, allowing the synaptic-resolution mapping of sensorimotor circuits^40–45^. The functional complexity of neural networks arises from both the physical wiring and molecular diversity of their constituent neurons. However, the absence of molecular annotations, particularly for neuromodulatory neurons, limits functional insights from most connectomes. While traditional EM connectomics identifies neuron types using structural landmarks, mapping cell types that lack distinct anatomical signatures requires correlative light-electron microscopy (CLEM)^45^, immunogold labeling^46^, or genetically encoded tags for EM (GETEM)^47^. Among these, GETEM offers a highly feasible approach, as it leverages genetically encoded tags to embed molecular identity directly within the ultrastructural map, sidestepping computational registration challenges and ensuring sample preservation.

We therefore built Fish-X, a whole-brain, synaptic-resolution serial-section EM (ssEM) dataset with multiplexed molecular annotations from a 6-day post-fertilization (dpf) larval zebrafish, spanning the CNS from the retina to the anterior spinal cord (aSC). Within this dataset, diverse neuromodulatory (norepinephrinergic, NE; dopaminergic, DA; serotonergic, 5HT; hypocretinergic, Hcrt) and inhibitory glycinergic (Gly) neurons were precisely identified using enhanced ascorbate peroxidase 2 (APEX2)^48^, which was targeted to different subcellular compartments of distinct neuron types using signal peptides and neuron type-specific promoters. Furthermore, we inferred glutamatergic (Glu) and GABAergic (GABA) types of neurons through the zebrafish mesoscopic atlas^12^-assisted morphology comparison. A whole-brain algorithmic reconstruction was then performed, identifying 176,810 cells and 25.25 million synapses, thereby providing a foundation for systematic analysis of the organization of synaptic inputs to these neuromodulatory neuronal populations.

We focused on NE neurons in the evolutionarily conserved locus coeruleus (LC), a key nuclei for brain-wide regulation^49–53^. Viral tracing revealed cellular-level input convergence onto LC-NE neurons^7,8^, yet the spatial organization of synaptic-level inputs remains unknown. To address this, we performed somatodendritic reconstructions and synaptic mapping of APEX2-labeled LC-NE neurons. These reconstructions enabled us to define their characteristic perisomatic ultrastructural features, establishing structural criteria for the precise, label-free identification of LC-NE neurons^54,55^. Brain-wide, synaptic-resolution input reconstruction revealed that individual LC-NE neurons integrate widely distributed inputs. These inputs were spatially organized by functional modality and E/I type, with sensory-related I inputs biasing to distal dendrites and motor-related E inputs to proximal ones. While most inputs are unique to individual neurons, a subset constitutes intra-system common inputs that innervate multiple LC-NE neurons. Moreover, this co-innervation principle extends across different monoaminergic systems, suggesting an inter-system organization for orchestrating NE, DA, and 5HT release, which may collectively modulate neural network activities. Take together, leveraging the Fish-X resource, a whole-brain, synaptic-resolution dataset with molecularly defined neuron-type annotations, we have quantified the ultrastructural features of diverse neuromodulatory neurons and delineated the organizational principles governing synaptic inputs to the LC-NE system. Integrated with the zebrafish mesoscopic atlases^12,13^, the Fish-X can serve as a valuable resource for deciphering how, at the level of synaptic architecture, neuromodulatory systems interact with sensorimotor pathways to shape neurocomputing in the larval zebrafish brain.

## Multiplexed APEX2-labeled whole-brain ssEM of larval zebrafish

We achieved simultaneous EM labeling of multiple molecularly defined neuron types within a single brain preparation using APEX2, an enzyme that catalyzes the hydrogen peroxide (H2O2)-dependent localized polymerization of diaminobenzidine (DAB)^48^. APEX2 fused with signal peptides and fluorescent proteins was expressed via cell type-specific promoters for labeling distinct subcellular compartments in different types of neurons. After the DAB reaction, the localized APEX2 generated EM-visible staining patterns, thereby enabling the identification of different neuron types (Fig. 1a; see Methods). We achieved EM labeling of multiple subcellular compartments including the nucleus, cytosol, plasma membrane, mitochondria, Golgi apparatus, and presynaptic sites (Fig. 1b, six leftmost images, and Extended Data Fig. 1a). In addition, by fusing APEX2 with self-assembling proteins, Polymer King-size Unit (PKU) tags^56^, we achieved fibrous and spherical staining patterns localized in the nucleus or cytosol (Fig. 1b, three rightmost images, and Extended Data Fig. 1a). Thus we generated a versatile genetic toolbox to target a wide range of neuromodulatory types, including NE, DA, 5HT, and Hcrt neurons, as well as Glu, GABA, and Gly neurons across the CNS (Supplementary Table 1, 2).

**Fig. 1.**
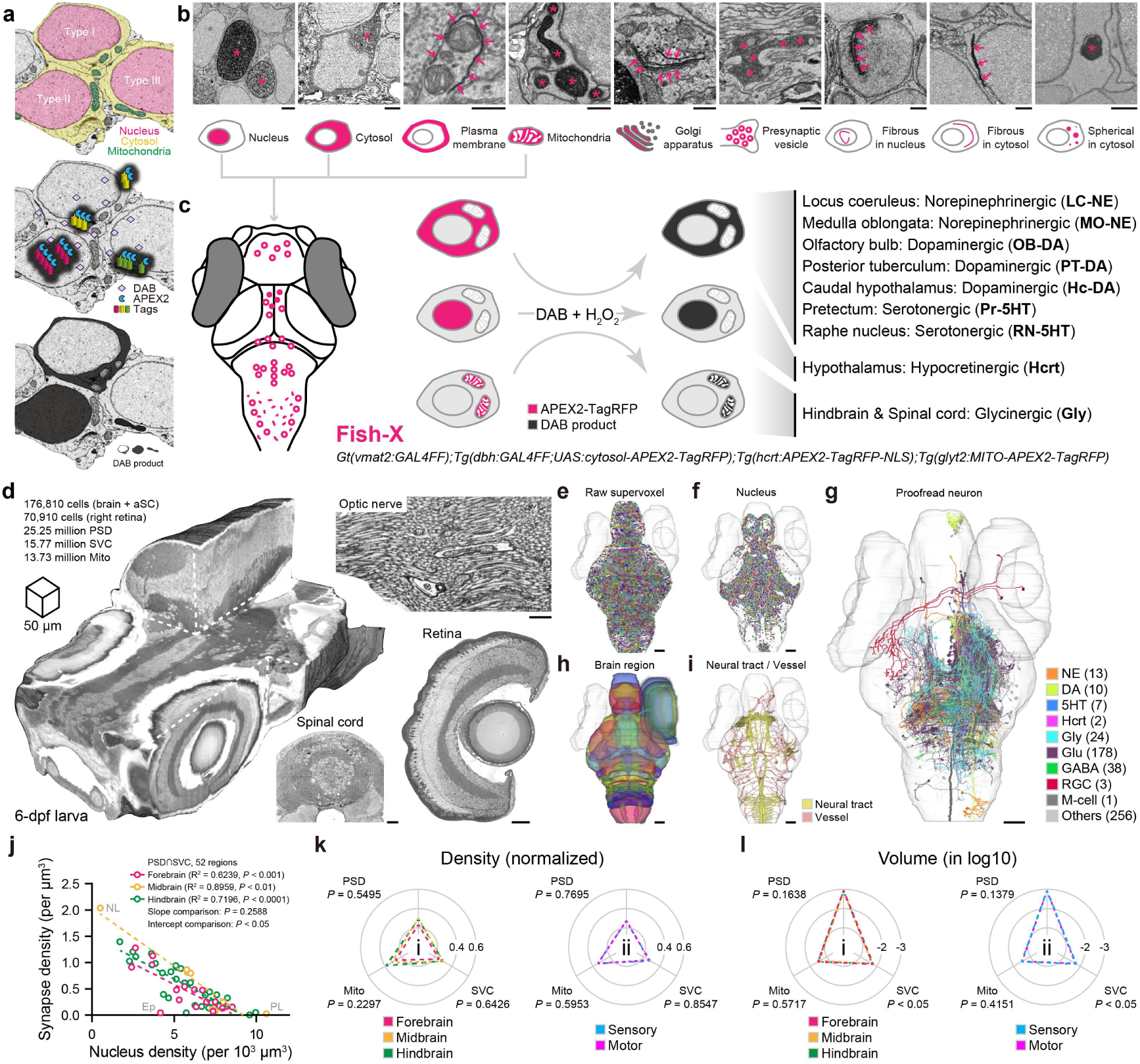
Multiplexed neuromodulatory-type-labeled whole-brain ssEM of larval zebrafish. **a-c,** Neuron-type annotation via multiplexed subcellular peroxidase labeling. (**a**) Schematic of APEX2-based multiplexed EM labeling strategy. Assuming three adjacent neurons of distinct types (top) have APEX2 targeted to different subcellular compartments via specific signal peptides and promoters (middle). Following the DAB reaction, APEX2 catalyzes the conversion of DAB into osmiophilic substances and generates localized non-transmembrane electron-dense staining (bottom), which enables the identification of neuron types in the EM dataset. (**b**) Nine distinct labeling patterns were generated by localizing APEX2 (red) to different compartments (six leftmost images), including nucleus, cytosol, plasma membrane, mitochondria, Golgi apparatus, and presynaptic vesicle. Fusion of APEX2 with PKU tags yielded additional fibrous or spherical self-assembled aggregates in the nucleus and cytosol (three rightmost images). Labeling patterns are indicated by red stars or arrows. (**c**) *Gt(vmat2:GAL4FF);Tg(dbh:GAL4FF;UAS:cytosol-APEX2-TagRFP);Tg(hcrt:APEX2-TagRFP-NLS);Tg(glyt2:MITO-APEX2-TagRFP)*, a quadruple transgenic fish line, was established to enable the collection of the Fish-X dataset. Combined with soma location, cytosol, nucleus, and mitochondria labeling patterns were used to identify monoaminergic, Hcrt, and Gly neurons, respectively. The distinct neuron types are listed on the right. **d,** Overview of the multiplexed neuromodulatory-type-annotated whole-brain ssEM dataset (“Fish-X”) of a 6-dpf zebrafish larva, comprising three partially overlapping volumes: the brain and aSC, the right retina, and the right optic nerve. This panel shows the rendering of the intact head (left; 992×992×660 nm^3^ per voxel) and representative EM images of the aSC (bottom middle; 32×32 nm^2^ per pixel), right optic nerve (top right; 8×8 nm^2^ per pixel resolution), and right retina (bottom right; 32×32 nm^2^ per pixel). **e-i,** Multidimensional automated reconstruction or manually proofread results. (**e**) Rendering of 46,536 automatically reconstructed supervoxels in the brain and aSC (dorsal view), with length ranging between 100 – 300 μm. (**f)** Rendering of randomly color-coded nuclei distributed in a representative horizontal plane at a depth of 108 μm from the dorsal epidermal surface. (**g)** Rendering of the partially reconstructed morphologies for 523 neurons following manual verification. These include 13 NE neurons (orange) in the LC (*n* = 10) and MO (*n* = 3), 10 DA neurons (light green) in the OB (*n* = 3), PT (*n* = 4) and Hc (*n* = 3), 7 5HT neurons (dark blue) in the Pr (*n* = 3) and RN (*n* = 4), 2 Hcrt neurons (magenta) in the hypothalamus (magenta), 24 Gly neurons (sky blue) in the hindbrain, 178 Glu neurons (purple), 38 GABA neurons (green), 3 RGCs (red), 1 M-cell (dark gray), and 256 type-unidentified neurons (light gray; Among them, 9 were PNs to a single OB-DA neuron). (**h)** A total of 52 brain regions and 7 retinal regions were annotated using the CLEM approach, followed by manual verification. (**i)** Rendering of brain-wide neural tracts (yellow) and vessels (red). Most vessels remained well-preserved, though some collapse was observed in the telencephalon. Neural tracts, identified as a bundle of parallel-oriented fibers, were manually traced. Vessels were automatically reconstructed, followed by manual proofreading. **j-l,** Quantification of nucleus and synapse distribution across brain regions. (**j**) Linear regression analysis of synapse density versus nucleus density within each of the forebrain (red), midbrain (yellow), and hindbrain (green) regions. Dashed lines represent the fitted regression lines, respectively (Forebrain: Y = −158.1 × X + 1,373, R^2^ = 0.6239, *P* < 0.001; Midbrain: Y = −221.1 × X + 2,037, R^2^ = 0.8959, *P* < 0.01; Hindbrain: Y = −165.0 × X + 1,513, R^2^ = 0.7196, *P* < 0.0001). Each circle represents a single region. For analysis, a synapse was defined by the co-occurrence of the PSD and SVC. The observed excess of putative synapses (23.62 million) over SVCs (15.77 million) arises because multiple PSDs often associate with a single presynaptic SVC. The normalized density (**k**) and volume in log10 (**l**) of PSD, SVC, and Mito were analyzed across anatomical and functional groupings. For each structure, the density in a given region was normalized to the highest density of that structure across regions. The volume was log10-transformed for analysis. Brain regions were categorized by anatomy (forebrain, red; midbrain, yellow; hindbrain, green) and function (sensory, blue; motor, magenta). The dashed line in each radar plot represents the mean value across all corresponding structures for each group. Panel i in (**k**): *P* (PSD) = 0.5495, *P* (SVC) = 0.6426, *P* (Mito) = 0.2297, Kruskal-Wallis test; Panel ii in (**k**): *P* (PSD) = 0.7695, *P* (SVC) = 0.8547, *P* (Mito) = 0.5953, Mann Whitney test. Panel i in (**l**): *P* (PSD) = 0.1638, *P* (SVC) < 0.05, *P* (Mito) = 0.5717, Kruskal-Wallis test with Dunn’s multiple comparison test; Panel ii in (**l**): *P* (PSD) = 0.1379, *P* (SVC) < 0.05, *P* (Mito) = 0.4151, Mann Whitney test. C, caudal; L, left; V, ventral; LC, locus coeruleus; MO, medulla oblongata; OB, olfactory bulb; PT, posterior tuberculum; Hc, caudal hypothalamus; Pr, pretectum; RN, raphe nucleus; NL, neuropil layers (optic tectum); NE, noradrenergic; DA, dopaminergic; 5HT, serotonergic; Hcrt, hypocretinergic; Gly, glycinergic; Glu, glutamatergic; GABA, GABAergic; RGC, retinal ganglion cell; M-cell, Mauthner cell; PSD, postsynaptic density; SVC, synaptic vesicle cloud; Mito, mitochondria. Scale bars, 0.5 µm (**b**), 1 µm (**d**, optic nerve), 20 µm (**d**, spinal cord and retina), and 50 µm (**e-i**).

To examine the inputs and their spatial organization of multiple neuromodulatory neuronal populations within the same animal, we made a quadruple transgenic zebrafish line, *Gt(vmat2:GAL4FF);Tg(dbh:GAL4FF;UAS:cytosol-APEX2-TagRFP);Tg(hcrt:APEX2-TagRFP-NLS);Tg(glyt2:MITO-APEX2-TagRFP)*, in which we were able to simultaneously annotate cytosol-labeled NE neurons in the LC and medulla oblongata (MO), cytosol-labeled DA neurons in the olfactory bulb (OB), posterior tuberculum (PT) and caudal hypothalamus (Hc), cytosol-labeled 5HT neurons in the pretectum (Pr) and raphe nucleus (RN), nucleus-labeled Hcrt neurons in the hypothalamus, and mitochondria-labeled Gly neurons in the hindbrain and aSC (Fig. 1c and Extended Data Fig. 1b). The specificity of labeling was confirmed through immunohistochemical staining or crossing with neuron type-specific reporter fish lines, albeit with variations in labeling efficiency across neuronal populations (Extended Data Fig. 1c).

We acquired a whole-brain ssEM dataset (named as Fish-X) from a 6-dpf multiplexed APEX2-labeled larva using the automated tape-collecting ultramicrotome – scanning electron microscopy (ATUM-SEM) approach^40^. With a voxel resolution of 4×4×33 nm^3^, Fish-X extended from the olfactory epithelium to the aSC (the vertebra 1) and included the right retina and optic nerve (Fig. 1d, Extended Data Fig. 2; see Methods). The imaged volume, generated from 22,887 consecutive coronal-view ultrathin sections, encompassed ∼3.1160×10^7^ μm^3^ for the entire brain and aSC, ∼5.5620×10^4^ μm^3^ for the right optic nerve, and ∼6.0877×10^6^ μm^3^ for the right retina. Utilizing an automated three-dimensional (3D) convolutional neural network (CNN)-based reconstruction of the brain and aSC (Fig. 1e; see Methods), we performed multidimensional structural annotations (Supplementary Video 1) encompassing nuclei (Fig. 1f and Extended Data Fig. 3), neuronal morphologies (Fig. 1g), brain regions (Fig. 1h, Supplementary Table 3, and Supplementary Video 2), neural tracts and vessels (Fig. 1i).

Based on the results of nucleus segmentation, a total of 176,810 cells, including 66 NE, 105 DA, 126 5HT, 12 Hcrt, and 2,689 Gly neurons, were identified in the brain and aSC, and 70,910 cells in the right retina (see Fig. 1d). We quantified nuclear morphology by integrating deep-learned and traditional features (see Methods) and discovered that morphological diversity formed a continuous spectrum, though vasculature-associated cell types, such as endothelial cells (ECs) and pericytes (PCs), clustered distinctly within the morphological space (Extended Data Fig. 3a,b). Spatial mapping revealed heterogeneity in cell density across brain regions (Extended Data Fig. 3c, Supplementary Table 4, and Supplementary Video 2).

In a parallel analysis of synaptic structures (Extended Data Fig. 4a; see Methods), automatic segmentation identified 25.25 million postsynaptic densities (PSDs), 15.77 million synaptic vesicle clouds (SVCs), and 13.73 million mitochondria (Mito) in the brain and aSC. While 93.5% (56.6% with SVC and Mito + 36.9% with SVC only) of PSDs colocalized with SVCs, only 59.2% (56.6% with SVC and Mito + 2.6% with Mito only) colocalized with presynaptic mitochondria (Extended Data Fig. 4b), suggesting heterogeneity in synaptic energy supply. Meanwhile, 84.3% of SVCs and 61.3% of mitochondria co-localized with PSDs (Extended Data Fig. 4c,d). All three structures exhibited heterogeneous yet highly concerted distribution patterns across regions (Extended Data Fig. 4e-j, Supplementary Tables 5,6,7, and Supplementary Video 2).

Across regions, the density of typical synapses with colocalized PSDs and SVCs was inversely correlated with cell density, which was consistently observed across forebrain, midbrain, and hindbrain, highlighting the spatial segregation of synapses and somata (Fig. 1j). When regions were grouped into broader anatomical (i.e., forebrain, midbrain, and hindbrain) or functional (i.e., sensory-related, and motor-related) categories, no significant difference in density was found (Fig. 1k; Kruskal-Wallis test for the anatomical category, Mann Whitney test for the functional category). Similarly, the volumes of all three components followed comparable distributions across regions, with the exception of a slight difference in SVC volume between the forebrain and hindbrain (Fig. 1l; Kruskal-Wallis test with Dunn’s multiple comparison test for the anatomical category, Mann Whitney test for the functional category). Regions from the same anatomical categories generally exhibited similar lognormal distributions of synaptic structure volumes (Extended Data Fig. 4k-n).

Collectively, we developed a whole-brain, synaptic-resolution dataset for larval zebrafish, annotated with neuromodulatory and glycinergic neurons via multiplexed subcellular APEX2 labeling.

## Unique perisomatic features and high dendritic indegrees of LC-NE neurons

Different neuromodulatory systems vary in the degree of axonal innervation patterns^7,10,17–21^. As evidenced by light microscopy (LM)-based single-cell morphological tracing in 6-dpf larval zebrafish, we found that LC-NE neurons exhibited more widespread axonal projections throughout the brain in comparison with other types of neuromodulatory neurons, including MO-NE, OB-DA, PT-DA, Hc-DA, Pr-5HT, RN-5HT, and Hcrt neurons (Extended Data Fig. 5). To characterize the ultrastructural properties of these neurons’ synaptic inputs, we reconstructed the somatodendritic morphology and synaptic input sites of these types of neurons at the right hemisphere, along with Gly neurons, and proofread and quantified their perisomatic ultrastructures (Fig. 2a-g, Extended Data Figs. 6,7, and Supplementary Table 8).

**Fig. 2.**
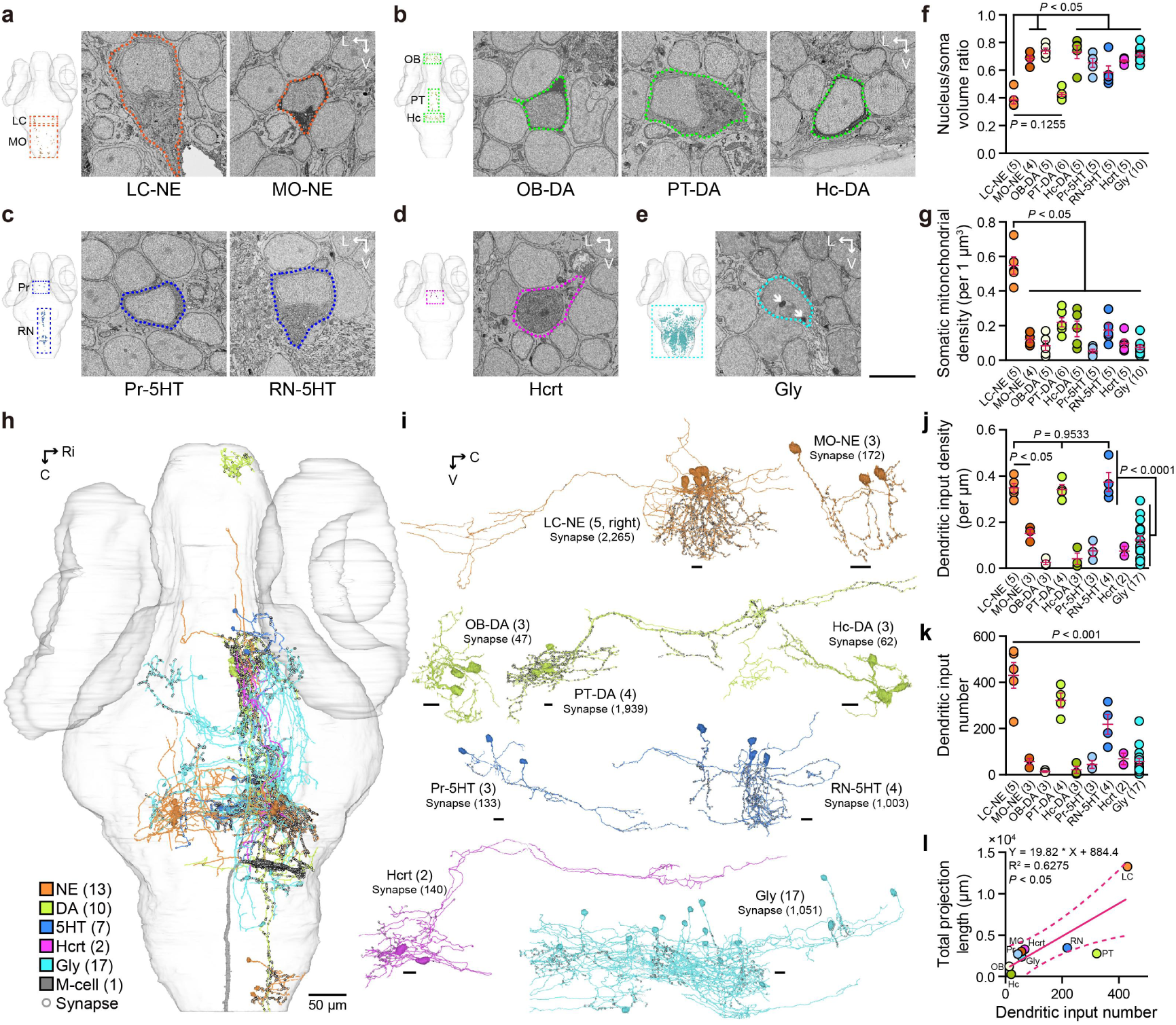
Reconstruction of perisomatic ultrastructures and dendritic synaptic inputs of multiple neuromodulatory neurons. **a-e,** Representative EM images (4×4 nm^2^ per pixel) of seven molecularly identified neuron types with APEX2 labeling (NE, orange; DA, green; 5HT, blue; Hcrt, magenta; Gly, cyan). Individual APEX2-labeled neurons display various degrees of staining intensity and are contoured by color-coded dashed lines. Renderings of all APEX2-labeled nuclei are shown on the left, with brain regions outlined by dashed squares. All types except Gly were manually verified. Gly neurons in hindbrain were identified automatically based on the presence of ≥4 labeled mitochondria in the soma. White arrows point to two APEX2-labeled mitochondria in a Gly neuron, with one adjacent to the nuclear envelope infoldings (**e**). **f,g,** The mean nucleus/soma volume ratio (**f**) and somatic mitochondrial density (**g**) were analyzed across neuron types. (**f**) LC-NE *vs* PT-DA, *P* = 0.1255; LC-NE *vs* all other types except PT-DA, *P* < 0.05. (**g**) LC-NE *vs* all other types, *P* < 0.05. The Mann Whitney test was used. **h,i,** Rendering of EM reconstructions for APEX2-labeled neuromodulatory and Gly neurons in the right hemisphere, shown in dorsal (**h**) and lateral (**i**) views. Gly neurons with APEX2 labeling (cyan), identified as LC-NE presynaptic neurons during reconstruction, are also shown. Presynaptic input sites are marked as small gray dots. The dendritic arbors were fully reconstructed for neuromodulatory types. Neuron and synapse numbers are shown in parentheses. Input mapping was not performed for five LC-NE neurons on the left sides. **j,k,** Dendritic input density (**j**) and number (**k**) across neuron types. Dendritic input density (per μm): LC-NE, 0.3493 ± 0.0194; MO-NE, 0.1511 ± 0.0185; OB-DA, 0.0255 ± 0.0094; PT-DA, 0.3420 ± 0.0212; Hc-DA, 0.0415 ± 0.0249; Pr-5HT, 0.0759 ± 0.0249; RN-5HT, 0.3749 ± 0.0413; Hcrt, 0.0747 ± 0.0204; Gly, 0.1228 ± 0.0190. *P* (LC *vs* MO) < 0.05 (Mann Whitney test); *P* (among LC, PT, and RN) = 0.9533 (Kruskal-Wallis test); *P* (LC/PT/RN *vs* all other types) < 0.0001 (unpaired *t* test). Dendritic input number: LC-NE, 430.2 ± 55.48; MO-NE, 56.67 ± 13.35; OB-DA, 13.67 ± 3.180; PT-DA, 321.8 ± 31.89; Hc-DA, 20.67 ± 15.64; Pr-5HT, 44.00 ± 16.52; RN-5HT, 218.0 ± 43.18; Hcrt, 68.00 ± 25.00; Gly, 57.47 ± 13.74. *P* (among all neuron types) < 0.01 (Kruskal-Wallis test). **l,** Linear regression analysis of the total projection length versus the dendritic input number (Y = 19.82 × X + 884.4, R^2^ = 0.6275, *P* < 0.05). The total projection length was calculated as the mean of three individual neurons from mesoscopic datasets for each neuron type. The dendritic input number were calculated from the same neurons in h to l. Ro, rostral; C, caudal; L, left; Ri, right; V, ventral. Error bars represent mean ± SEM. Scale bars, 5 µm (**a-e**), 10 µm (**i**), and 50 µm (**h**).

LC-NE neurons exhibited not only larger nuclear and somatic volumes but also featured extensive endoplasmic reticulum (ER) networks, numerous mitochondria, and a specialized cilium (Extended Data Fig. 7a and Supplementary Table 8). Compared with a 10-fold pool of randomly selected adjacent cells, LC-NE neurons had a lower mean nucleus/soma volume ratio (LC-NE neurons *vs* adjacent cells: 0.4123 ± 0.0808 *vs* 0.6530 ± 0.0113, *P* < 0.05, Mann Whitney test) and a higher somatic mitochondrial density (LC-NE neurons *vs* adjacent cells: 0.5449 ± 0.0496 *vs* 0.0843 ± 0.0088 per 1 μm^3^, *P* < 0.001, Mann Whitney test) (Extended Data Fig. 7b, left). Clustering analyses integrating both nuclear and somatic characteristics validated the above observation (Extended Data Fig. 7b, right). Based on these distinct perisomatic features, we identified an additional unlabeled NE neuron in the right LC region (Extended Data Fig. 7b, blue arrows), whose morphology closely matched those of APEX2-labeled LC-NE neurons (Extended Data Fig. 7c). We extended these analyses to other APEX2-labeled neuronal populations and found that, with the exception of PT-DA neurons (Extended Data Fig. 7f), other types of neuromodulatory neurons lacked perisomatic features that clearly distinguished them from adjacent cells (Extended Data Fig. 7d-k and Supplementary Table 8). This finding further highlights the value of APEX2 labeling for precise identification of neuron types. In companion with other types of neuromodulatory neurons, LC-NE neurons exhibited higher values in terms of nucleus/soma volume ratio (Fig. 2f; LC-NE *vs* PT-DA, *P* = 0.1255, Mann Whitney test; LC-NE *vs* all other types except PT-DA, *P* < 0.05, Mann Whitney test) and somatic mitochondrial density (Fig. 2g; LC-NE *vs* all other types, *P* < 0.05, Mann Whitney test), suggesting unique perisomatic properties of LC-NE neurons.

We then reconstructed and quantified all synaptic input sites across 27 APEX2-labeled neuromodulatory neurons (including 5 LC-NE, 3 MO-NE, 3 OB-DA, 4 PT-DA, 3 Hc-DA, 3 Pr-5HT, 4 RN-5HT, and 2 Hcrt) and 17 labeled Gly neurons at somatodendritic regions (Fig. 2h,i and Supplementary Videos 3,4). LC-NE, PT-DA, and RN-5HT neurons collectively displayed a higher mean synaptic input number and density compared with other types, with the LC-NE neurons exhibiting the highest synaptic input number (Fig. 2j,k; Input density among LC-NE, PT-DA, and RN-5HT, *P* = 0.9533, Kruskal-Wallis test; Input density of LC-NE/PT-DA/RN-5HT *vs* all other types, *P* < 0.0001, unpaired *t* test; Input number, *P* < 0.0001, Kruskal-Wallis test). Moreover, the synaptic input number showed a positive correlation with the total projection length of respective neuron types (Fig. 2l). Notably, these neuromodulatory neurons had sparse synaptic inputs on their somata (Supplementary Video 4), unlike the well-studied Mauthner cell, which received substantial somatic innervation (Extended Data Fig. 8 and Supplementary Video 5).

Thus these results show that LC-NE neurons constitute a structurally distinct system with unique perisomatic properties and high dendritic indegrees. Furthermore, the Fish-X dataset provides a structural ground truth for multiple neuromodulatory neuron types, serving as an essential reference for their identification in other EM datasets lacking molecular annotations^54,55^.

## Spatial organization of broad-yet-sparse inputs to individual LC-NE neurons

To examine the spatial organization of synaptic inputs on LC-NE neurons, we focused on LC-NE neuron #3 due to its possession of the longest dendrites (∼1.812 mm) and the highest dendritic indegree (560 synapses) among the five APEX2-labeled LC-NE neurons in the right hemisphere (Fig. 3a,b and Extended Data Fig. 9).

**Fig. 3.**
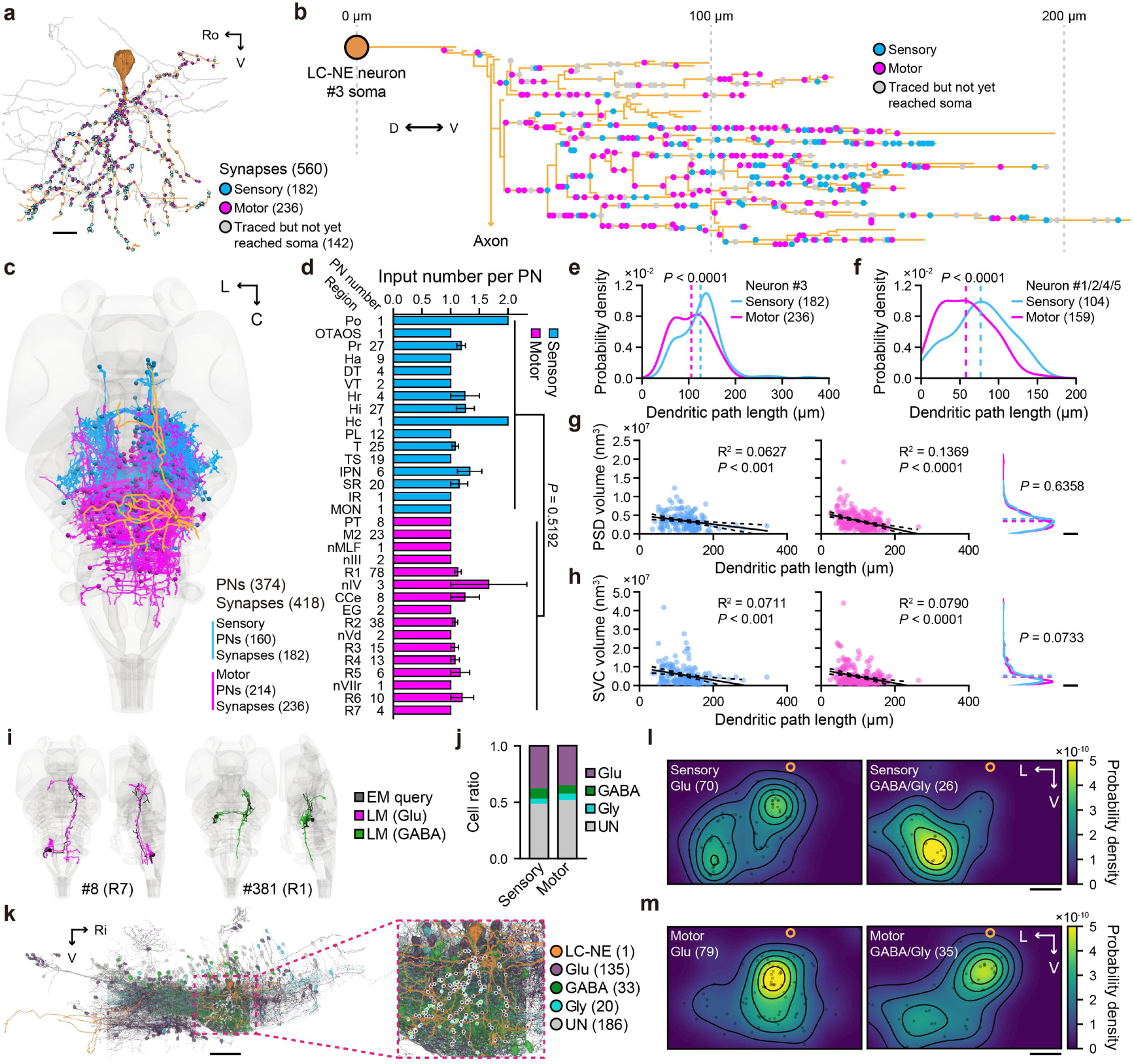
Spatial organization of broad-yet-sparse synaptic inputs of individual LC-NE neurons. **a,** Rendering of synaptic input sites on the dendrites of the LC-NE neuron #3, color-coded by sensory/motor modality. Synapses that were traced but not yet reached their soma are marked as gray dots. Dendritic and axonal processes are color-coded as orange and gray, respectively. The number in parentheses represent the number of synapses. **b,** 2D spatial distribution of synaptic input sites on the dendrites of LC-NE neuron #3. In zebrafish, LC-NE neurons are pseudounipolar neurons, with both axonal and dendritic processes emerging from a primary branch. The distance between each dot and the soma represents the path length of each input. **c,** Rendering of all 374 EM-reconstructed presynaptic neurons (PNs), which form 418 synapses on the LC-NE neuron #3 (orange). All presynapses for the LC-NE neuron #3 were traced and only those successfully traced back to their somata were used for analysis. Based on the soma locations of corresponding PNs, synaptic input sites were empirically classified as sensory-/sensory relay-related (blue; abbreviated as “sensory”) or motor-related (magenta). The same color scheme is applied to all related data below. The resulting counts are as follows: Sensory-related, 160 neurons and 182 synapses; Motor-related, 214 neurons and 236 synapses. **d,** The mean number of synapses formed by PNs in each of regions. Input number per PN: Sensory, 1.204 ± 0.08268; Motor, 1.102 ± 0.04288. *P* (sensory *vs* motor) = 0.5192 (Mann-Whitney test). Region name and PN number are shown on the left. The R1 and R2 rank as the two predominant input sources, accounting for 20.86% (78/374) and 10.16% (38/374) of total PNs, respectively. **e,f,** Differential probability distributions of sensory-/motor-related synaptic inputs along dendritic path length, from the soma to distal region. Separate analyses of neuron #3 alone (**e**) and the pooled data from the other four neurons (**f**: #1, #2, #4, and #5; Sensory-related synapse, *n* = 104; Motor-related synapse, *n* = 159) both revealed that motor-related inputs were distributed more proximally than sensory-related inputs. The mean distance to the soma are indicated by dashed lines. \*\*\*\**P* < 0.0001 (Mann Whitney test). **g,h,** Linear regression analyses of dendritic path length and the corresponding synaptic strength quantified by PSD volume (**g**) and SV volume (**h**). Sensory-related (blue): PSD volume *vs* path length (**g**, left), Y = −1.181e4 × X + 4.927e6, R^2^ = 0.0627, *P* < 0.001; SV volume *vs* path length (**h**, left), Y = −3.283e4 × X + 9.356e6, R^2^ = 0.0711, *P* < 0.001. Motor-related (magenta): PSD volume *vs* path length (**g**, middle), Y = −2.174e4 × X + 5.7e6, R^2^ = 0.1369, *P* < 0.0001; SV volume *vs* path length (**h**, middle), Y = −3.611e4 × X + 8.414e6, R^2^ = 0.0790, *P* < 0.0001. Right: probability distributions of PSD (**g**) and SV (**h**) volumes for both sensory-related (blue) and motor-related (magenta) inputs, with dashed lines indicating the mean volumes (Mann Whitney test). **i,** Representative examples showing partially EM-reconstructed query neurons (black) with their fully LM-reconstructed top-ranked E/I counterparts (Glu, magenta; GABA, green), revealed by morphology comparison. The index and soma location of the EM query were listed below. **j,** E/I composition ratio of PNs innervating the LC-NE neuron #3. The E/I type was determined for about half of sensory-related (*n* = 82) and motor-related (*n* = 106) PNs. Glu (purple, *n* = 135) and GABA (green, *n* = 33) neurons were inferred through morphology comparison, and Gly (blue) neurons were verified via mitochondrial APEX2 labeling (*n* = 17) or inferred through morphology comparison (*n* = 3). PNs with type unknown (UN, *n* = 186) are displayed as gray. **k,** Rendering of all 374 PNs of the LC-NE neuron #3, color-coded by sensory/motor modality (LC-NE, orange; Glu, purple; GABA, green; Gly, blue; UN, gray). Inset (right): highlighted somatodendritic area with synaptic input sites. **l,m,** Distribution of E (left) and I (right) input sites on the dendritic area of the LC-NE neuron #3, for sensory-related (**l**) and motor-related (**m**) inputs. Orange circles represent the soma. The values are derived from the number of synapses (shown in parentheses) formed by PNs identified as E or I in (**j**) and (**k**). Sensory-related I inputs (**l**, right) and motor-related E inputs (**m**, left) were more clustered in distal and proximal areas, respectively. Left-right axis: *P*(sensory-E *vs* sensory-I) < 0.05, *P*(motor-E *vs* motor-I) = 0.1072, *P*(sensory-E *vs* motor-E) < 0.001, *P*(sensory-I *vs* motor-I) < 0.05, *P*(sensory-E *vs* motor-I) = 0.6938, *P*(sensory-I *vs* motor-E) < 0.0001. Dorsal-ventral axis, *P*(sensory-E *vs* sensory-I) = 0.2084, *P*(motor-E *vs* motor-I) = 0.1809, *P*(sensory-E *vs* motor-E) = 0.0530, *P*(sensory-I *vs* motor-I) = 0.2053, *P*(sensory-E *vs* motor-I) = 0.6938, *P*(sensory-I *vs* motor-E) < 0.05. The two-sample Kolmogorov-Smirnov test was used. Ro, rostral; C, caudal; L, left; Ri, right; V, ventral. Scale bars, 10 µm (**a**), 20 µm (**l,m**), 50 µm (**k**), and 0.5×10^-7^ (**g,h**).

Presynaptic neurons (PNs) for this LC neuron were traced from all input sites and categorized by putative functional modality (i.e., sensory-related *vs* motor-related) based on PN soma location. We successfully traced 418 inputs to the somata of 374 PNs, averaging 1.12 inputs per PN. Among these, 182 synapses originated from 160 PNs in sensory-related regions (including sensory-relay regions), and 236 synapses from 214 PNs in motor-related regions (Fig. 3c). Spanning the forebrain, midbrain, and hindbrain, these PNs were found in 32 out of 52 total brain regions (Fig. 3d and Supplementary Video 6), indicative of a broad-yet-sparse connectivity pattern. The mean input number per PN did not differ significantly between sensory- and motor-related regions (Fig. 3d; Sensory *vs* Motor, *P* = 0.5192, Mann Whitney test), suggesting that both modalities conform to this connectivity pattern. Through a combinatorial analysis of dendritic position (relative to the LC-NE neuron’s soma) against PN functional modality, we found a modality-dependent spatial organization on the LC-NE neuron’s dendrites (Fig. 3e,f). Specifically, for LC-NE neuron #3, sensory-related inputs preferentially targeted distal dendritic segments, whereas motor-related ones were biased toward proximal segments (Fig. 3e; path length to the soma: sensory-related, 122.8 ± 3.2 μm *vs* motor-related, 103.3 ± 2.5 μm; *P* < 0.0001, Mann Whitney test). The similar organizational profiles were observed for the other four LC-NE neurons, though the synaptic sampling of these neurons was unintentionally biased toward the proximal dendrites, resulting in a shorter average path length (Fig. 3f; path length to the soma: sensory-related, 76.7 ± 3.6 μm *vs* motor-related, 57.8 ± 2.5 μm; *P* < 0.0001, Mann Whitney test). Furthermore, both sensory- and motor-related inputs exhibited a spatial gradient of synaptic strength, with a comparable trend of increasing PSD and SVC volumes when approaching the soma (Fig. 3g,h; *P* < 0.001, Mann Whitney test).

We next investigated the E/I type of these PNs. In mammals, Glu synapses are typically asymmetrical and characterized by thick PSDs, whereas GABA synapses are symmetrical with thin PSDs^57^. However, an analysis of additional EM datasets from Glu and GABA neurons with mitochondria-localized APEX2 labeling showed no significant difference in PSD thickness or volume between the two types of synapses, with only minor exceptions in rare cases (Extended Data Fig. 10a-d), suggesting that synapses in zebrafish may not follow the criterion observed in mammals. Then we inferred the neuron type by comparing the morphology of the PNs with a reference dataset of LM-based fully-reconstructed neurons from the zebrafish mesoscopic atlas^12^. This dataset comprised 12,219 Glu, 5,371 GABA, and 2,193 Gly neurons. We pre-registered the EM query neurons to the mesoscopic brain template via a CLEM pipeline and extracted mesoscopically traced neurons within a 25-μm radius sphere centered at each query neuron’s centroid. Skeleton matching against these references yielded penalty scores, with lower scores indicating higher morphological similarity (see Methods). Manual verification of top-ranked matches enabled the classification of all 467 EM query neurons as matched or unmatched. Among them, 389 queries were successfully matched to morphologically similar E or I reference neurons (Fig. 3i and Extended Data Fig. 10e,f), while 78 showed morphological discrepancies from their top references (Extended Data Fig. 10g), suggesting the need for more complete reconstructions and broader reference coverage. Matched queries had lower penalty scores than unmatched ones (Extended Data Fig. 10h; *P* < 0.0001, Mann Whitney test), supporting the efficacy of morphology-based matching. Of the 389 matched queries, 56% (219) were E/I-specific match, with top references predominantly of a single type, and the remaining 170 were E/I-ambiguous match, showing top matches of mixed types. For instance, query neuron #41’s top references included Glu, GABA, and Gly neurons, preventing reliable E/I typing by morphology match (Extended Data Fig. 10i). Its identity as a Gly neuron was, however, confirmed by mitochondrial APEX2 labeling (Extended Data Fig. 10j), again demonstrating the value of APEX2 labeling for definitive identification of neuron types.

Based on the morphology matching, we performed E/I inference for PNs of all five APEX2-labeled LC-NE neurons (Fig. 3j,k and Extended Data Fig. 10i-l). For the LC-NE neuron #3, we inferred the E/I identity for 188 of the 374 PNs, with Glu being the predominant type (Fig. 3j; Sensory-related, 60 Glu, 15 GABA, 7 Gly; Motor-related, 75 Glu, 18 GABA, 13 Gly). Spatial analysis along the left-right dendritic axis revealed that the spatial distribution of synaptic inputs showed both E/I type- and sensory/motor modality-dependency. In comparison with I-type inputs, E-type inputs relatively preferred to proximal dendritic segments, in particular for sensory-related inputs (Fig. 3l,m; *P* < 0.05 for sensory-related inputs, *P* = 0.1072 for motor-related inputs, two-sample Kolmogorov-Smirnov test). I-type sensory-related inputs were distributed more distally than I-type motor-related inputs (*P* < 0.05, two-sample Kolmogorov-Smirnov test), while E-type motor-related inputs were biased more proximally than E-type sensory-related inputs (*P* < 0.001, two-sample Kolmogorov-Smirnov test) (Fig. 3l,m).

These results together show that LC-NE neurons receive a broad and diverse range of synaptic inputs, organized by sensory/motor modality, E/I identity, and synaptic strength at their dendrites (Supplementary Video 7).

## Intra- and inter-system common synaptic inputs among monoaminergic systems

Among the 374 PNs of the LC-NE neuron #3, 69 of them also targeted at least one other neuron among the five APEX2-labeled LC-NE neurons (Fig. 4a), thereby serving as common presynaptic neurons (CPNs) within the LC-NE system (“intra-CPNs”). These intra-CPNs were widely distributed across sensory- and motor-related regions (Fig. 4a,b), and the majority of identified intra-CPNs were E neurons (Fig. 4c). Moreover, these intra-CPNs displayed diverse innervation patterns, with the majority synapsing onto two LC-NE neurons (Fig. 4d). To assess whether intra-CPNs co-innervate LC-NE neurons randomly or selectively, we performed theoretical simulations using identical cell numbers and innervation distributions. The simulation revealed that the LC-NE neuron #2 was co-innervated with #3 more frequently than with other LC-NE neurons (Fig. 4e; *P* < 0.05, chi-square goodness-of-fit test), suggesting the potential existence of LC-NE subpopulations with preferential co-innervation. These intra-CPNs may represent as a structural substrate for synchronous activity within LC-NE neuron populations, as reported previously^9,58,59^, and help coordinate functional outputs of the LC system.

**Fig. 4.**
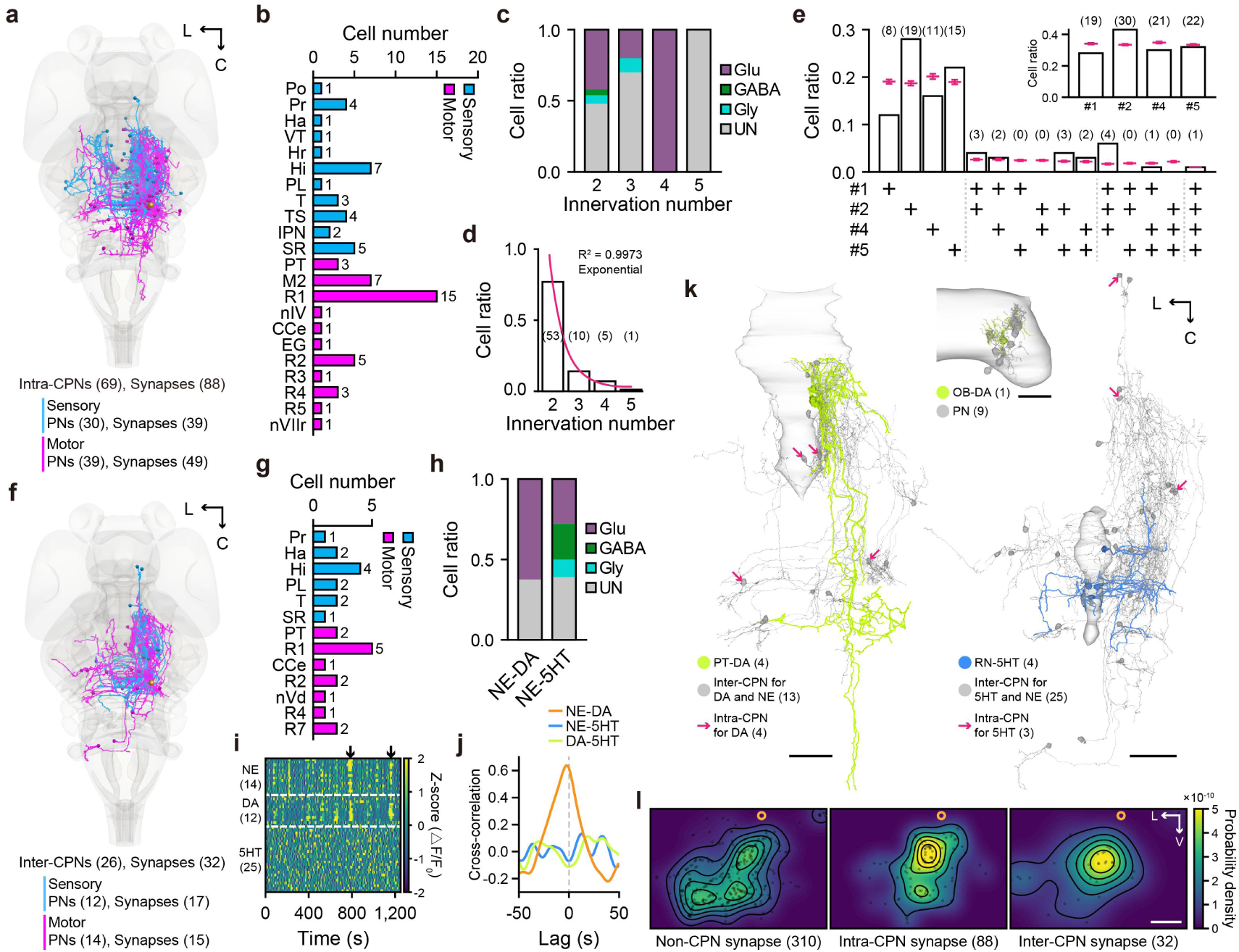
Intra- and inter-system common synaptic inputs among monoaminergic systems. **a,** Rendering of all 69 EM-reconstructed intra-LC system common presynaptic neurons (“intra-CPNs”), which formed 88 synapses on the LC-NE neuron #3. Sensory-related (blue), 30 neurons and 39 synapses; Motor-related (magenta), 39 neurons and 49 synapses. **b,** Regional distribution of 69 intra-CPNs. Neuron numbers are shown at the right of each bar. **c,** E/I composition ratios for intra-CPNs with various innervation numbers. **d,** Cell ratio of intra-CPNs with various innervation numbers. PN numbers are shown in parentheses. Exponential fitting: Y = 0.0255 + 28.09 × e^-1.816 × X^, R^2^ = 0.9973. **e,** Cell ratio of intra-CPNs categorized by innervation combination and target LC-NE identity (inset). Neuron numbers are shown in parentheses. All intra-CPNs analyzed innervated the LC-NE neuron #3. The red symbols represent theoretical cell ratios obtained from 100 iterations of random combinations, using identical cell numbers and matched innervation number distributions. Chi-square goodness-of-fit test was performed (E, χ^2^(3) = 7.8727, *P* < 0.05). Error bars represent mean ± SEM. **f,** Rendering of all 26 EM-reconstructed inter-monoaminergic system common presynaptic neurons (“inter-CPNs”), which formed 32 synapses on the LC-NE neuron #3. Sensory-related (blue), 12 neurons and 17 synapses; Motor-related (magenta), 14 neurons and 15 synapses. **g,** Regional distribution of 26 inter-CPNs. Neuron numbers are shown at the right of each bar. **h,** E/I composition ratios for inter-CPNs with different monoaminergic-type combinations. **i,j,** Analysis of synchrony in spontaneous activities among three monoaminergic neuron types. Heatmap in (**i**) depicts spontaneous calcium activities (Z-scored) in multiple monoaminergic neurons, and the cross-correlation curves between each pair are shown in (**j**). *Ki(th:GAL4-VP16);Ki(tph2:GAL4FF,cmlc2:EGFP);Tg(UAS:GCaMP6s)* larvae at 6 dpf were used for *in vivo* confocal calcium imaging. Black arrows indicate two events of spontaneous activities synchronized between LC-NE and PT-DA neurons. **k,** Rendering of representative DA (green) and 5HT (blue) neurons with their partially reconstructed PNs (gray). Only synapses successfully traced back to their PN somata are shown. PNs of the OB-DA neuron (top-middle) were obtained by dedicated tracing, while PNs of the PT-DA (left) and RN-5HT (right) neurons were identified common inputs (inter-CPNs) discovered during the reconstruction of the five LC-NE neurons’ inputs. Notably, some PNs (red arrows) also innervate multiple PT-DA or RN-5HT neurons, serving as intra-CPNs for these systems. The counts of neurons are shown in parentheses. **l,** Distribution of non-CPN (left), intra-CPN (middle), and inter-CPN (right) inputs on the dendrites of the LC-NE neuron #3. Orange circles represent the soma. The number in parentheses indicate the number of synapses analyzed. Both intra-CPN and inter-CPN inputs favor proximal distributions, whereas non-CPN inputs display a more dispersive distribution. Left-right axis: *P*(non-CPN *vs* intra-CPN) < 0.0001, *P*(non-CPN *vs* inter-CPN) = 0.5716, *P*(intra-CPN *vs* inter-CPN) < 0.05. Dorsal-ventral axis: *P*(non-CPN *vs* intra-CPN) < 0.001, *P*(non-CPN *vs* inter-CPN) < 0.01, *P*(intra-CPN *vs* inter-CPN) = 0.3284. The two-sample Kolmogorov-Smirnov test was used. C, caudal; L, left; V, ventral. Scale bars, 20 µm (**l**) and 50 µm (**k**).

In contrast, 305 of the 374 PNs (81.28%) reconstructed for the LC-NE neuron #3 did not target any of the other four LC-NE neurons and accounted for 330 of 418 (78.95%) synapses onto this neuron. We also observed that the two reconstructed Hcrt neurons exclusively innervated the LC-NE neuron #4 (see Extended Data Fig. 10k). Together, these findings indicate that individual LC-NE neurons receive both unique and shared synaptic inputs, aligning with recent studies supporting the heterogeneous and modular organization of the LC-NE system^8,9,60–63^.

We then extended the analysis to other neuromodulatory systems. Among the 374 PNs of the LC-NE neuron #3, we identified 26 widely distributed CPNs that co-innervated PT-DA or RN-5HT neurons (“inter-CPNs”; Fig. 4f,g). Morphology-based assessment of E/I type indicated that inter-CPNs connecting LC-NE and PT-DA neurons were predominantly Glu (5/8), with the remainders being E/I-ambiguous (3/8) (Fig. 4h). Meanwhile, inter-CPNs linking LC-NE and RN-5HT neurons showed a balanced E/I characteristics (Glu, 5/18; GABA, 4/18; Gly, 2/18; E/I-ambiguous, 7/18) (Fig. 4h). We thus hypothesized that LC-NE and PT-DA neurons may receive shared feedforward driving signals, whereas LC-NE and RN-5HT neurons may be subject to common inhibitory control. To support this hypothesis, we performed *in vivo* calcium imaging across these monoaminergic nuclei in the same larval zebrafish, and observed that LC-NE neurons exhibited spontaneous activities synchronized with PT-DA neurons, but not with RN-5HT neurons (Fig. 4i,j). These inter-CPNs could provide a structural basis for coordinated activity across distinct monoaminergic systems, enabling more diversified regulatory functions. Some inter-CPNs also innervated multiple PT-DA or RN-5HT neurons, serving as intra-CPNs for these monoaminergic systems (Fig. 4k, left and right). Similar to the LC-NE system, PNs targeting PT-DA and RN-5HT neurons were also widely-distributed, which contrasts with the spatially restricted input profiles of OB-DA neurons (Fig. 4k, top-middle), suggesting a conserved organizational principle across monoaminergic systems. Notably, synaptic inputs from intra- and inter-CPNs preferentially targeted proximal dendritic regions, whereas those from non-CPN sources showed more dispersed innervation patterns (Fig. 4l).

Taken together, these results offer a multi-level view on the spatial organization of broad-yet-sparse synaptic inputs onto LC-NE neurons’ dendrites, characterized by: 1) broad-yet-sparse synaptic inputs; 2) proximity-dependent synaptic strength scaling; 3) modality and E/I type-based distribution; 4) dual input architecture (private *vs.* shared); 5) Cross-system shared inputs with proximal bias (Fig. 5 and Supplementary Video 7).

**Fig. 5.**
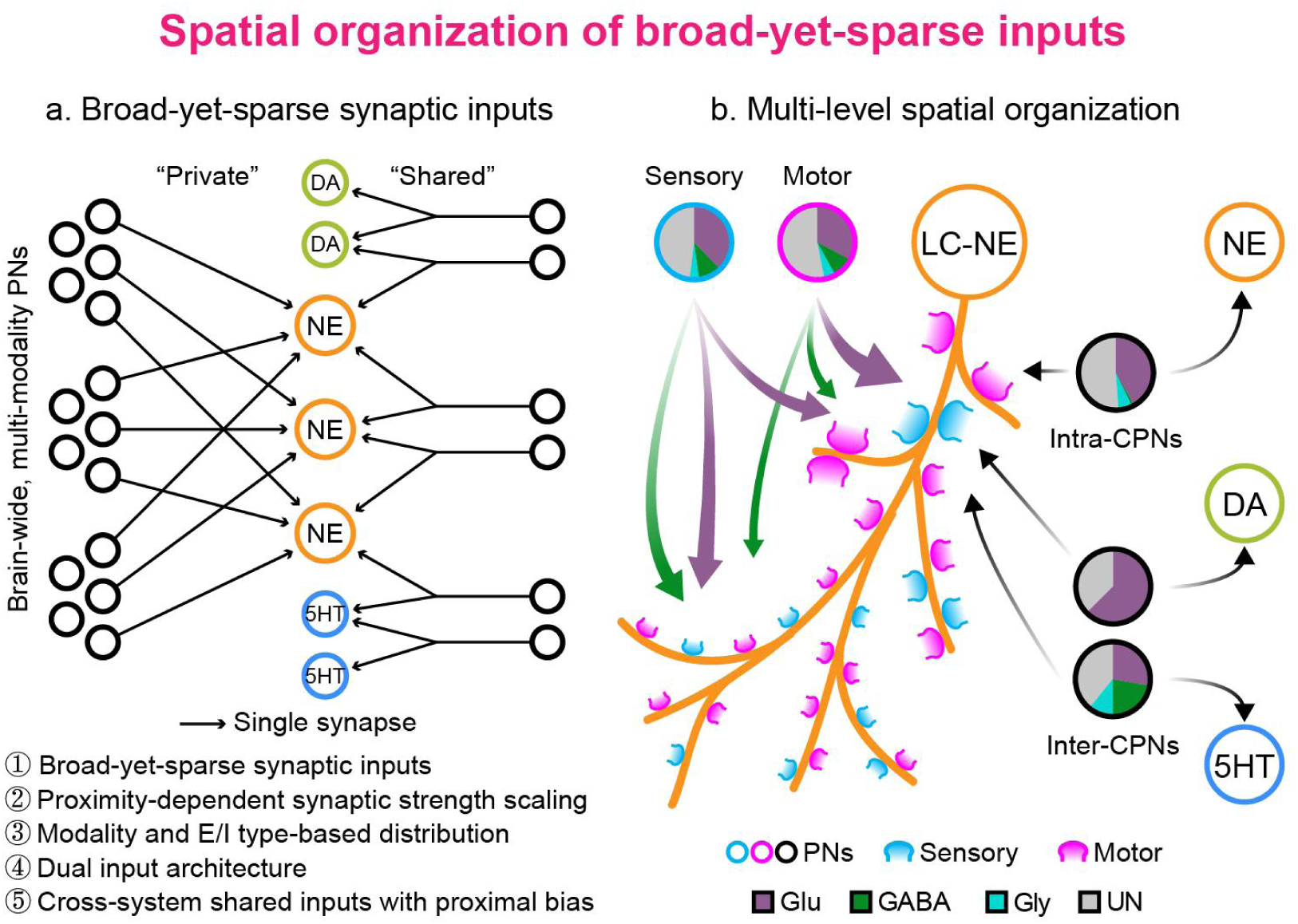
Multi-level spatial organization of broad-yet-sparse synaptic inputs to individual LC-NE neurons. Individual LC-NE neurons receive broad-yet-sparse synaptic inputs (**a**), which are spatially organized by sensory/motor modality, E/I identity, synaptic strength, and private/shared types (**b**).

## Discussion

Fish-X represents the first successful implementation of multiplexed APEX2 labeling within an intact vertebrate brain, supported by a suite of genetic tools including relevant stable zebrafish lines and plasmid constructs. The APEX2-based GETEM strategy provides distinct advantages: (i) registration-free neuron identification, (ii) multiplexed annotation via orthogonal compartmental labeling that bypasses the spectral limits of fluorescent proteins^64,65^, and (iii) reliable labeling of long-range projecting axons in partial EM reconstructions where full neuronal morphologies are unavailable^66^. In comparison, the traditional CLEM approach offers an alternative by transferring neuronal identities from fluorescently labeled LM data into EM volumes via image registration^37,41,42,44,45^. By leveraging existing transgenic lines and utilizing an H_2_O_2_-free sample preparation, this strategy is generally compatible with high-quality EM preservation. Importantly, the two strategies can be synergistically combined—the fluorescent signals from APEX2-fused proteins precisely match their DAB products, which serve as intrinsic fiducial landmarks that streamline and enhance the accuracy of CLEM registration. Furthermore, by implementing the CLEM framework beyond a single specimen^67^, our mesoscopic atlas-based strategy operates as a computational CLEM variant that matches EM neuronal morphologies to a mesoscopic reference atlas. Collectively, these complementary strategies constitute a versatile toolkit for comprehensive molecular annotation of zebrafish EM connectomes.

Using this whole-brain, molecularly annotated EM dataset encompassing multiple neuromodulatory types, we investigated the fine-scale spatial organization of synaptic inputs to neuromodulatory systems through the lens of the LC-NE system, an evolutionarily conserved hub for brain-wide regulation. Our analysis reveals a structural blueprint for global integration in the LC-NE system, which receives broad, sparse inputs from sensory-, sensory relay-, and motor-related regions across the brain. With individual PNs contributing very few synapses—on average, fewer than two—this “broad-yet-sparse” connectivity pattern is ideally suited for monitoring global brain state through consensus^68^. Individual LC-NE neurons therefore detect when widespread, coincident activity across brain regions reaches a threshold signaling a salient event, rather than responding to a single, localized signal. Within this architecture, dominant proximal excitatory inputs from motor-related regions create a low-threshold channel for motor commands or feedback, ensuring rapid and reliable NE release to facilitate swift behavioral state transitions^69–71^. In contrast, distal sensory inputs, particularly inhibitory ones, likely exert modulatory control by setting global neuronal excitability through dendritic calcium spike modulation or shunting inhibition, rather than directly vetoing action potentials^72–75^. This fine-tuning enables context-dependent NE neuromodulation. The contrast between this organizational logic and that of cortical pyramidal neurons^11,76–81^ underscores the fundamental diversity in dendritic integration mechanisms across cell types and highlights the necessity of building neuron models grounded in cell-type-specific connectomes.

At the population level, accumulating molecular, morphological, and functional evidence indicates that the LC-NE system exhibits modular rather than homogeneous organization^8,9,60–63^. Our study provides new insights into this organization from the perspective of synaptic input architecture. We found that individual LC-NE neurons receive both “private” (i.e., non-CPNs) and shared (i.e., intra-CPNs) inputs, collectively forming an architectural basis for population-level activity. Private inputs likely drive sparse, asynchronous NE release from single neurons. In contrast, shared inputs with varying innervation numbers can recruit specific subsets or even the entire population of LC-NE neurons. This recruitment, potentially further synchronized via gap junctions among LC-NE neurons, enables regionally distinct and intensity-graded NE release. An intriguing question for future investigation is whether this private–shared input dichotomy corresponds to the well-established tonic and phasic firing modes of LC-NE neurons, potentially encoding distinct presynaptic information streams^82,83^. Furthermore, by extending this modularity concept beyond the LC-NE system, we identified shared inputs (inter-CPNs) that co-innervate both LC-NE and other monoaminergic neurons, suggesting the existence of higher-order, inter-system modules that provide a structural basis for the coordinated neuromodulation observed in diverse physiological and pathological brain states.

The observed co-innervation architecture between distinct neuromodulatory systems not only demonstrates the value of our multiplexed molecularly annotated EM resource, but also underscores the need for more complete reconstructions to fully resolve the inputome of each neuromodulatory system and the organizational principles governing their interactions. Notably, the anatomical comprehensiveness of Fish-X, encompassing the retina, optic nerve, brain, and aSC, enables the systematic investigation of both neuromodulatory and sensorimotor pathways across the entire CNS. To fully realize this potential, the dataset can be computationally refined through advanced alignment and segmentation algorithms combined with online collaborative proofreading to improve reconstruction efficiency^84–88^. Furthermore, the ATUM-SEM approach has preserved all sections, enabling targeted re-imaging of both suboptimal regions and previously unmapped areas. In parallel, there is a pressing need for semi- or fully-automated reconstruction algorithms specifically designed for GETEM-labeled neurons, particularly those with challenging cytosol labeling^89^. Our dataset, with its rich APEX2-labeled resource encompassing a wide range of staining intensities, provides an ideal benchmark for developing and refining such algorithms. Beyond enhancing the current resource, this study has established a versatile APEX2-based genetic toolbox and optimized procedures—including real-time quality control, image repair, and image denoising—that enable the creation of next-generation EM datasets with expanded simultaneous neuron type annotation and higher image quality.

In summary, we have established a feasible pipeline for generating molecularly annotated whole-brain EM-reconstructions in larval zebrafish and provided a valuable resource for the community. Based on this dataset, we have revealed the principles of synaptic input architecture of the LC-NE system. Further investigation building upon this work will not only expand the ultrastructural mapping of input – output connectomes for diverse neuromodulatory neuronal populations—providing a substrate for their realistic neuronal modeling—but also deepen our understanding of how neuromodulatory neurons are synaptically integrated within the bi-pathway brain architecture.

## Methods

### Zebrafish husbandry

Adult Zebrafish (*Danio rerio)* were kept in a circulating system at 28℃ under a 14:10 hour light-dark cycle. All animal protocols were reviewed and approved by the Animal Care and Use Committee of the Center for Excellence in Brain Science and Intelligence Technology (CEBSIT), Chinese Academy of Sciences (NA-046-2023). Larvae were fed with *Brachionid* rotifers from 4.5 – 9.0 dpf, then switched to freshly hatched *Artemia* nauplii thereafter. The sex of embryonic and larval zebrafish was undetermined.

### Zebrafish lines

All transgenic zebrafish larvae were in *nacre* background. The *vmat2*, *th1*, *dbh*, *tph2*, *hcrt*, *glyt2*, *vglut2a*, *gad1b* promoters were used to label NE/DA/5HT, NE/DA, NE, 5HT, Hcrt, Gly, Glu, and GABA neurons, respectively. Published fish lines used in this study include *Tg(slc6a5:GFP)cf3* (*abbreviation*: *Tg(glyt2:GFP)*)^90^, *Tg(elavl3:Hsa.H2B-GCaMP6f)jf7*(*Tg(HuC:H2B-GCaMP6f)*)^91^, *Tg(5×UAS-hsp70l:GCaMP6s)nkUAShspzGCaMP6s13a* (*Tg(UAS:GCaMP6s)*)^92^, *Ki(th:GAL4-VP16)ion32d* (*Ki(th:GAL4-VP16)*)^93^, *Ki(tph2:EGFP)ion39d* (*Ki(tph2:EGFP)*)^93^, *Tg(slc17a6:GAL4FF)ion177d* (*Tg(vglut2a:GAL4FF)*)^94^. The unpublished fish lines *TgBAC(gad1b:GAL4-VP16)ion19h*^12^ and *gSAIzGFFM242A* (*Gt(vmat2:GAL4FF)*) were provided by Dr. Jie He and Dr. Koichi Kawakami, respectively. *Ki(tph2:GAL4FF,myl7:EGFP)ion127d* (*Ki(tph2:GAL4FF,cmlc2:EGFP)*) was constructed using a CRISPR/Cas9-based knockin method^93^. To generate the larva for Fish-X dataset, *Tg(dbh:GAL4FF;5×UAS-hsp:cytosol-APEX2-TagRFP)ion142d* (*Tg(dbh:GAL4FF;UAS:cytosol-APEX2-TagRFP)*), *Tg(hcrt:APEX2-TagRFP-NLS)ion100d* (*Tg(hcrt:APEX2-TagRFP-NLS)*), and *Tg(slc6a5:Hsa.Cox8a-APEX2-TagRFP)ion97d* (*Tg(glyt2:MITO-APEX2-TagRFP)*) were constructed using the Tol2-based transgenic system^95^. All other APEX2-based transgenic lines and plasmids were listed in Supplementary Table 1. The *APEX2* sequence was cloned from the commercial plasmid *Bact2-APEX2-GBP*^96^ (Addgene, Cat# 67668). The *dbh*, *hcrt*, and *glyt2* promoter sequences were cloned from plasmids *pTol2-dbh (short)-GAL4FF-5XUAS-hsp-TRPV1-RFPT*, *pTol2-hcrt-GAL4-VP16*, and *pTol2-glyt2-P2A-ECFP*, respectively. Signal peptide sequences were listed in Supplementary Table 2. Original PKU constructs were provided by Dr. Yu-Long Li.

### Immunostaining

Larvae were fixed overnight on a shaker at 4℃ using 4% paraformaldehyde (PFA, Sigma, Cat# 158127) prepared in phosphate-buffered saline (PBS). Following three 5-min washes with PBS containing 1% Tween-20 (PBST), brains were carefully dissected using fine forceps and subjected to another three PBST washes. Samples then underwent graded dehydration and rehydration through sequential incubation in (v/v): 25% methanol/75% PBST (5 min, RT), 50% methanol/50% PBST (5 min, RT), 75% methanol/25% PBST (5 min, RT), 100% methanol (4 h, −20℃), followed by reverse gradient to 25% methanol, then three 5-min PBST washes at RT. After ddH_2_O rinse (5 min, RT) and acetone treatment (10 min, −20℃), samples were rehydrated through ddH_2_O and three final PBST washes. Blocking was performed overnight at 4℃ with gentle shaking, followed by incubation with primary antibodies (1:200 in blocking buffer) at 4℃ for 24 – 48 h. After primary antibody recovery and three quick PBST rinses, extended washing was performed (2 × 1 h, 1 × 20 min, 1 × 10 min). Secondary antibodies (1:500 in blocking buffer) were applied at 4℃ for 36 h, followed by removal, three quick PBST washes, and final rinses (3 × 10 min, 1 × 30 min) before light-protected storage on ice for fluorescence imaging. Commercial antibodies included Anti-Orexin-A antibody (Millipore, Cat# AB3704), Anti-Tyrosine Hydroxylase antibody (Millipore, Cat# MAB318), Alexa Fluor^®^ 488 Goat anti-Rabbit secondary antibody (Abcam, Cat# AB150077), and Alexa Fluor^®^ 568 Goat anti-Mouse secondary antibody (Invitrogen, Cat# A-11004).

### *In vivo* structural and functional imaging

At 3 dpf, multiplexed APEX2-labeled larvae were initially screened using an upright fluorescence stereomicroscope. The preparations, embedded in 1.5% low-melting-point agarose (Invitrogen, Cat# 15517014), were carefully verified at 5 dpf using an upright fluorescence microscope. The labeling efficiency was evaluated to determine larvae were used for further EM sample preparation. At 6 dpf, larvae were paralyzed with 1 mg/mL α-bungarotoxin (Tocris, Cat# 2133) and dorsally embedded in 1.5% low-melting-point agarose. Morphological fluorescence imaging was performed using an upright confocal laser scanning microscope (FV3000, Olympus, Japan) equipped with a 40× water-immersion objective (NA = 0.8). Imaging parameters included a frame size of 635.9×635.9 μm² (0.621×0.621 µm^2^ per pixel), with two-field stitching along the rostrocaudal axis to cover the SC, Z-axis intervals of 1 – 1.5 µm, and 2 – 3 line averaging. Selected brain regions were further imaged at higher zoom factors to resolve fine spatial patterns of APEX2 signals.

To simultaneously record calcium dynamics in LC-NE, PT-DA, and RN-5HT neurons, we performed *in vivo* calcium imaging on 6-dpf *Ki(th:GAL4-VP16);Ki(tph2:GAL4FF,cmlc2:EGFP);Tg(UAS:GCaMP6s)* larvae, using a confocal microscope (FV3000, Olympus, Japan). Imaging parameters included a frame size of 636.4×636.4 μm² (1.243×1.243 µm^2^ per pixel), with resonant scanning mode, imaging depth of ∼200 μm, Z-axis intervals of 10 μm, volumetric imaging rate at ∼0.72 Hz, and total acquisition duration of ∼30 min per session.

### DAB staining

The optimized DAB staining protocol was adapted from published procedures^97^. The 0.1 M Tris-HCl buffer (pH 7.4) was prepared by dissolving 0.6055 g Tris-Base (Millipore, Cat# 648311) in 50 mL of ultrapure water, with pH adjustment using concentrated HCl. A 10 mg tablet of 3,3’-diaminobenzidine tetrahydrochloride (DAB·4HCl, Sigma, Cat# D5905) was subsequently dissolved in 10 mL of this buffer through vigorous vortexing under light-protected conditions. The solution was then filtered through a 0.22-μm filter (Millipore, Millex^TM^-GP 0.22-μm filter) to obtain a 1 mg/mL DAB stock solution, which was stored at 4℃ and protected from light. Samples were sequentially rinsed with 0.1 M phosphate buffer (3 × 10 min), Tris-HCl buffer (2 × 10 min), and DAB solution (2 × 10 min) on a shaker at 4℃ to remove residual aldehydes. For the reaction, 1 μL of 30% H_2_O_2_ (Sinopharm Chemical Reagent Co., Ltd., Cat# 10011218) was mixed with 1 mL DAB stock to prepare the working solution, and then samples were incubated for 2 h at RT in complete darkness. The reaction was terminated by washing with Tris-HCl buffer (2 × 10 min). The catalysis by APEX2 of DAB polymerization resulted in dark brown precipitates, which were clearly visible under bright-field stereomicroscope. Post-fixation with osmium tetroxide (OsO_4_) was conducted immediately following DAB staining to preserve the osmiophilic reaction products.

### EM sample preparation

A mixed fixative containing 2% PFA and 2% glutaraldehyde (GA) was used for effective whole-brain fixation. A 1 mL aliquot of 25% GA (Sigma, Cat# G5882, stored at −20℃) was thawed at 4℃ under light protection. Concurrently, 1 g PFA powder was dissolved in 20 mL of ultrapure water within a fume hood, with 1 – 2 drops of 1 M NaOH added. The solution was then sealed, protected from light, and heated at 65℃ with magnetic stirring until fully clarified. After cooling to RT, the solution was filtered through a 0.22 μm filter to yield 5% PFA stock. The 10-mL working fixative was prepared immediately before dissection by mixing 4 mL of 5% PFA, 0.8 mL of 25% GA, 5 mL of 0.1 M phosphate buffer (pH 7.4), and 0.2 mL of ultrapure water. Larvae were embedded in 1.5% low-melting-point agarose dissolved in extracellular solution (in mM: 134 NaCl, 2.9 KCl, 2.1 CaCl_2_·2H_2_O, 1.2 MgCl_2_·6H_2_O, 10 HEPES, 10 glucose; 290 mOsm, pH 7.8). The agarose covering dorsal epidermis and pectoral fins was removed. Using glass micropipettes, the skin was carefully dissected from the hindbrain ventricle while avoiding damage to tissues. The trunk was rapidly transected anterior to the swim bladder using ophthalmic scissors, after which the head was gently extracted with fine forceps and immediately immersed in fresh fixative. Fixed samples were sealed and stored at 4℃ in darkness overnight.

Subsequent procedures were adapted from the rOTO protocol (*98, 99*). First, a 1:1 (v/v) mixture of 0.2 M phosphate buffer and 2% OsO_4_ (TED PELLA, Cat# 18451) was prepared to obtain an OsO_4_-phosphate solution, which was then used to prepare a reduced OsO_4_-phosphate solution containing 0.03 g/mL potassium hexacyanoferrate trihydrate (Sigma, Cat# P9387). All OsO_4_-containing solutions were freshly prepared within a fume hood, tightly sealed, protected from light at 4℃, and properly collected for disposal after use. Fixed samples were washed with 0.1 M phosphate buffer at 4℃ (4 × 10 min) and then immersed in the OsO_4_-phosphate solution (200 – 500 μL per 1.5 mL tube, 2 h, 4℃). Concurrently, a 0.01 g/mL aqueous thiocarbohydrazide (TCH, Sigma, Cat# 88535) solution was placed in a 60℃ oven and gently flipped to facilitate dissolution. The TCH solution was removed from the oven and allowed to cool slightly above RT before use. The solution was then filtered through a 0.22 μm filter directly into the sample tube that had been rinsed with ultrapure water (3 × 10 min, 4℃; 1 × 10 min, RT). After a 30-min light-protected TCH treatment at RT, samples were washed with ultrapure water (4 × 10 min, 4℃). The reduced OsO_4_-phosphate solution was then added (200 – 500 μL per 1.5 mL tube, 30 min, RT). Afterward, samples were washed with ultrapure water (4 × 10 min, 4℃). Meanwhile, a 2% aqueous uranyl acetate solution (EMS, Cat# 22400) was centrifuged at 13,000 rpm, and the supernatant was carefully transferred into the sample tube (200 μL per 1.5 mL tube). Samples were then sealed and stored at 4℃ overnight, protected from light.

On the next day, the uranyl acetate waste solution was collected for proper disposal, followed by washing samples with ultrapure water (3 × 10 min). Subsequently, a graded dehydration series was performed on ice using ethanol solutions (30%, 50%, 70%, 80%, 90%, 95%, and 100% v/v, each for 10 min) followed by anhydrous acetone (2 × 10 min). On the day of fixation, the resin volume was calculated, and the premix was prepared by sequentially pipetting 10 mL of Pon-812 epoxy resin monomer (SPI, Cat# 90529-77-4), 3.36 mL of dodecenyl succinic anhydride (DDSA, SPI, Cat# 26544-38-7), and 7.04 mL of nadic methyl anhydride (NMA, SPI, Cat# 25134-21-8) into a pre-dried tube at RT to form a three-component premix. This was then sealed and mixed on a rotary mixer for at least 3 hours. Resin infiltration was conducted immediately following dehydration. 400 μL DMP-30 epoxy accelerator (SPI, Cat# 90-72-2) was added to 20 mL premix, followed by 30-min rotary mixing to prepare the final resin solution. The resin solution was either used immediately after a 30-min bubble-settling period or kept mixing to prevent polymerization at RT. Samples underwent graded acetone-resin infiltration at ratios of 3:1, 1:1, and 1:3 (v/v) for 8 h, 8h, and overnight, respectively. They were then transferred to resin-filled vials, subject to negative pressure using a 5 mL syringe, and placed on a horizontal shaker for resin infiltration.

This process was conducted over two days with resin changes occurring twice daily at intervals of 8 – 12 h. All procedures were performed under controlled environmental conditions (temperatures < 25℃; humidity < 30%). Following the two-day resin infiltration, samples were transferred to pre-dried 21-well flat molds and oriented using sterile toothpicks. The molds were then placed in a drying box containing silica gel beads and polymerized at 37℃ for 8 – 12 h. After verifying and minimally adjusting sample orientation at night, the temperature was first increased to 45℃ overnight, then to 60℃ the following morning. Fully polymerized blocks were removed from the oven after 48 h and stored in a dry cabinet.

### X-ray tomography

X-ray tomography was conducted to rapidly and non-destructively evaluate the quality of EM samples using the Xradia 520 Versa system (Zeiss, Germany) at the Instrumental Analysis Center of Shanghai Jiao Tong University. Samples exhibiting uniform staining, minimal deformation, and no apparent mechanical damage were selected for further ultrathin sectioning and SEM imaging. Samples were scanned at 60 kV with 1 μm^3^ voxel resolution (∼2 h per sample). Data processing was conducted using Dragonfly (Object Research Systems, Canada). Sequential images were imported with resampling parameters and voxel resolution specified. Sample orientation was interactively manipulating reference axes in the display window. Selected views were exported as projection sequences via the “*Derive new from current view*” function. 3D animations were created using the *Video marker* tool. The video denoising was performed using an open-source Real-ESRGAN-General-x4v3 model.

### Whole-brain ultrathin section collection

Using an ultramicrotome equipped with an automated tape-collecting system (ATUMtome, RMC, USA), 33-nm serial sections were obtained^40^. Prior to collection, trimmed excess blank resin around the larva using a double-edged razor blade. The front guide slot of the ATUMtome was properly positioned, and parameters including sectioning speed (0.8 mm/s), tape-advancing speed (0.6 mm/s), and water refilling interval (5-sec duration every 8 min) were set. The anti-static device and intermittent water replenishment system were activated to ensure sequential and stable section collection on hydrophilic, conductive polyimide film strips (KAPTON-PET, DuPont, USA). If section instability was observed during collection (e.g., incomplete sections), immediately halted sectioning and either repositioned the knife edge or replaced the diamond knife (UltraMaxi, Diatome, Switzerland). Throughout the process, ambient conditions were maintained at 25℃ and 60% humidity. Following collection, section-bearing strips were segmented and mounted onto 4-inch monocrystalline silicon wafers using 0.8-mm-wide conductive adhesive tape (SPI, USA), with careful pressing to eliminate air bubbles. The mounted wafer was placed into a LEICA EM ACE 600 high-vacuum coater for carbon deposition (6 – 10 nm film thickness). The section surfaces were cleaned using nitrogen gas blowing, and subsequently stored in a dry cabinet at 25℃ and 30% humidity. The entire collection process, which spent three days and utilized three diamond knives, yielded an overall damage rate of 6.56%. Sections were numbered #2594 – #25480 in a caudal-to-rostral order. The damaged 1,503 sections with folds, tears, and severe contamination were algorithmically repaired to ensure that intact portions were registered to adjacent sections^100^. The majority of defects did not occur at identical locations across adjacent sections, thus collectively posing no substantial impact on proofreading.

### SEM image acquisition

To enhance sample stability and backscattered electron (BSE) signal contrast, wafers were pre-irradiated using a custom-built irradiation system (9 mA, 2 kV, 10 s continuous exposure). Post-irradiation wafers were transferred to an optical microscope (Clippers FBT, China) for the acquisition of navigation images via line-scan imaging with 7.6 μm pixel resolution. Image acquisition was conducted simultaneously across three SEMs (Navigator NeuroSEM100, FBT, China), each equipped with a custom-built collection software designed to maximize efficiency in image output. The pixel resolution was set to 4 nm, the dwell time to 43.5 ns, and the acceleration voltage to 2 kV. Auto focus and astigmatic correction were configured to 80 s per section. To expedite acquisition efficiency and imaging success rate, a region-specific focusing protocol coupled with photography was implemented. Each section was divided into a grid of ∼54 tiles (arranged in 9 rows × 6 columns) at 12,000×12,000 pixels per tile. The adjacent tiles were acquired with a 12% overlapping area. 16 tiles (4×4 arrangement) constituted one standard region. Edge regions containing fewer than 16 tiles were processed equivalently. Regions were sequentially numbered and navigated in a serpentine pattern. Prior to high-resolution scanning, a low-resolution focus was conducted to ensure the algorithm efficacy in pinpointing the optimal focal range at the center of the section, which was set at a resolution of 50 nm with a dwell time of 80 ns. The stage was then oriented to the first region’s center for high-resolution focusing and astigmatism correction with a resolution of 4 nm and a dwell time of 80 ns, followed by snake-pattern tile acquisition. After capturing all tiles within a region, the stage advanced to the next region (2.5 s/tile for positional adjustment and data transfer). This cycle iterated until full-section coverage was achieved, maintaining serpentine navigation between regions. The process was repeated automatically for subsequent sections until wafer-wide imaging completion.

To enhance the accuracy and robustness of large-scale EM image processing, we introduced an image quality assessment and tissue validity detection module prior to algorithmic analysis. A 2D U-Net model was trained to evaluate image sharpness and assigned a quality score to each section. Low-quality images were then reviewed using a custom annotation tool, allowing experts to determine whether re-acquisition or replacement was necessary. Besides, we implemented CNN-based tissue mask generation. A U-Net model was trained on three manually annotated chunks (8,600×8,600×31 voxels at 32×32×33 nm^3^ per voxel) to classify voxels into brain tissues, backgrounds, and defect regions (e.g., folds, tears). All predicted masks were manually reviewed. The quality scores and tissue masks were applied throughout the processing pipeline for artifact exclusion and region-of-interest selection, supporting segmentation and agglomeration.

### Image stitching and alignment

The 3D volume registration of EM images was accomplished through tile stitching and section alignment. The stitching workflow proceeded through three computational stages: 1) feature detection and matching using Scale-Invariant Feature Transform (SIFT) descriptors^101^ extracted from tile overlap areas; 2) pairwise displacement estimation via coordinate transformation analysis; 3) global position optimization through linear least squares minimization. Specifically, SIFT-matched feature point pairs served as the basis for calculating local translation vectors between tiles. To mitigate error accumulation, a global optimization framework simultaneously adjusted all tile positions by minimizing the total displacement error across the entire section.

A multi-scale alignment method was implemented to register serial sections, first through coarse-scale processing followed by fine-scale refinement. To address error accumulation, the pipeline employed a sub-volume-division strategy, partitioning the original dataset into 764 spatially coherent stacks (each containing 31 consecutive sections). Continuity across stacks was established by selecting the first section of each stack as a reference section (764 total). These references underwent affine transformation to correct notable deformations while maintaining spatial invariance throughout alignment, serving as stable pivots to prevent cumulative error propagation. Alignment then proceeded within each stack, from coarse to fine scales. Sections were downsampled 32-fold during coarse-scale alignment. Correspondences between two adjacent sections were initially extracted using SIFT descriptors and subsequently filtered using the RANSAC algorithm^102^. Affine transformation was applied to align adjacent sections. Sections were downsampled 8-fold during fine-scale alignment. Damaged sections were first aligned to their neighbors using an expected affine method^91^. Subsequently, an optimized elastic method^103^ generated dense correspondence fields between adjacent sections, guiding the iterative refinement of deformation meshes that balanced local adaptability with global smoothness. Finally, Thin-Plate Spline (TPS) transformations^104^ extrapolated these deformations to original-resolution sections, preserving ultrastructural continuity at synaptic resolution.

### Bidirectional CLEM

To match the resolution of the brain template of the Zebrafish Mesoscopic Atlas (1 μm^3^ per voxel), the 764 reference sections mentioned above were further downsampled in the xy-plane to generate downsampled whole-brain EM images (992×992×990 nm³ per voxel). Multimodal image registration was accomplished through a three-stage approach. First, whole-brain nuclei were extracted from both the Zebrafish Mesoscopic Atlas and the ssEM dataset to create nuclei point clouds, which were then rigidly aligned using the Iterative Closest Point (ICP) algorithm^105^. Manually-annotated nuclei from four 6-dpf *Tg(HuC:H2B-GCaMP6f)* larvae were used to generate the average point cloud for the LM brain template, while nuclei from the ssEM dataset were automatically segmented. Second, the EM images were globally constrained using X-ray tomography data, to correct the distortion along the rostral-caudal axis (“banana effect”). Third, bidirectional deformation fields between LM and EM spaces were computed through 3D template matching and TPS transformations. Soma regions across the entire brain, along with other EM-identifiable regions including the pineal gland, pituitary gland, and bilateral optic tectal neuropils, were manually annotated on downsampled whole-brain EM images and registered with their counterparts in the brain template of the Zebrafish Mesoscopic Atlas, serving as reference landmarks for joint optimization.

### Chunk-based 3D reconstruction

Whole-brain 3D reconstruction was performed through an automated pipeline consisting of: (1) image enhancement and denoising, (2) local registration, (3) dense segmentation, (4) intra-chunk agglomeration, and (5) inter-chunk stitching. To efficiently process large-scale computational tasks, we implemented a distributed computing strategy based on spatial block partitioning. Specifically, the original 3D volume was divided into a series of overlapping chunks to ensure boundary continuity and consistency in prediction. These chunks were then preprocessed and spatially realigned, and subsequently treated as independent inference tasks scheduled for parallel execution. The entire computational workflow was deployed on a high-performance distributed parallel computing architecture managed by the Slurm (Simple Linux Utility for Resource Management) workload manager. All sub-tasks were automatically submitted via batch scripts to a Slurm-managed HPC (High-Performance Computing Cluster) consisting of more than 30 nodes. Based on the availability of computational resources, the scheduler dynamically distributed tasks across multiple physical nodes. Each node was equipped with a heterogeneous configuration of GPUs, including NVIDIA^®^ A100, V100, and GeForce RTX 4090 models, thereby maximizing utilization of available hardware accelerators. Upon completion, the outputs of all sub-tasks were automatically aggregated and post-processed to reconstruct the full prediction results over the entire volume. For tasks including the segmentation of neurons, synapses, and mitochondria, the block and overlap sizes were set to 8,600×8,600×31 and 600×600×1 voxels, with a resolution of 4×4×33 nm^3^ per voxel. For tasks including the segmentation of nuclei and vessels, the block and overlap sizes were set to 1,075×1,075×31 and 75×75×1 voxels, with a resolution of 32×32×33 nm^3^ per voxel.

### Preprocessing and dense segmentation

To address the Poisson and Gaussian noise inherent in EM images and to overcome the challenges of self-supervised denoising with degraded inputs, we implemented Blind2Unblind for implicit lossless self-supervised denoising^106^. This framework leveraged a mask-guided medium to enhance and denoise EM images. It consisted of a globally-aware mask mapper driven by mask-based sampling and a non-blind lossless training strategy based on re-visible loss. Next, to enhance local 3D continuity and improve inference performance, we adopted a spatial registration strategy inspired by the preprocessing pipeline of the Flood-Filling Network (FFN)^107^. For each chunk mentioned above, a 4×4 affine transformation matrix was estimated to align it with a global reference space. Neural network inference, aggregation, and merging were then performed on the aligned chunks. After dense segmentation, inverse transformations were applied to map the results back to the original coordinate system, followed by cropping to restore the original dimensions. All affine matrices were precomputed and cached prior to processing to reduce computational overhead.

After preprocessing, we employed a boundary detection-based dense segmentation approach to process the whole-brain EM dataset. The image segmentation network adopted a U-shaped architecture with residual blocks, and was trained to predict nearest-neighbor affinity maps along all three spatial dimensions. The training dataset consisted of seven randomly selected volumes, each comprising 30 consecutive sections with a spatial size of 2,000×2,000×30 voxels at a resolution of 4×4×33 nm^3^. All these volumes were manually annotated by experts. A composite loss function combining weighted cross-entropy and Dice loss was employed to enhance segmentation performance, particularly in the presence of class imbalance. To improve model generalization, we applied a range of data augmentation techniques during training, including basic transformations such as rotation, flipping, and elastic deformation, as well as robustness-oriented strategies such as CutBlur, CutNoise, and Mixup. In addition, we simulated common imaging artifacts including registration shifts, motion blur, and partial slice dropouts, to enhance the model’s capability to handle real-world imperfections, thereby improving the stability and accuracy of the segmentation.

### Intra-chunk agglomeration

To construct the signature graph, we first performed over-segmentation of the membrane probability map using a 2D watershed algorithm based on distance transform. Specifically, small connected membrane boundary components were removed through thresholding. Given the strong anisotropy of the data (with lateral resolution of 4 nm per voxel and axial resolution of 33 nm per voxel), the Euclidean distance transform was applied to each membrane probability map. The maximum distance value was then used as the seed point to execute the watershed algorithm, producing 2D watershed superpixels. These superpixels were subsequently merged to achieve the 3D reconstruction. The image could be represented using watershed superpixels, where the superpixels served as vertices and the physical connectivity between them formed the edges of a graph model. The superpixel merging problem was thus formulated as a vertex partitioning problem on the graph. In this study, the problem is modeled as the minimum cost multicut problem (MCMP) on the superpixel adjacency graph^108^. For feature extraction, three types of features were manually extracted: boundary features between superpixels, edge features within each superpixel, and inter-layer features derived from the anisotropic dataset. A feature vector of length 625 was designed for each edge between two superpixels. A random forest classifier was then trained to predict the probability of merging each pair of superpixels. Based on maximum likelihood estimation, the predicted probability is converted into the cost for merging the two superpixels. This resulted in an instance of the multicut problem. Additionally, previously reconstructed and expert-verified whole-brain nucleus and synapse segmentation results were introduced as soft priors^109^. Superpixels belonging to the same nucleus should be merged preferentially, while superpixels on opposite sides of the same synapse should be merged less likely. At the same time, the aggregation range was confined to the effective tissue mask area to prevent the merging of defect regions with other superpixels, thereby avoiding erroneous propagation. Given that the combinatorial optimization problem defined here is NP-hard, and exact methods are not polynomial-time solvable, this study framed the problem as a sequential decision-making task for node aggregation on the graph. The problem was solved optimally using the heuristic greedy additive edge contraction (GAEC) framework^110^. Finally, the results were mapped back to the original over-segmented image obtained from the watershed distance transform.

### Inter-chunk stitching

Following the preliminary reconstruction of all individual chunks, we developed an overlap-aware fragment agglomeration algorithm to identify and merge neuronal segments across adjacent chunks, thereby enabling the reconstruction of globally continuous neuronal structures. This algorithm integrated a deep learning-based candidate identification module with a set of rule-based merging constraints. Specifically, we trained a 3D-ResNet-based neural network classifier to evaluate fragment pairs extracted from sliding 3D windows within overlapping regions, producing a probability score indicating whether the pair originated from the same cell. Candidate merges were then further filtered based on multiple criteria, including intersection-over-union (IoU) thresholds, voxel overlap chunk, and the presence of defect masks, to ensure that the final agglomeration decisions maintained both structural continuity and biological plausibility. By combining large-scale parallel processing enabled by the Slurm workload manager mentioned above with this customized post-processing pipeline, we established an efficient workflow for dense reconstruction.

### Segmentation of nuclei and vessels

We annotated nuclei and vessels in every fifth section of the downsampled ssEM dataset with a resolution of 32×32×165 nm^3^, generating high-quality training sets with dimensions of 9,000×6,000×764 voxels. These results were used to train a 2D segmentation network. We employed nnU-Net^111^ for training and inference, with a patch size set to 1,024×1,024×4 and a batch size of 2. To maintain training stability, we chose stochastic gradient descent (SGD) with a momentum of 0.99 as the optimizer and applied a “poly” learning rate policy, starting with a learning rate of 0.01 and a maximum of 1,000 epochs. Upon completion of inference, we restored the results to the original space, resulting in a coarse reconstruction of the entire volume. Next, we implemented a sliding window strategy along the z-axis, systematically traversing connected instances within each window, to achieve a refined reconstruction of the whole-brain nuclei and vessels. The window length was set to 30, with an overlap of 4. When the number of voxels within the receptive field was less than 1000, the z-axis voxel range was under 2, or the instance did not contact adjacent windows, we classified it as reconstruction noise and removed it. Approximately 35% of nuclei and all vessels underwent further manual proofreading, followed by quantitative analysis and 3D visualization.

### Nucleus clustering

We developed a feature extraction pipeline, integrating both traditional handcrafted morphological features and high-level representations learned through self-supervised learning, to analyze whole-brain nuclei with diverse morphologies. Manually-proofread 176,789 nuclei (nuclei with incomplete structures in rostralmost and caudalmost sections were excluded) were processed using the CloudVolume framework to compute 3D surface meshes and then extracted 13 traditional morphological features using the Trimesh library. These 13 features collectively characterized nucleus compactness (*Φ*, *κ_A_*, *κ_V_*), surface irregularity (Δ*A*, Δ*A_b_*), shape elongation (*ρ_max_*, *R_range_*), and spatial dispersion (*r_max_*, *r_min_*). The following notations were adopted:

*M*: 3D triangular mesh representing a nucleus surface.

*V*: Volume of the nucleus mesh (μm³).

*A*: Surface area of the nucleus mesh (μm²).

*C*: Geometric centroid of the nucleus mesh.

*M_h_*: Convex hull mesh of *M*.

*V_h_*, *A_h_*: Volume and surface area of convex hull, respectively.

*S_b_*: Minimal bounding sphere of *M*.

*V_b_*, *A_b_*: Volume and surface area of bounding sphere, respectively.

*v_i_*: *i*-th vertex of mesh *M*.

||·||: Euclidean norm in *ℝ*^3^.

*Φ*: Sphericity defined as *Φ* = π^1/3^(6*V*)^2/3^/*A*, where *Φ* ∈ [0, 1] and *Φ* = 1 defines a perfect sphere.

*κ_A_*: Surface convexity defined as *κ_A_* = *A*/*A_h_*

*κ_V_*: Volume convexity defined as *κ_V_* = *V/V_h_*

Δ*A*: Surface concavity defined as Δ*A* = *A* - *A_h_*

Δ*V*: Volume concavity defined as Δ*V* = *V_h_* - *V*

*ρ_max_*: Elongation ratio defined as *ρ_max_* = max*_i_*(||*v_i_* - *C*||)/min*_i_*(||*v_i_* - *C*||)

*R_range_*: Radial range defined as *R_range_* = max*_i_*(||*v_i_* - *C*||) - min*_i_*(||*v_i_* - *C*||)

*r_max_*: Normalized max radius defined as *r_max_* = (max*_i_*(||*v_i_* - *C*||))^3^/*V*

*r_min_*: Normalized min radius defined as *r_min_* = (min*_i_*(||*v_i_* - *C*||))^3^/*V*

*σ_A_*: Spherical area ratio defined as *σ_A_* = *A*/*A_b_*

*σ_V_*: Spherical volume ratio defined as *σ_V_* = *V*/*V_b_*

Δ*A_b_*: Spherical surface excess defined as Δ*A_b_* = *A_b_* - *A*

Δ*V_b_*: Spherical volume excess defined as Δ*V_b_* = *V_b_* - *V*

*e* : Eccentricity computed based on principal component analysis of the mesh vertices as 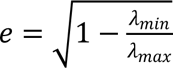, where *λ_max_* and *λ_min_* are the largest and smallest eigenvalues of the mesh vertex covariance matrix, respectively.

To complement these interpretable handcrafted features, a self-supervised deep learning framework was implemented to extract high-level geometric representations of the nucleus shape. Unlike voxel- or mesh-based formats, point clouds provide a compact and resolution-independent representation that preserved fine morphological details. Each nucleus surface mesh *M* was uniformly sampled using Poisson disk sampling in Trimesh to obtain a point cloud *P* ∈ *ℝ^n×^*^3^, where *n* = 3,072 represented the number of sampled vertices. The sampling process ensured uniform spatial distribution across the mesh surface while preserving geometric details. The point clouds were processed using a Siamese self-supervised architecture based on SimSiam^112^ to learn transformation-invariant descriptors. During training, each point cloud *P* was augmented into two distorted views *P_1_* and *P_2_* using random rotations, Gaussian noise, and jittering. The augmented views were processed through a three-stage neural network pipeline:

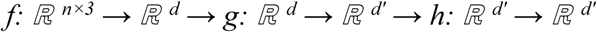

where *f*(·) (Encoder network, PointNet-based backbone) mapped point clouds to *d* = 1,024-dimensional embeddings, *g*(·) (Projector network, 2-layer MLP) mapped to *d’* = 256-dimensional projections, and *h*(·) (Predictor network, 2-layer MLP) with same dimensionality. The model was trained to minimize a symmetric negative cosine similarity loss:

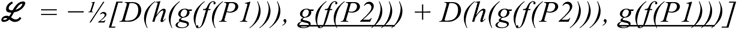

where *D(x, y) = x·y/(||x||_2_ ||y||_2_)* denoted the cosine similarity function. The underlined terms were treated as constants during backpropagation and did not contribute gradients. This objective encouraged the network to learn rotation- and noise-invariant representations while preventing representational collapse. Each nucleus was ultimately represented by a *d* = 1,024-dimensional embedding vector *z* = *f(P)* extracted from the trained encoder. To reduce redundancy and enable integration with handcrafted features, we applied Uniform Manifold Approximation and Projection (UMAP)^113^ with the following hyperparameters (Target dimensionality: *d_UMAP_* = 8; Number of neighbors: *k* = 15; Minimum distance: *d_min_* = 0.1; Distance metric: Euclidean). This yielded an 8-dimensional deep feature representation *z_UMAP_ ∈ ℝ^8^* for each nucleus. The final feature representation for each nucleus was constructed by concatenating the 8-dimensional deep feature embedding *z_UMAP_ ∈ ℝ^8^* with the 13-dimensional handcrafted morphological feature vector *f_morph_ ∈ ℝ*^13^: *f_hybrid_* = *[f_morph_; z_UMAP_] ∈ ℝ*^21^, where [·; ·] denoted vector concatenation. Prior to concatenation, all features were standardized using z-score normalization: *f’ = (f − μ)/σ*, where *μ* and *σ* represented the mean and standard deviation computed across the entire dataset, respectively. This hybrid descriptor *f_hybrid_* leveraged both interpretable morphological traits (providing biological insight) and high-capacity deep embeddings (capturing complex geometric patterns), enabling robust and scalable analysis of nucleus morphology.

Similarly, we adopted the same methodology for morphological analysis of somata. Since somata could be characterized with the aid of associated structures such as nuclei and mitochondria, the computation of traditional somatic features incorporated not only the aforementioned morphological descriptors but also integrated nucleus- and mitochondria-based features as auxiliary representations.

### Segmentation of PSDs, SVCs, and mitochondria

A three-class 3D U-Net network was trained to predict PSDs and SVCs at the resolution of 8×8×33 nm^3^ per voxel. We annotated four regions (∼12×12×1 μm^3^ per volume), generating training sets encompassing 146M voxels. We optimized the network using the multi-class cross-entropy loss and SGD optimizer. During training, a warm-up learning rate strategy was employed with an initial learning rate of 0.01, a linear warm-up period of 2k steps, and cosine decay over a total of 500k training steps. Data augmentation was performed to improve the network’s robustness, including random rotation, random flipping, grayscale transformation, elastic transformation, and blur simulation. To test the network performance, we manually annotated another three regions as test sets (∼8×8×1 μm^3^ per volume). The evaluation of these regions yielded an average precision of 85.60 ± 3.21% and an average recall of 88.37 ± 1.59%.

A 3D U-Net network was trained to detect the mitochondria at the resolution of 8×8×33 nm^3^ per voxel. We annotated five regions (∼12×12×1 μm^3^ per volume), generating training sets encompassing 467M voxels. To separate adjacent mitochondria, we designed a network to simultaneously predict both a mitochondrial mask (trained with binary cross-entropy loss) and an instance boundary mask (optimized using a combined binary cross-entropy and Dice loss). Training proceeded for 500k iterations with SGD optimizer, applying a learning rate scheduling strategy identical to the synapse detection task. Data augmentation protocols were consistently implemented as in the synapse detection framework. For automated identification of glycinergic neurons, we annotated additional 284 soma-localized mitochondria labeled with APEX2. Neurons were classified as glycinergic if their somata contained ≥ 4 labeled mitochondria.

### Proofreading

We conducted extensive manual and semi-automated proofreading of neurons, PSDs, SVs, mitochondria, vessels, and brain regions using a custom software Advanced Connectome Engine (ACE) that integrated the Hadoop- and HBase-based data management, computing modules, and collaborative tools. We used this software to coordinate proofreading tasks, facilitate collaborative proofreading workflows, and log all user operations. Current and immediately preceding proofreading results were compared to evaluate proofreaders’ performance. The proofreading team consisted of 30 – 40 full-time proofreaders from commercial companies and several experts from the laboratories involved. Reconstruction results were represented as graph structures that connected segmentation contours across all serial sections belonging to a single supervoxel. The contours of adjacent layers were compared to identify anomalies and guide proofreaders to target regions requiring correction.

To aid the proofreading workflow, we utilized a series of foundation models for segmentation^114–116^, including the Segment Anything Model (SAM), SAM 2 and SAEM². Based on these models, we implemented two additional strategies to enhance ACE’s human-computer interaction. First, the SAM ViT-H model checkpoint was exported to an ONNX (Open Neural Network Exchange) model with additional quantization and optimization. ONNX Runtime was used to execute these models on CPU/GPU, which supported cross-platform inference and improved software runtime. Second, the SAM embeddings generated by encoders were precomputed and saved in CloudVolume using kempressed format, which demonstrated higher compression ratios compared to raw and fpzip encodings. For 2D semi-automated segmentation, we utilized manually traced neuronal skeletons with extra positive/negative points as input prompts for the SAM model. However, the pretrained SAM failed to accurately segment densely packed, homogeneous neuronal processes^117^. To address this, we proposed the finetuned SAEM² with auxiliary tasks to acquire complementary multi-head information for neuron segmentation in the ssEM dataset. The auxiliary task involved identifying cell boundaries (plasma membranes) to complement the original SAM that primarily focused on cell masks. For 3D segmentation, we employed the SAM 2 augmented with two additional strategies: (1) We trained the system to predict cross-correlation maps and subsequently computed the shifts between adjacent 2D EM images. (2) We utilized the distance transformation maps from the previous cell masks to sample prompt points for generating the next cell masks.

### Micro-mesoscopic morphology comparison

We inferred the neuronal types of APEX2-unlabeled neurons based on morphology comparison between the ssEM dataset and a zebrafish mesoscopic atlas^12^. A preprocessed reference file was generated, containing the x-y-z coordinates of all E and I neurons and their mirrored counterparts from the Zebrafish Mesoscopic Atlas (n_Glu_ = 12,219, n_GABA_ = 5,371, n_Gly_ = 2,193). We first calculated neuronal skeletons (stored as swc files) from manually-corrected EM supervoxels. These skeletons were then transformed into the common physical space of the mesoscopic brain template using bidirectional CLEM approach mentioned above. Using the soma center of each EM query neuron as the origin, we retrieved all single-neuron morphological data within a 25 μm spherical radius based on the preprocessed reference file. For each EM query neuron, we performed pairwise morphology comparisons with the retrieved mesoscopic reference neurons. Cross-scale skeleton matching established morphological correspondences between the same type of neurons across imaging modalities by quantifying spatial relationships between their reconstructed skeletons. The matching similarity was constructed by calculating the unidirectional distance (*D*) from EM skeletons (*S_E_*) to LM skeletons (*S_L_*) as follows,

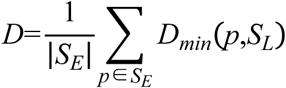

where *D_min_* represents the shortest distance from *S_L_* to the point *p* in *S_E_*. The distance for each point in *S_E_* was calculated, and the average was then determined. The *S_L_* that minimized *D* was chosen as the most similar skeleton corresponding to *S_E_*. This distance-based scoring system ranked similarity (lower scores indicated closer matches), with the top-ranked matches (swc filenames and scores) exported. These results were manually verified using a custom software Brain Atlas Viewer (BrAVe)^13^.

### 3D visualization

In accordance with open-science principles, we employed the Neuroglancer precomputed format and CloudVolume interface for data management and 3D visualization through both Neuroglancer and webKnossos^84^. For data encoding, we used JPEG compression for ssEM images and compressed_segmentation format for region silhouettes and segmentation masks. We further implemented the Igneous^118^ for multiprocessed block-wise downsampling, skeletonization and meshing with the following agglomeration. For skeletonization, we used the Kimimaro implementation of the automated TEASAR algorithm^119^. We then merged the skeleton splits that might appear based on nearest Euclidean neighbors, calculated the path length of the skeleton, and determined the radius of each vertex in the skeleton. For meshing, we used the automated block-wise Zmesh to generate multi-resolution meshes. These skeletons and meshes were stored in sharded formats.

We employed mesh representations to visualize brain regions and reconstructions in Neuroglancer, while utilizing annotation points to demonstrate synaptic locations. The Neuroglancer *matrix* function served as an affine transformation tool for manipulating (translate/flip/rotate/zoom) the original image layers. We used the parallel projection during Neuroglancer rendering to generate 3D scale bars. Furthermore, we optimized two existing workflows for video creation: (1) Neuroglancer workflow using Python scripts from the *video_tool* package, and (2) the open-source computer graphics software Blender, with Python scripts and JSON descriptions from neuVid. Since we acquired high-resolution images only for the retina, optic nerve, brain and aSC, we processed a pre-acquired low-resolution dataset (400×400×∼1,650 nm^3^ per voxel) of the complete zebrafish head to provide comprehensive 3D rendering context used in Fig.1b and Supplementary Video 1. We first verified the section number corresponding to the whole-brain high-resolution dataset, and then generated uniformly spaced, downsampled images (992×992×660 nm^3^ per voxel) along the rostral-caudal axis using linear interpolation. These images were subsequently registered to the high-resolution dataset, ensuring alignment with multiple segmentation and reconstruction results within the same physical space. Finally, grayscale histogram equalization was performed. The ssEM dataset are accessible through the following dedicated links:

https://ng.zebrafish.digital-brain.cn/Fish-X/Brain_EM_images

https://ng.zebrafish.digital-brain.cn/Fish-X/Retina_EM_images

https://ng.zebrafish.digital-brain.cn/Fish-X/Brain_regions

https://ng.zebrafish.digital-brain.cn/Fish-X/Auto_supervoxels

https://ng.zebrafish.digital-brain.cn/Fish-X/Manual_supervoxels

https://ng.zebrafish.digital-brain.cn/Fish-X/Nuclei

https://ng.zebrafish.digital-brain.cn/Fish-X/Vessel

### Statistical analysis

No randomization and blinding were used during data collection and analysis. The statistical analysis was performed using GraphPad Prism. Error bars represent mean ± SEM. Kolmogorov-Smirnov test was used for testing normality. Kruskal-Wallis test was used in Fig. 1k (left panel, PSD/SVC/Mito), 1l (left panel), 2j (among LC-NE, PT-DA, and RN-5HT). Unpaired *t* test was used in Fig. 2j (between LC-NE/PT-DA/RN-5HT and all other types). Two-tailed Mann Whitney *U* test was used in Figs. 1k (right panel, PSD/SVC/Mito), 1l (right panel), 2f,g, 3d-h, and Extended Data Fig. 10d,h. Two-tailed two-sample Kolmogorov-Smirnov test was used in Figs. 3l,m, 4l. Linear regression analysis was used in Figs. 1j, 2l, 3g,h, and Extended Data Fig. 4h-j. Exponential fitting was performed in Fig. 4d. Chi-square goodness-of-fit test was performed in Fig. 4e. Statistical significance is defined as *P* < 0.05. The specific statistical tests, *P*-value ranges, and sample sizes are provided in the corresponding figures and/or figure legends.

### Reporting summary

Further information on research design is available in the Nature Port-folio Reporting Summary linked to this article.

## Data availability

All raw EM images are openly accessible for visualization at https://ng.zebrafish.digital-brain.cn/Fish-X/Brain_EM_images; additional image data and reconstructed results can also be explored as detailed in the Methods. Further information and requests for materials and additional details will be fulfilled by the corresponding author.

## Code availability

All original codes and algorithms are available at https://github.com/MiRA-Han-Lab/Fish-X.

## Acknowledgements

We are grateful to Drs. Yu-Long Li and Rong-Bo Sun for sharing PKU constructs, Jie He, Misha Ahrens, Koichi Kawakami, and Joe Fetcho for sharing fish lines, and Yun-Feng Hua, Jun Zhang, Xiao-Hong Xu, Ke Zhang, Yu Mu, Dan-Qian Liu, Kai Yang, and Sheng-Xiong Wang for insightful advice. We would like to thank Xu-Long Wen, and other staffs from Grand View (Beijing) Technology Co., Ltd, and Zhi Zeng, Le Wang, and other staffs from Chengdu Huizhong Tianzhi Tech. Co., Ltd. (www.hztzai.com) for proofreading. We would like to thank Kui Wang for contribution to plasmid construction, Yu Kong, Li-Jun Pan, Xu Wang, Xue Xu from Electron Microscopy Core Facility (CEBSIT) for assistance in sample preparation, Ming-Quan Chen and Hong-Li Wan from Facility of Mapping Brain-Wide Mesoscale Connectome (FMBWMC, CEBSIT) for assistance in utilizing the zebrafish mesoscopic atlas, Ting-Ting Zhao and Yong-Wei Zhong from FMBWMC for quality assessment of automated segmentation results, Jia-Wen Huang, Liu-Qin Qian, Chen-Xi Jin, and other staffs from Big Data of Brain Atlas Facility (CEBSIT) for establishing a web-sharing platform, Qian Hu from Optical Imaging Core Facility (CEBSIT) for optical imaging and image processing support, Jia Li for suggestion on fish line generation and cell type identification, Chun-Feng Shang, Rong-Wei Zhang, Yu-Fan Wang, Sha Li, Le Sun, Qiu-Sui Deng, Li-Jun Chen for discussions, Ji-Wen Bu and Ye Hua for lab management, Yan-Hua Zhu from Shanghai Jiao Tong University for supporting X-ray tomography. We thank our institute (CEBSIT) for providing access to GeminiSEM 300 (Zeiss, Germany), Talos L120C TEM (Thermo Scientific, USA), JEM-1230 TEM (JEOL, Japan), FV3000 confocal laser scanning microscope (Olympus, Japan), and high-performance computing capabilities at the Center for Data and Computing in Brain Science. We thank the Transdisciplinary Platform of Functional Connectome and Brain-Inspired Intelligence in Huairou Science City in Beijing and the Microscopic Technology & Analysis Center at the Institute of Automation, Chinese Academy of Sciences for providing technical support and device resources. This work was supported by Brain Science and Brain-like Intelligence Technology - National Science and Technology Major Project (2021ZD0204500, 2021ZD0204502 and 2021ZD0204503), Creative Research Groups (32321003) and general grant (32171461) of the National Natural Science Foundation of China, Shanghai Municipal Science and Technology Major Project (18JC1410100, 2018SHZDZX05, 25511102500), and Key Research Program of Frontier Sciences (QYZDYSSW-SMC028) and Strategic Priority Research Program (XDB32010200) of Chinese Academy of Sciences.

## Author contributions

Conceptualization and Supervision: JLD, HH, FNL

Methodology - fish line: FNL, CS, RKT, HYH

Methodology - EM sample preparation: FNL, CS

Methodology - sectioning: LLL, XHD

Methodology - imaging: LNZ, HTM, LML

Methodology - registration: YNL, XC, FXZ, TX, BHC, HRC, SC

Methodology - automated reconstruction: JZL, JL, JYG, QWX

Methodology - proofreading: FNL, CS, ZHF, YSW, SRJ

Methodology - software: JBY, LJS, XHZ

Methodology - visualization: HZ

Methodology - mesoscopic atlas: XFD

Methodology - others: YCZ, HYY, JLZ, CYQ

Analysis - LC-NE system: FNL, CS, CMZ, PSX, YXY

Analysis - others: FNL, JZL, CS, JBY, JL, YNL, HZ, YCZ, YQ, YCG

Writing - original draft (methods): JZL, CS, JBY, JL, YNL, YCZ, HYY, HZ, LLL, LNZ, XHD

Writing - original draft (all sections): FNL, JLD

Writing - review & editing: FNL, JLD

## Extended Data Figures and Legends

**Extended Data Fig. 1.**
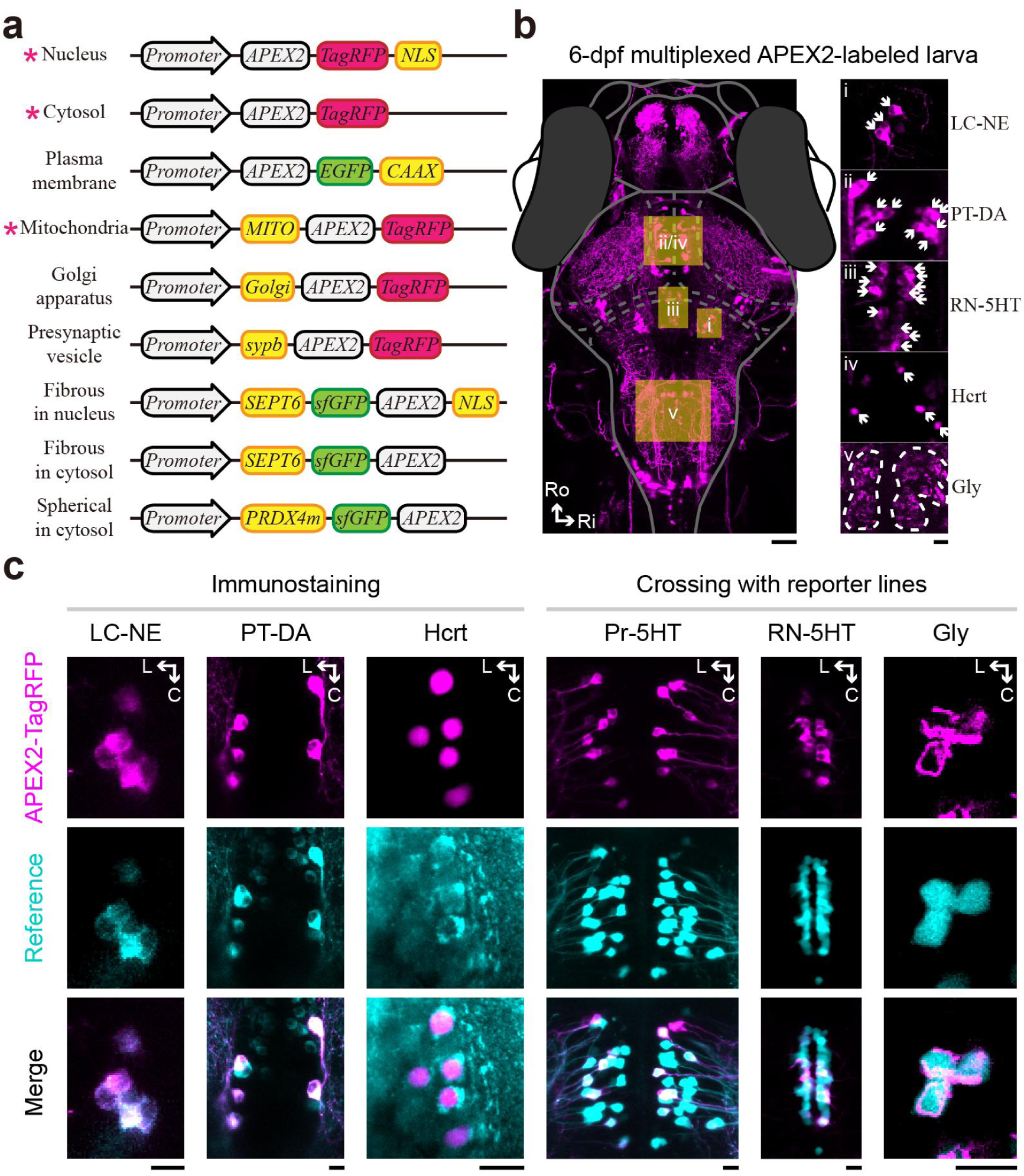
Versatile APEX2 genetic toolbox for multiplexed labeling. **a,** Schematic of genetic constructs for the nine APEX2 labeling patterns. Red asterisks mark the three transgenic lines employed for the production of larvae used in the Fish-X collection. Yellow boxes indicate corresponding signal peptides or PKU tags. **b,** Z-projected whole-brain (left) and expanded (right) fluorescent images of the 6-dpf multiplexed APEX2-labeled larva used for whole-brain ssEM in this study. White arrows indicate cytosol- or nucleus-localized APEX2-conjugated TagRFP signals in LC-NE (i), PT-DA (ii), RN-5HT (iii), and Hcrt (iv) neurons. Mitochondria-localized signals are outlined by white dashed lines in hindbrain Gly neurons (v). **c,** Specificity of APEX2 labeling in 6-dpf larvae was validated by immunohistochemical staining (left) and crossing with neuronal type-specific reporter lines (right). Representative fluorescent images show APEX2-TagRFP (top), immunofluorescence/reporter GFP (middle), and merged channels (bottom). *Gt(vmat2:GAL4FF);Tg(dbh:GAL4FF;UAS:cytosol-APEX2-TagRFP)* larvae were immunostained with an anti-tyrosine hydroxylase antibody to validate NE/DA neurons, and were crossed with *Ki(tph2:EGFP)* larvae to validate 5HT neurons. *Tg(hcrt:APEX2-TagRFP-NLS)* larvae were immunostained with an anti-hypocretin antibody to validate Hcrt neurons. *Tg(glyt2:MITO-APEX2-TagRFP)* larvae were crossed with *Tg(glyt2:GFP)* larvae to validate Gly neurons. Ro, rostral; C, caudal; L, left; Ri, right. LC, locus coeruleus; MO, medulla oblongata; OB, olfactory bulb; PT, posterior tuberculum; Hc, caudal hypothalamus; Pr, pretectum; RN, raphe nucleus; NE, noradrenergic; DA, dopaminergic; 5HT, serotonergic; Hcrt, hypocretinergic; Gly, glycinergic. Scale bars, 10 µm (**b**, right; **c**) and 50 µm (**b**, left).

**Extended Data Fig. 2.**
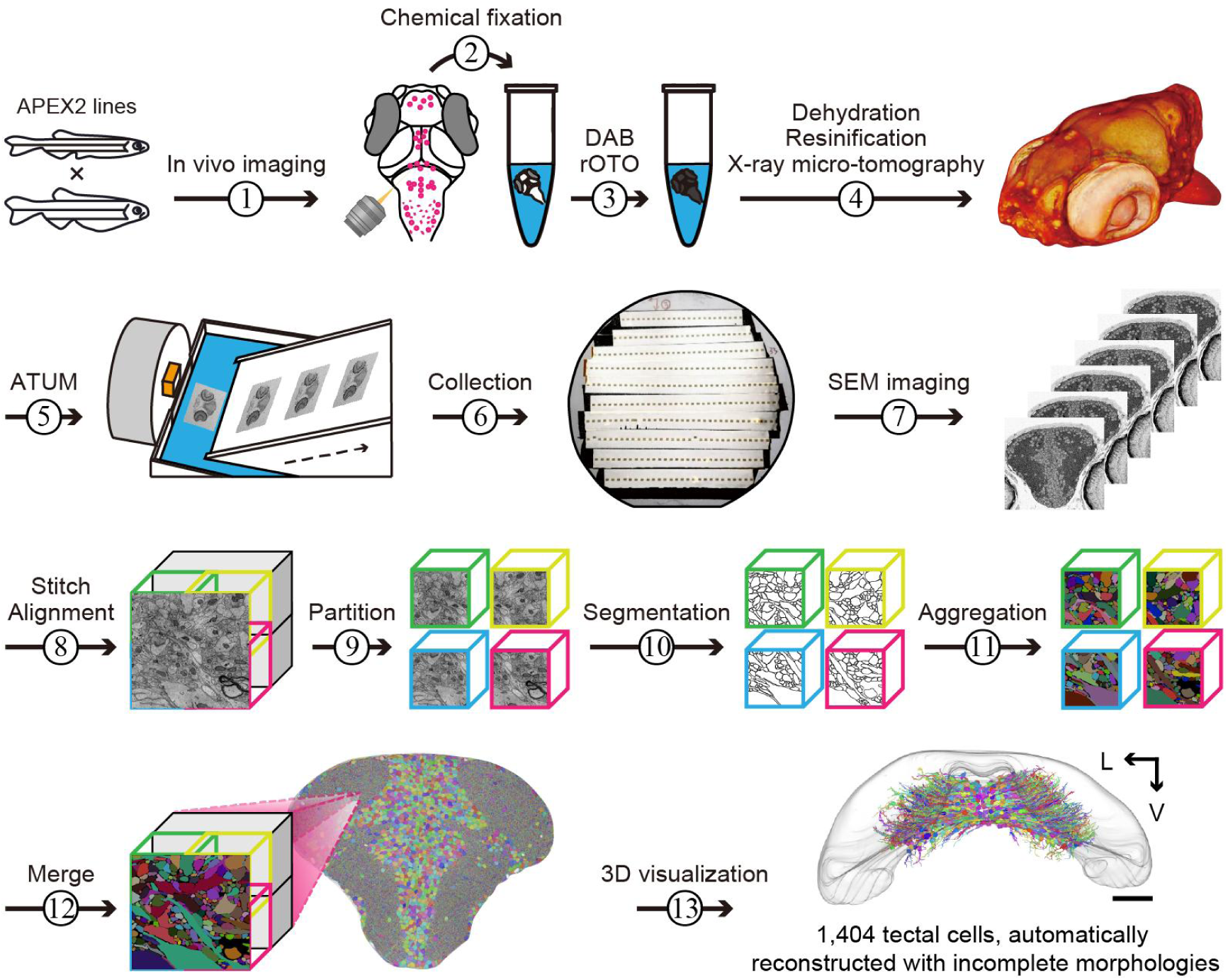
Integrated workflow for whole-brain ssEM of larval zebrafish. This workflow outlines the key steps for generating the Fish-X dataset. We began by using *in vivo* confocal fluorescence imaging to assess the coverage of labeled nuclei and the number of labeled neurons in a larva featuring simultaneous labeling of NE, DA, 5HT, Hcrt, and Gly neurons. The sample was then chemically fixed overnight. The following day, it was treated with a DAB-H_2_O_2_ solution, and the labeling performance was initially evaluated under a bright-field stereoscope. Subsequently, the sample underwent rOTO staining, dehydration, and resinification. We employed X-ray micro-tomography to rapidly and non-destructively verify the sample’s integrity, with the resulting signal serving as a second round of labeling assessment. Using the ATUM system, the sample was sectioned along the rostrocaudal axis to create a library of 33-nm sections, which were collected on 49 wafers. Image acquisition was then carried out over 6 months using three SEMs. Following this, the acquired images were stitched and aligned. The resulting whole-brain volume was partitioned into 34.4×34.4×0.99 μm^3^ blocks for distributed segmentation, aggregation, and merging, culminating in a fully automated 3D reconstruction (see Methods). The magnified rendering of 1,404 automatically reconstructed optic tectal periventricular cells with incomplete morphologies is shown (bottom right). Scale bars, 50 µm.

**Extended Data Fig. 3.**
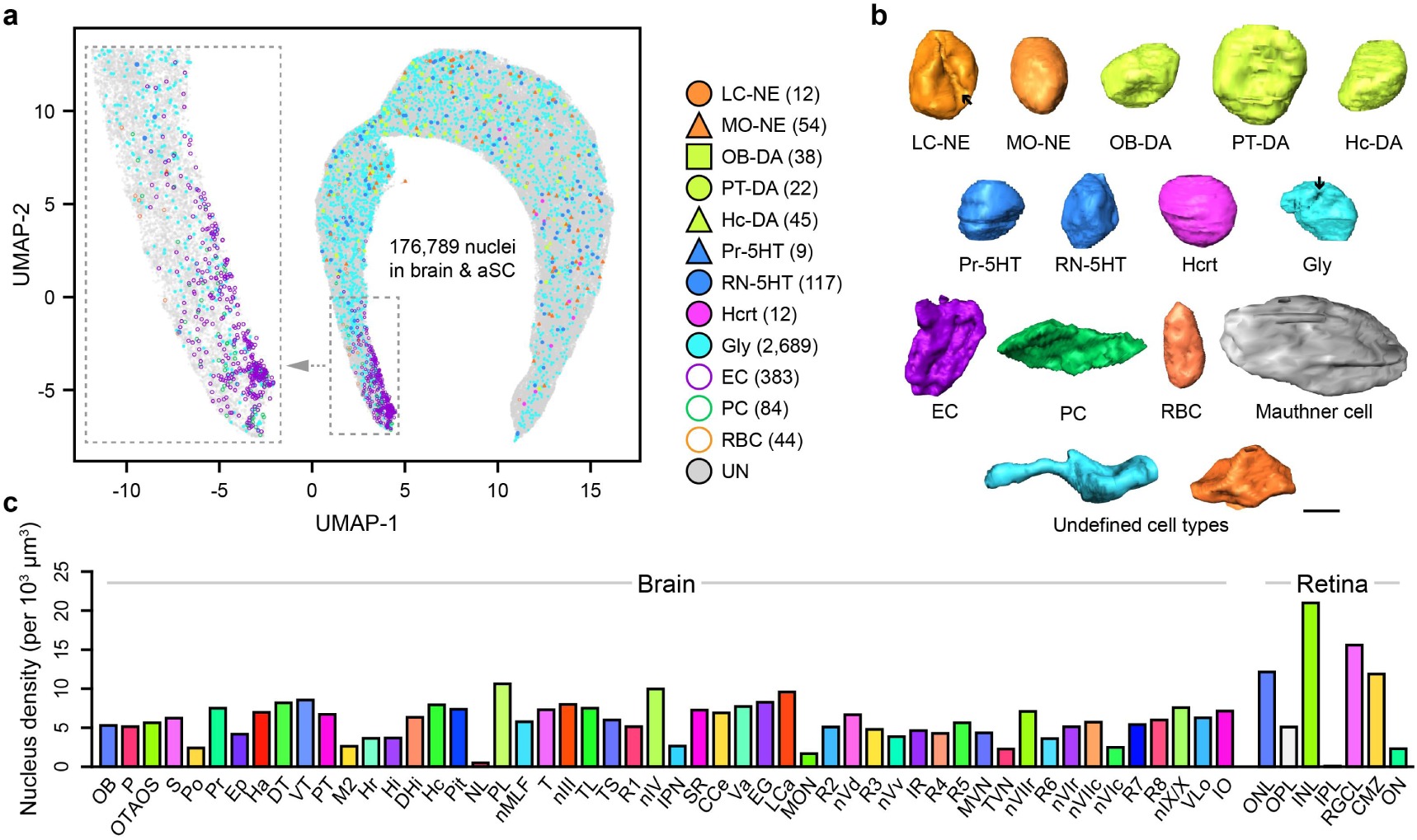
Brain-wide heterogeneity of nuclear morphology and density. **a,** UMAP visualization of whole-brain nucleus clustering (176,789 cells) using combined 8-dimensional deep-learned and 13 traditional features (see Methods). All nuclei were manually proofread after automated segmentation, resulting in optimal 3D morphological reconstruction. 21 nuclei located at dataset boundaries were excluded from clustering due to incomplete morphologies. Based on APEX2-labeling, 2,689 Gly neurons were automatically identified, and 309 neurons of neuromodulatory types were manually annotated by experts. The number of neuromodulatory-type neurons are shown in parentheses. In addition, 511 vasculature-associated cells, including endothelial cells (ECs, *n* = 383), pericytes (PCs, *n* = 84), and red blood cells (RBCs, *n* = 44), were manually annotated based on their location and perisomatic features. Inset (left): vasculature-associated cells tending to aggregate. Among all identified cells, we classified 70,114 as putative neurons based on the presence of, at least one synapse in their automatically reconstructed supervoxels. The remaining cells were presumed to be glial cells or neurons with poor automated process reconstruction. **b,** Rendering of nuclei from diverse cell types. Black arrows indicate shallow grooves and infoldings on the nuclei of LC-NE and Gly neurons. Two examples of unclassified neurons with irregular nuclear morphologies are also shown. **c,** Distribution of nucleus density distribution across 52 brain regions and 7 retinal regions. Color code corresponds to brain regions as in Fig. 1i. Scale bars, 3 µm (**b**).

**Extended Data Fig. 4.**
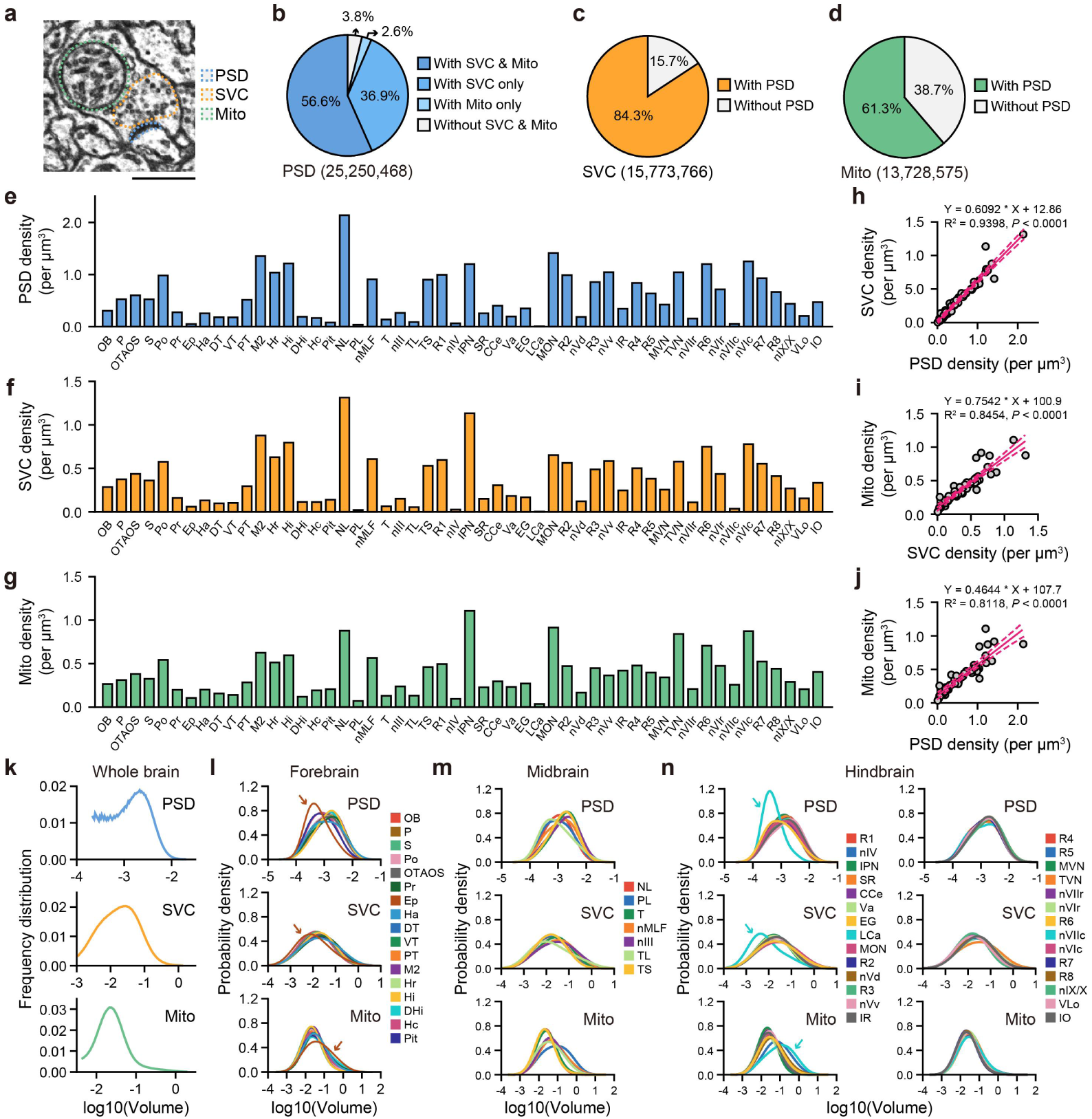
Density and volume distributions of PSDs, SVCs, and Mito. **a,** A representative EM image (4×4 nm^2^ per pixel) of a synapse, with the PSD (blue), SVC (orange), and a mitochondrion (green) outlined. **b-d,** Summary of spatial colocalization among PSDs (**b**), SVCs (**c**), and Mito (**d**). Automated 3D reconstructed supervoxels were used to identify whether these structures were co-localized within the same presynaptic compartments. **e-g,** Density distributions of PSDs (**e**), SVCs (**f**), and Mito (**g**) across 52 brain regions. **h-j,** Pairwise linear regression of densities among PSDs, SVCs, and Mito. The solid red lines represent the fitted regression line, and the dashed red lines indicate the 95% confidence interval of the fit. Each circle represents individual brain regions. SVC *vs* PSD: Y = 0.6092 × X + 12.86, R^2^ = 0.9398, *P* < 0.0001 (**h**); Mito *vs* SVC: Y = 0.7542 × X + 100.9, R^2^ = 0.8454, *P* < 0.0001 (**i**); Mito *vs* PSD: Y = 0.4644 × X + 107.7, R^2^ = 0.8118, *P* < 0.0001 (**j**). **k,** Lognormal frequency distributions of volumes among PSDs (top), SVCs (middle), and Mito (bottom) across brain regions. The volume was calculated using the automated 3D reconstructed supervoxels, and the data from all 52 regions were pooled together. **l-n,** Lognormal probability density distributions of volumes among PSDs (top), SVCs (middle), and Mito (bottom) from the forebrain (**l**), midbrain (**m**), and hindbrain (**n**). All regions exhibit comparable lognormal distributions, with Ep (**l**, brown) and LCa (**n**, cyan) showing slight differences from the majority, as indicated by arrows. Regions are randomly color-coded. PSD, postsynaptic density; SVC, synaptic vesicle cloud; Mito, mitochondria. Scale bars, 500 nm (**a**).

**Extended Data Fig. 5.**
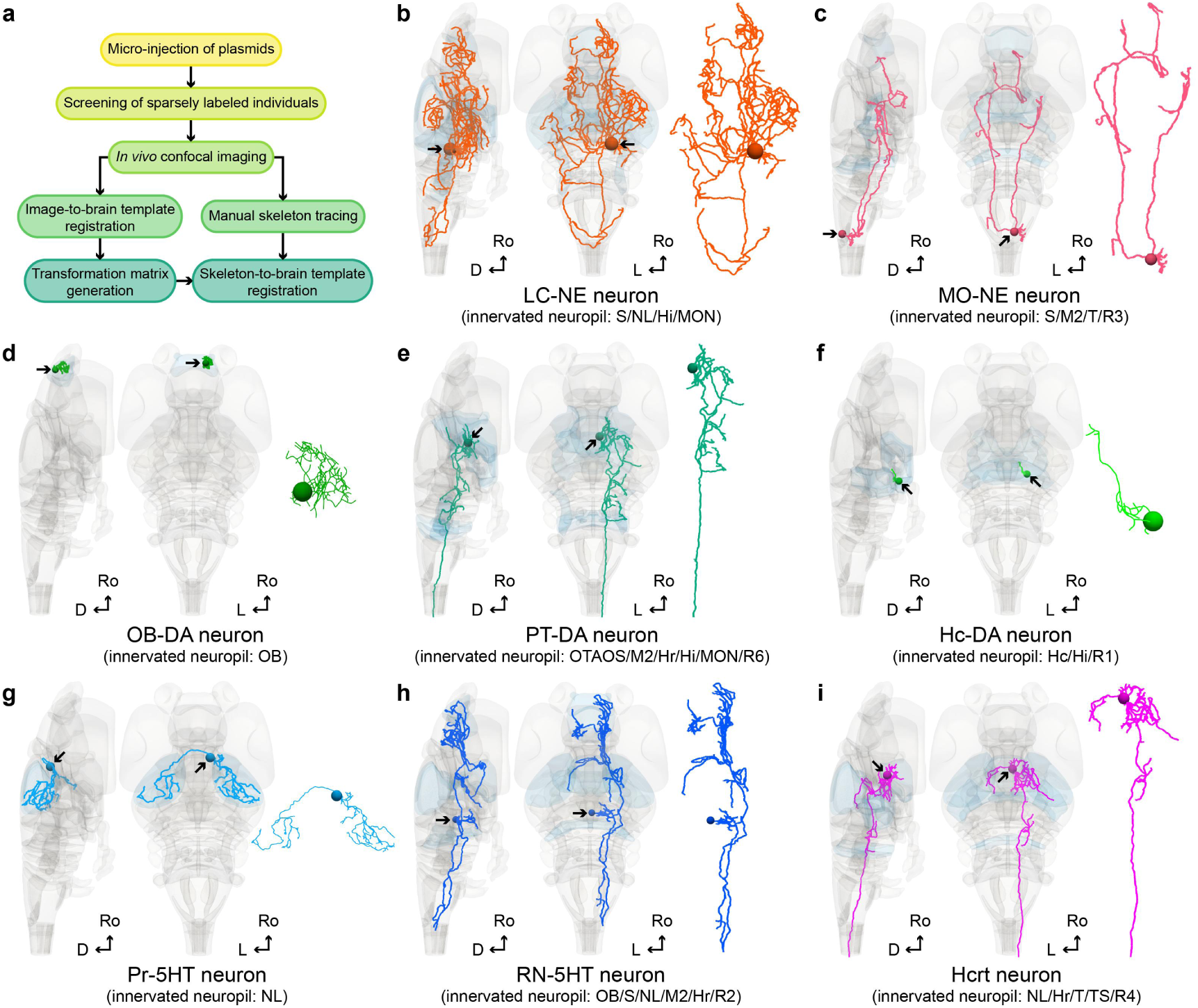
Single-cell morphology of diverse neuromodulatory neurons reconstructed by LM. **a,** Workflow for reconstructing single-cell morphologies by LM, adapted from Du et al^45^. Fluorescent protein-expressing plasmids were injected into fertilized eggs derived from neuromodulatory-type-specific Gal4 driver lines. This approach simultaneously enabled sparse labeling for morphology reconstruction and provided pan-neuronal fluorescence for image registration. At 3 dpf, larvae were screened by fluorescence microscopy to identify those with single-cell labeling. Selected larvae were then subjected to *in vivo* confocal imaging at 6 dpf. Neuronal morphologies were reconstructed through manual skeleton tracing based on the acquired fluorescent images. In parallel, the confocal images were registered to a mesoscopic brain template, and the resulting transformation matrices were applied to align the reconstructed skeletons with the standard reference space. **b**-**I,** Representative mesoscopic single-cell morphologies of diverse neuromodulatory types, shown in lateral (left), dorsal (middle), and detailed dorsal (right) views. Examples include LC-NE (**b**), MO-NE (**c**), OB-DA (**d**), PT-DA (**e**), Hc-DA (**f**), Pr-5HT (**g**), RN-5HT (**h**), and Hcrt (**i**) neurons. Black arrows indicate somata. The predominantly innervated neuropil (highlighted in light blue) is shown in parentheses. Ro, rostral; L, left; D, dorsal; L, left.

**Extended Data Fig. 6.**
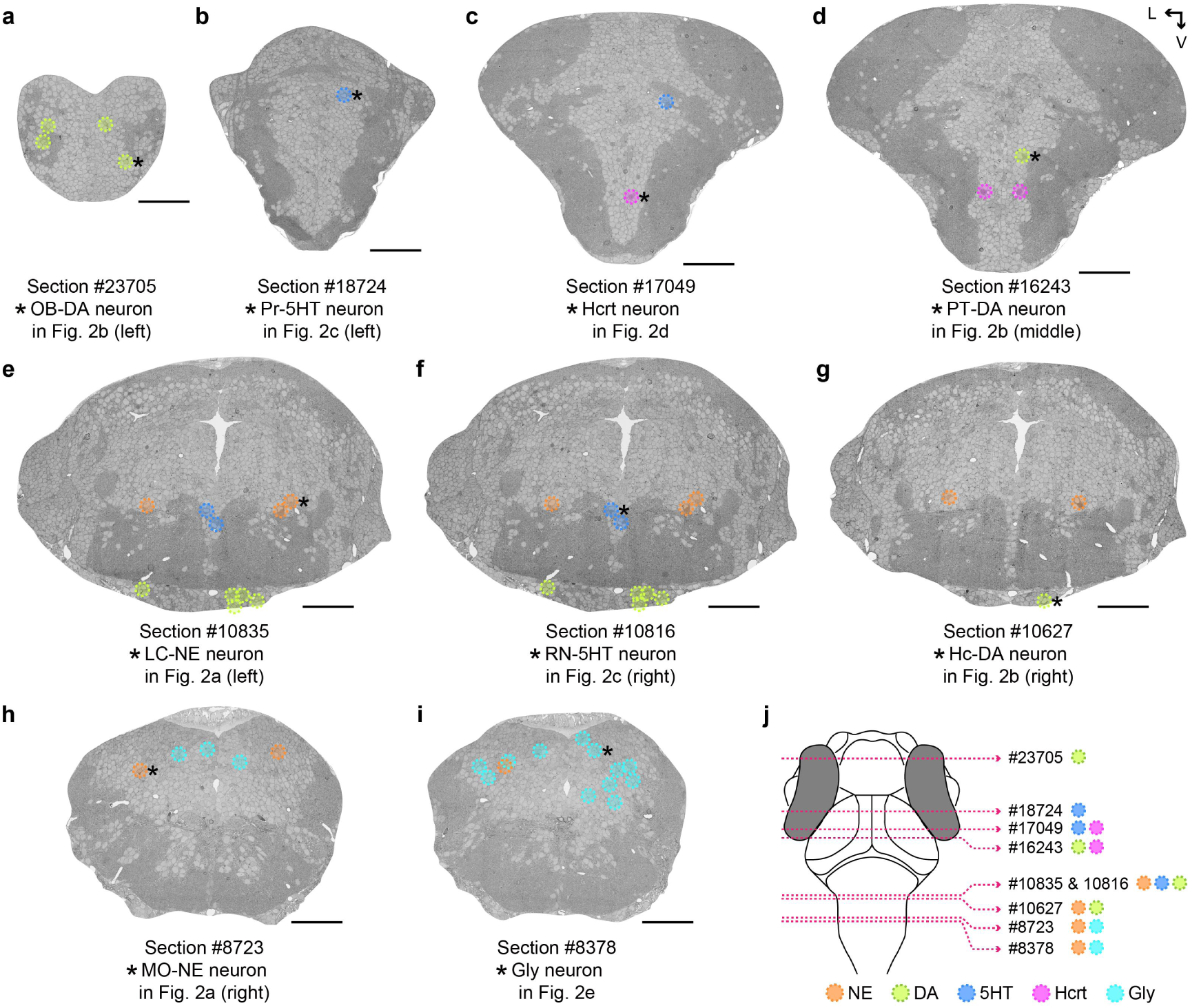
Neuron type identification based on soma location and APEX2 labeling patterns. **a-i,** Corresponding 33-nm EM sections (64×64 nm^2^ per pixel) containing the representative APEX2-labeled neurons (marked by asterisks) from Fig. 2a-e. All other APEX2-labeled neurons within the same sections are also outlined with color-coded circles (NE, orange; DA, green; 5HT, blue; Hcrt, magenta; Gly, cyan). Section indices are listed below each image. The sections are arranged sequentially from left to right and top to bottom, following the rostral-to-caudal axis. **j,** The schematic indicates the exact rostral-caudal positions of these sections, with corresponding section indices and neuronal types listed to the right. V, ventral; L, left. Scale bars, 50 µm (**a-i**).

**Extended Data Fig. 7.**
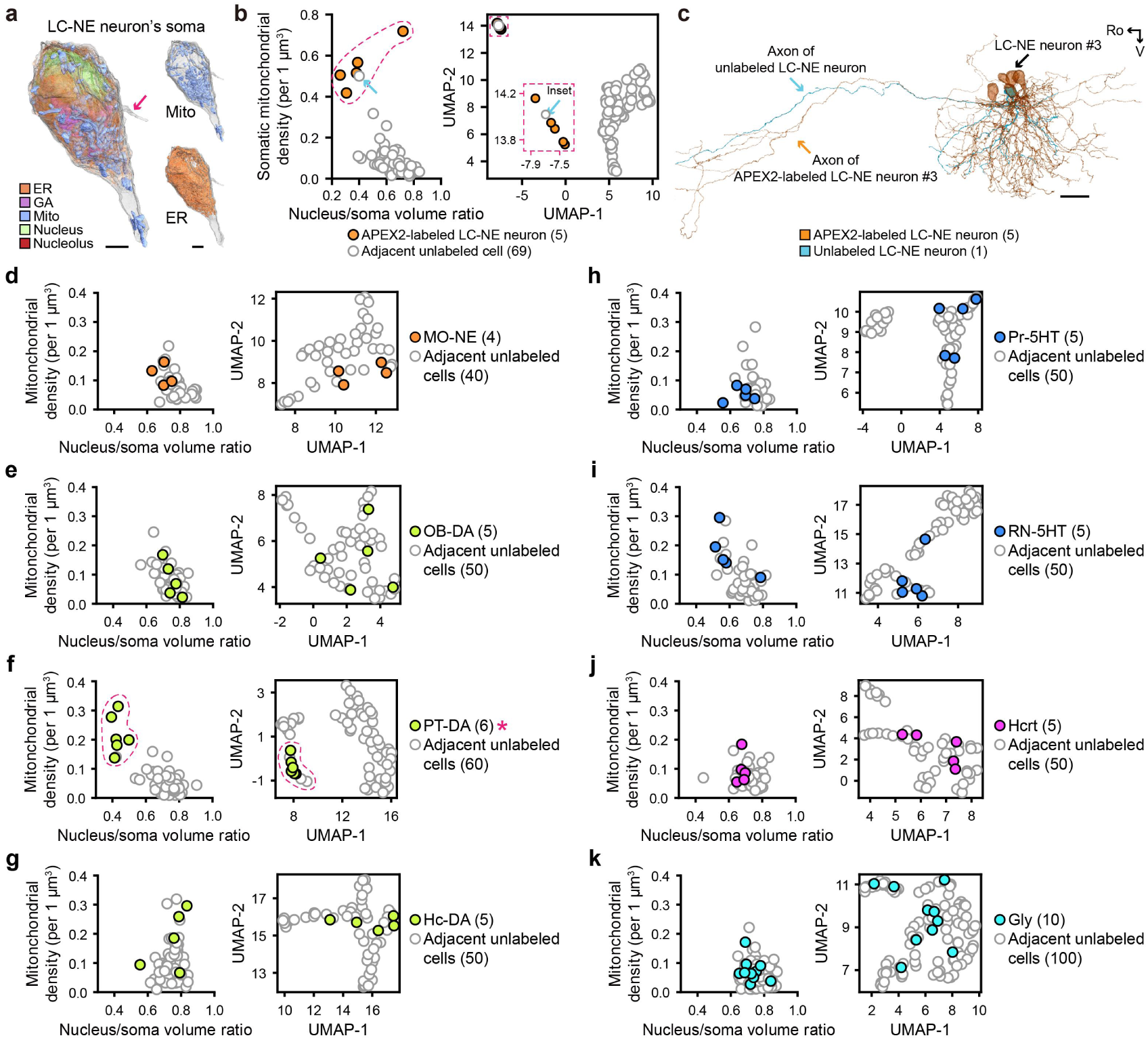
Comparison analysis of perisomatic features between neuromodulatory neurons and their adjacent unlabeled cells. **a,** Rendering of a single LC-NE neuron’s soma, containing endoplasmic reticulum (ER, orange), Golgi apparatus (GA, pink), mitochondria (Mito, blue), nucleus (green), and nucleolus (red). The red arrow indicates a primary cilia. For clarity, Mito (top right) and ER (bottom right) are additionally shown in separate renderings. **b,** Comparison of perisomatic features between APEX2-labeled LC-NE neurons (orange, *n* = 5) and their unlabeled neighbors (gray, *n* = 69). Labeled LC-NE neurons exhibited a lower nucleus-to-soma volume ratio and a higher density of somatic mitochondria compared to the randomly-selected neighboring cells (left). Based on a combination of deep-learned and traditional morphological features of the nucleus and soma, UMAP visualization revealed a consistent structural distinction between the two groups (right, inset). The blue arrow points to an unlabeled cell that clustered with LC-NE neurons and displayed similar properties to those of LC-NE neurons, suggesting it may be an unlabeled member of the same type. **c,** Rendering of the five APEX2-labeled (orange) and one unlabeled (blue) LC-NE neurons shown in **b**. Morphological reconstruction revealed similar axonal projection patterns between the LC-NE neuron #3 and unlabeled LC-NE neuron. **d**-**k,** Comparisons of perisomatic features between APEX2-labeled neurons and a 10-fold number of adjacent unlabeled cells. Left, nucleus-to-soma volume ratio *vs* somatic mitochondrial density. Right, UMAP visualization of cell clustering based on combined perisomatic morphological features. A similar structural divergence was observed for PT-DA neurons (red outline and asterisk, **f**), when compared to their neighbors. Cell numbers are shown in parentheses. Ro, rostral; V, ventral. Scale bars, 2 µm (**a**) and 20 µm (**c**).

**Extended Data Fig. 8.**
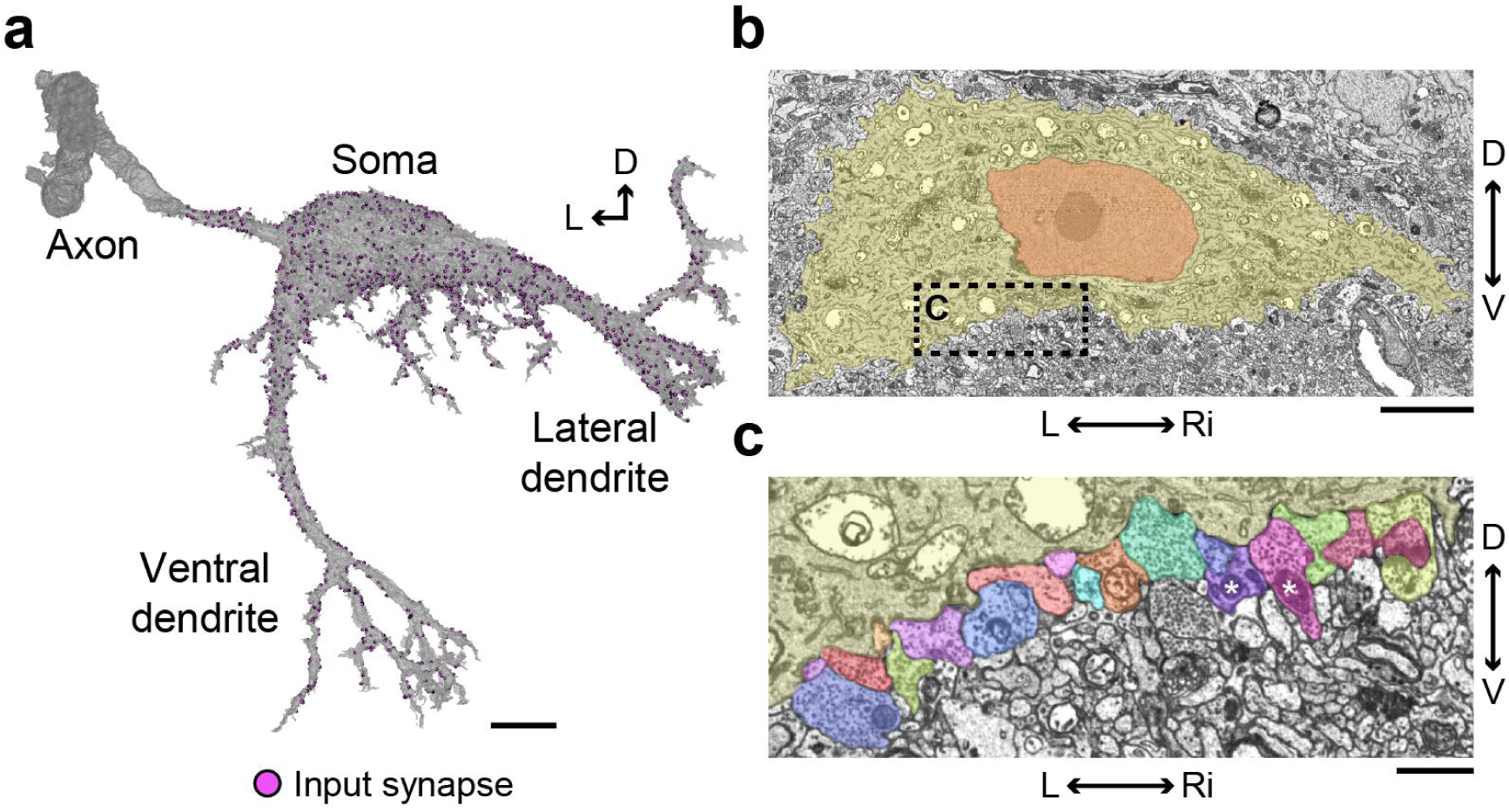
Somatodendritic reconstruction and synaptic input mapping of a Mauthner cell. **a,** Rendering of a right M-cell’s soma, lateral dendrites, ventral dendrites, and proximal axon. The axonal branch is not fully displayed. Small violet dots indicate synaptic input sites (*n* = 1,966). Both the morphology and synapses were manually verified based on automated segmentation results. **b,c,** Representative EM images (4×4 nm^2^ per pixel) of the M-cell’s soma, shown in coronal (**b**) and detailed coronal (**c**) views. The nucleus and cytosol are labeled in red and yellow, respectively. A typical region of the ventral cell surface was densely innervated by multiple presynaptic inputs with random color-coding. L, left; Ri, right; D, dorsal; V, ventral. Scale bars, 1 µm (**c**), 5 µm (**b**), and 10 µm (**a**).

**Extended Data Fig. 9.**
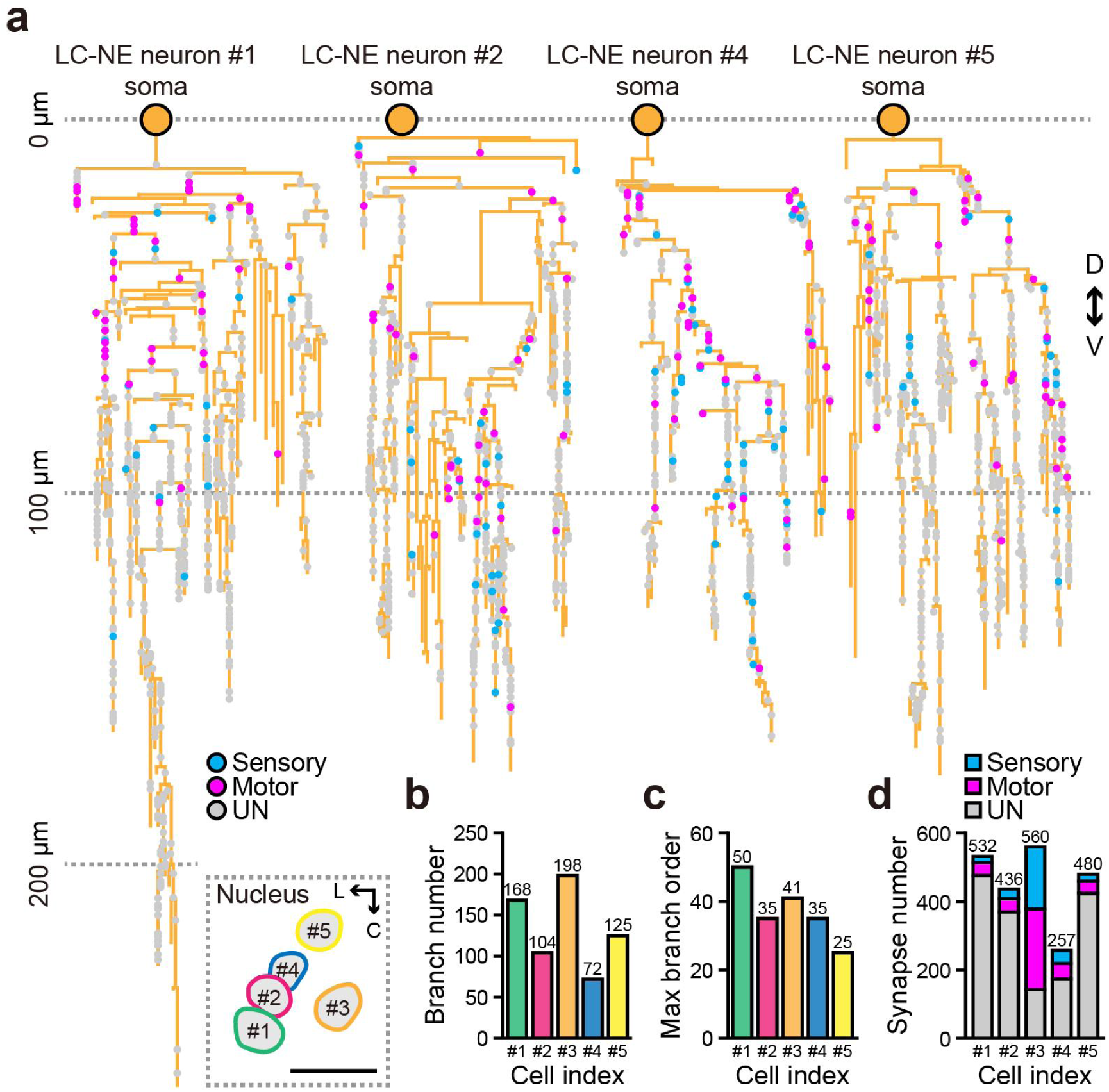
Dendritic architecture and synaptic inputs of the other four APEX2-labeled LC-NE neurons. **a,** 2D spatial distribution of dendritic synaptic inputs on LC-NE neurons (#1, #2, #4, and #5). Each dot represents a single synaptic input site, color-coded by the modality of PNs (Sensory-related, blue; Motor-related, magenta). Gray dots indicate synaptic sites untraced or traced but not yet reached soma (UN, undefined). The distance between each dot and the soma represents the path length of the corresponding synaptic input. Spatial distribution of five LC-NE neurons is shown via nuclear positioning in a dorsal view (bottom left). **b,c,** Dendritic heterogeneity across individual LC-NE neurons in both branch number (**b**) and maximum branch order (**c**). Values are listed above each bar. **d**, Dendritic synaptic input counts for individual LC-NE neurons, with values listed above each bar. Synapses of known modality are proportionally represented within each bar and are color-coded consistently with A. C, caudal; L, left; D, dorsal; V, ventral. Scale bars, 20 µm (**a**).

**Extended Data Fig. 10.**
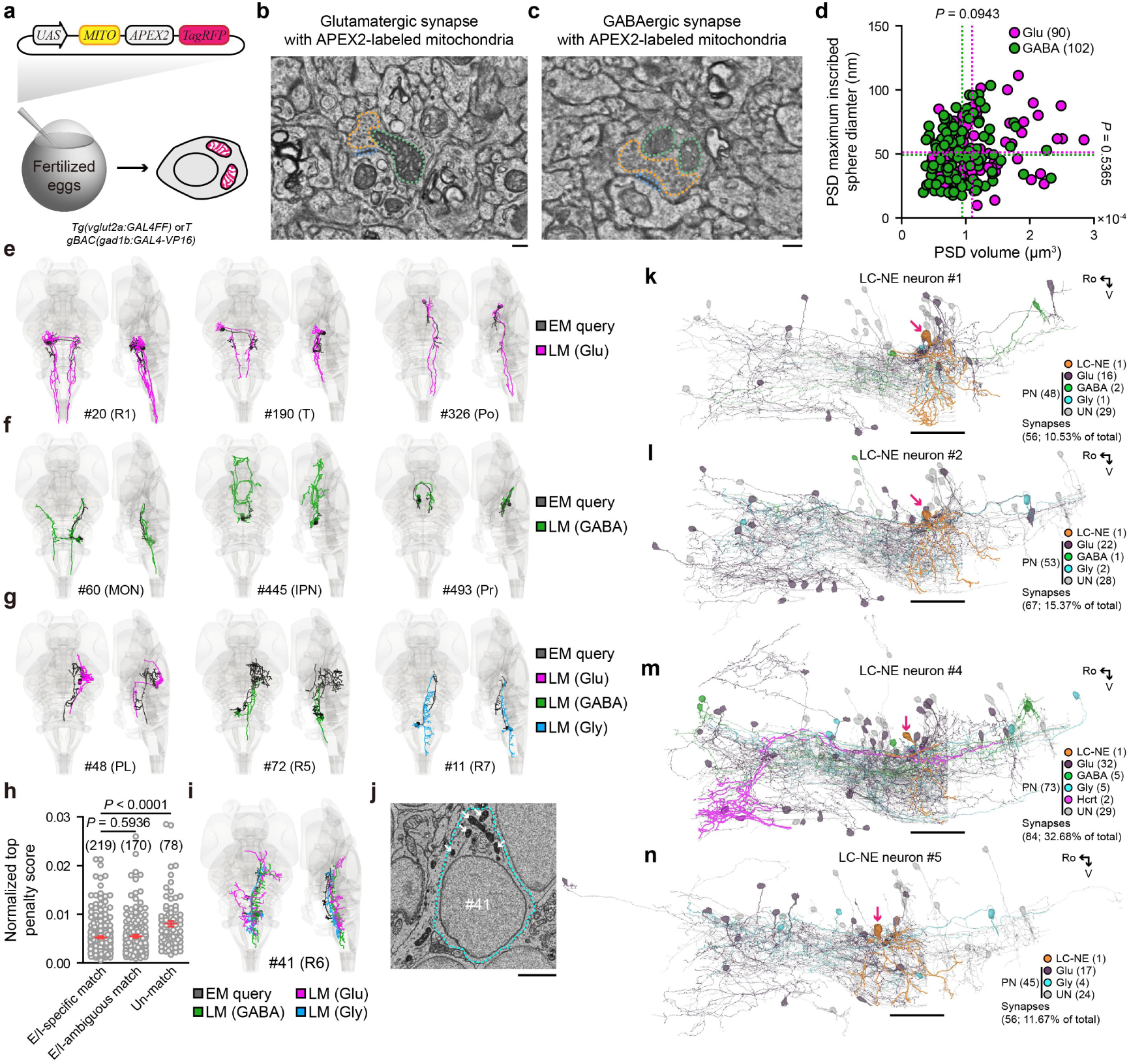
E/I type of reconstructed neurons can be identified via morphology comparison but not synaptic features. **a,** The plasmid *UAS:MITO-APEX2-TagRFP* was injected into fertilized eggs derived from *Tg(vglut2a:GAL4FF)* or *TgBAC(gad1b:GAL4-VP16)* lines to specifically label mitochondria in Glu or GABA neurons, respectively. **b**,**c** Representative EM images of APEX2-labeld mitochondria in axon terminals of Glu (**b**) and GABA (**c**) neurons. PSDs, blue; SVCs, orange; Labeled mitochondria, green. **d**, Comparisons of ultrastructural features for PSDs between Glu (magenta, *n* = 90) and GABA (green, *n* = 102) synapses. PSD thickness was estimated from the diameter of its maximum inscribed sphere. Dashed lines indicate the average values of PSD thickness (horizontal) and volume (vertical). *P*(thickness) = 0.5365, *P*(volume) = 0.0943 (Mann Whitney test). **e**-**g**, Representative examples showing partially EM-reconstructed query neurons (black) with their fully LM-reconstructed top-ranked E/I type counterparts (Glu, magenta; GABA, green; Gly, blue), revealed by morphology comparison. Some EM queries were matched to LM neurons with similar morphology (**e,f**), whereas others exhibit substantial morphological differences from their top-ranked LM references (**g**). All EM queries used in morphology comparison were PNs of LC-NE neurons. The soma location is shown in parentheses. **h,** Comparison of the normalized top penalty scores for all 467 EM queries, with lower penalty scores indicating higher degrees of matching. The penalty score was normalized by skeleton node number of EM queries. All results were manually verified after morphological matching and categorized into E/I-specific match, E/I-ambiguous match, and Un-match. In some brain regions, E and I neurons exhibit similar morphology, causing good matches between EM queries and both E/I neurons. Cell numbers are shown in parentheses. The Mann Whitney test was used, and the error bars represent mean ± SEM. **i**,**j**, EM query neuron #41 (black) exhibited morphological similarity to adjacent LM-reconstructed Glu, GABA, and Gly neurons (**i**), suggesting ambiguous identity based on structure alone. However, it could be unambiguously identified as Gly by its mitochondrial-APEX2 labeling, as indicated by white arrows (**j**). **k-n,** Rendering of EM-reconstructed PNs and corresponding LC-NE neuron #1 (**k**), #2 (**l**), #4 (**m**), and #5 (**n**). The counts of PNs with E/I profiles and their corresponding synaptic outputs onto the target LC-NE neuron are shown in parentheses. Notably, two Hcrt neurons (magenta) with nuclear-localized APEX2 labeling specifically targeted the LC-NE neuron #4 (**m**), rather than other LC-NE neurons. Ro, rostral; V, ventral. Scale bars, 200 nm (**b,c**), 2 µm (**j**), and 50 µm (**k-n**).

## SUPPLEMENTARY INFORMATION

### SUPPLEMENTARY VIDEOS

**Supplementary Video 1 |** Fish-X: A multidimensional whole-brain EM reconstruction of a 6-dpf larva characterized by multiplexed APEX2-based neuromodulatory type labeling, comprising 529 reconstructed neurons of various types, 176,810 nuclei, neural tracts, and vessels in the brain and anterior spinal cord. A *Gt(vmat2:GAL4FF);Tg(dbh:GAL4FF;UAS:cytosol-APEX2-TagRFP);Tg(hcrt:APEX2-TagRFP-NLS);Tg(glyt2:MITO-APEX2-TagRFP)* larva at 6 dpf was used.

**Supplementary Video 2 |** The whole-brain density distribution for nuclei, PSDs, SVCs, and Mito. The sequence first shows brain regions emerging along the rostral-to-caudal axis, then displays the corresponding cellular components in descending order of density.

**Supplementary Video 3 |** Somatodendritic reconstruction of multiple neuromodulatory and Gly neuronal populations.

**Supplementary Video 4 |** Dendritic input site mapping of multiple monoaminergic neuronal populations.

**Supplementary Video 5 |** Mauthner cell reconstruction with dendritic input site mapping.

**Supplementary Video 6 |** Widely distributed inputs from 374 PNs across 32 brain regions converging onto the LC-NE neuron #3.

**Supplementary Video 7 |** The spatial organization of synaptic inputs to the LC-NE neuron #3. Inputs are categorized as follows: 1) modalities (sensory-/motor-related), 2) E/I types (Glu/GABA/Gly), 3) shared inputs within the LC-NE system (intra-CPNs), and 4) shared inputs across monoaminergic systems (inter-CPNs).

### SUPPLEMENTARY TABLES

**Supplementary Table 1.** Genetic toolbox for subcellularly-targeted APEX2 labeling.

| No. | Full name | Simple name | Targeting | Type | Source |
| --- | --- | --- | --- | --- | --- |
| 1 | *Tg(dbh:GAL4FF;5×UAS-hsp:APEX2-TagRFP-NLS)ion143d* | *Tg(dbh:GAL4FF;UAS:APEX2-TagRFP-NLS)* | Nucleus | NE | Line |
| 2 | *Tg(dbh:GAL4FF;5×UAS-hsp:cytosol-APEX2-TagRFP)ion142d* | *Tg(dbh:GAL4FF;UAS:cytosol-APEX2-TagRFP)* | Cytosol | NE | Line |
| 3 | *Tg(hcrt:APEX2-TagRFP-NLS)ion100d* | *Tg(hcrt:APEX2-TagRFP-NLS)* | Nucleus | Hcrt | Line |
| 4 | *Tg(hcrt:MITO-APEX2-TagRFP)ion101d* | *Tg(hcrt:MITO-APEX2-TagRFP)* | Mitochondria | Hcrt | Line |
| 5 | *Ki(slc17a6b:GAL4FF,5×UAS-hsp:sfGFP-APEX2-NLS)ion214d* | *Ki(vglut2a:GAL4FF,UAS:sfGFP-APEX2-NLS)* | Nucleus | Glu | Line |
| 6 | *TgBAC(slc17a6:cytosol-APEX2-TagRFP)ion140d* | *TgBAC(vglut2a:cytosol-APEX2-TagRFP)* | Cytosol | Glu | Line |
| 7 | *TgBAC(gad1b:Hsa.Cox8a-APEX2-P2A-EGFP)ion170d* | *TgBAC(gad1b:MITO-APEX2-P2A-EGFP)* | Mitochondria | GABA | Line |
| 8 | *TgBAC(Gad1b:APEX2-EGFP-CAAX)ion141d* | *TgBAC(gad1b:APEX2-EGFP-CAAX)* | Plasma membrane | GABA | Line |
| 9 | *Ki(Gad1b:SEPT6-sfGFP-APEX2)ion202d* | *Ki(gad1b:SEPT6-sfGFP-APEX2)* | Fibrous in cytosol | GABA | Line |
| 10 | *Tg(slc6a5:APEX2-TagRFP-NLS)ion99d* | *Tg(glyt2:APEX2-TagRFP-NLS)* | Nucleus | Gly | Line |
| 11 | *Tg(slc6a5:Hsa.Cox8a-APEX2-TagRFP)ion97d* | *Tg(glyt2:MITO-APEX2-TagRFP)* | Mitochondria | Gly | Line |
| 12 | *Tg(5×UAS-hsp:APEX2-TagRFP-NLS)ion195d* | *Tg(UAS:APEX2-TagRFP-NLS)* | Nucleus | - | Line |
| 13 | *Tg(5×UAS-hsp:sypb-APEX2-TagRFP)ion144d* | *Tg(UAS:sypb-APEX2-TagRFP)* | Presynaptic vesicle | - | Line |
| 14 | - | *yTol2-dbh:GAL4FF,UAS:SEPT6-sfGFP-APEX2-NLS* | Fibrous in cytosol | DBH | Plasmid |
| 15 | - | *yTol2-hcrt:Golgi-APEX2-TagRFP* | Golgi apparatus | Hcrt | Plasmid |
| 16 | - | *pMD19-T-gad1b-P2A-tetoff-SEPT6-sfGFP-APEX2-NLS KI donor* | Fibrous in cytosol | GABA | Plasmid |
| 17 | - | *yTol2-UAS:MITO-APEX2-TagRFP* | Mitochondria | - | Plasmid |
| 18 | - | *yTol2-UAS:cytosol-PRDX4m*<br>*-sfGFP-APEX2* | Spherical in<br>cytosol | - | Plasmid |
| 1986 | Transgenic zebrafish larvae were in *nacre* background. All transgenic lines and<br>plasmids were first published in this paper. |  |  |  |  |
| 1987 |  |  |  |  |  |
| 1988 |  |  |  |  |  |

**Supplementary Table 2.**
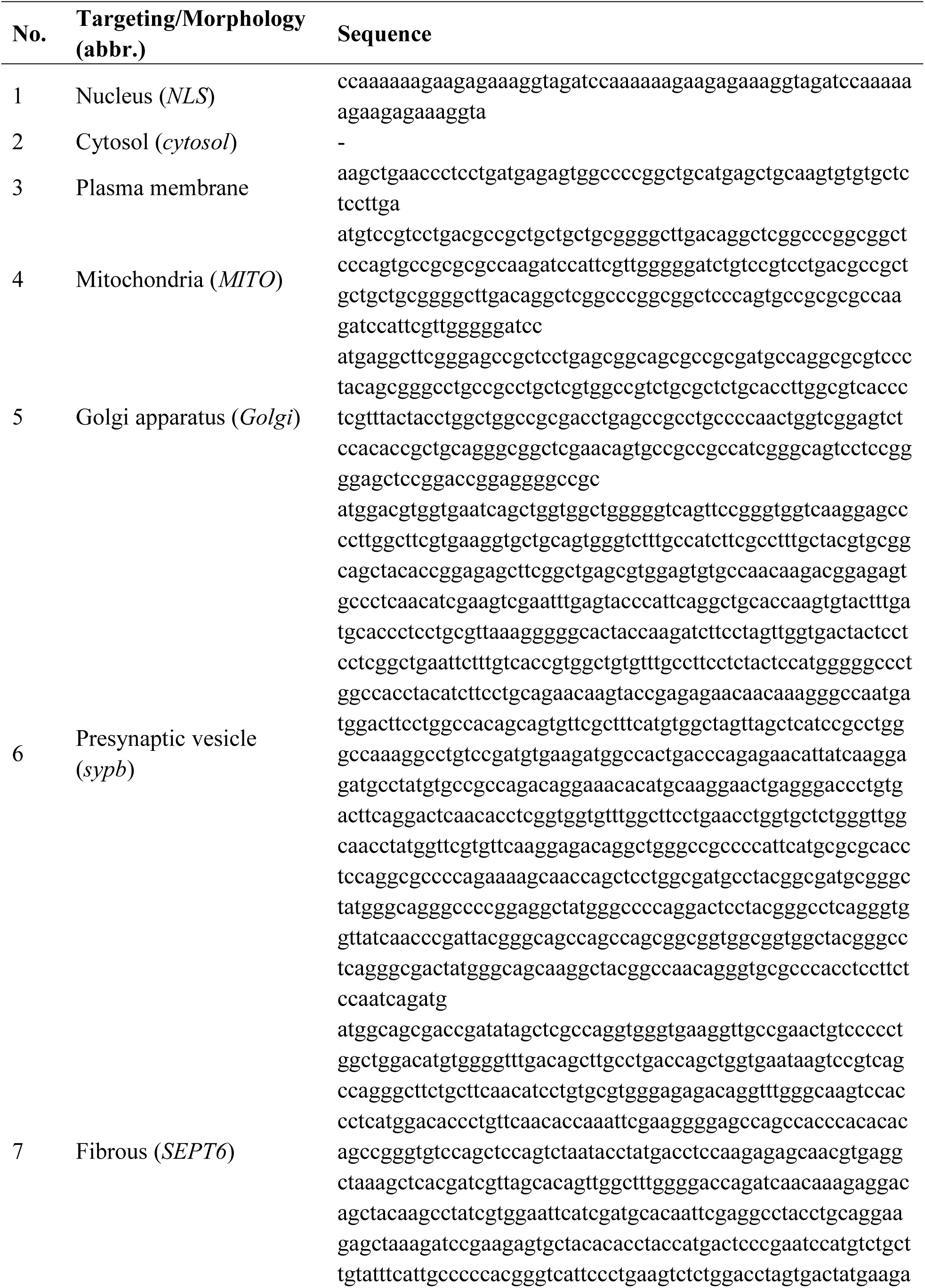
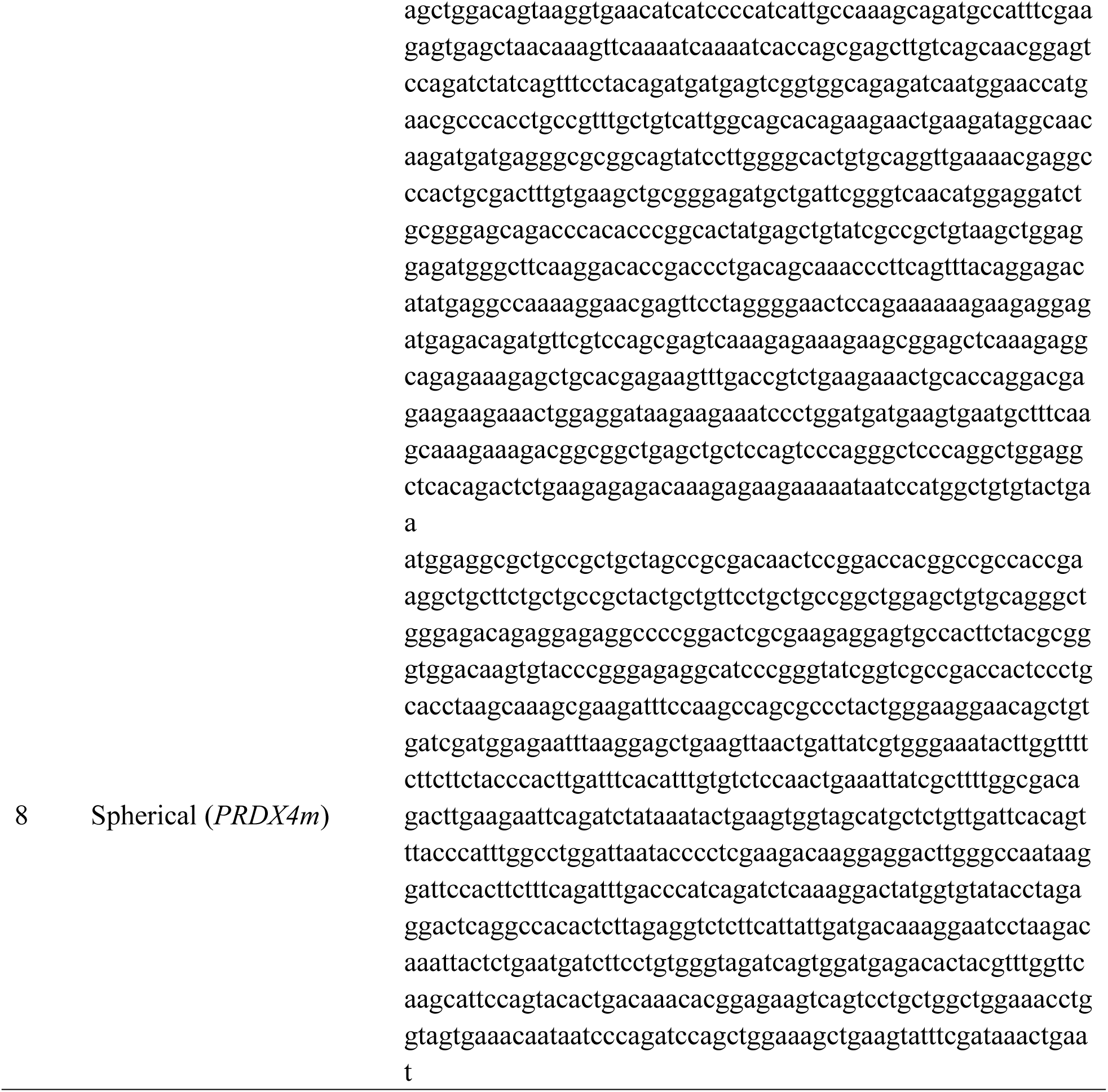
Signal peptide sequences for subcellular targeting and self-assembly of APEX2.

**Supplementary Table 3.** Standardized nomenclature of brain regions.

| No. | Name | Abbr. | Volume<br>($\mu\text{m}^3$) |
| --- | --- | --- | --- |
| 1 | Olfactory bulb | OB | 552,907.80 |
| 2 | Pallium | P | 1,176,906.63 |
| 3 | Subpallium | S | 954,796.37 |
| 4 | Preoptic region | Po | 759,356.45 |
| 5 | Optic tract and accessory optic system | OTAOS | 986,748.95 |
| 6 | Pretectum | Pr | 669,170.64 |
| 7 | Epiphysis (Pineal gland) | Ep | 110,533.43 |
| 8 | Habenula | Ha | 289,854.83 |
| 9 | Dorsal thalamus | DT | 286,673.99 |
| 10 | Ventral thalamus | VT | 336,431.48 |
| 11 | Posterior tuberculum | PT | 377,980.15 |
| 12 | Migrated posterior tubercular region | M2 | 658,202.84 |
| 13 | Rostral hypothalamus | Hr | 1,204,135.20 |
| 14 | Intermediate hypothalamus | Hi | 1,874,093.02 |
| 15 | Diffuse nucleus of the intermediate hypothalamus | DHi | 250,549.79 |
| 16 | Caudal hypothalamus | Hc | 176,238.95 |
| 17 | Pituitary | Pit | 83,580.57 |
| 18 | Neuropil layer (optic tectum) | NL | 2,939,375.09 |
| 19 | Periventricular cell layer (optic tectum) | PL | 3,760,502.17 |
| 20 | Tegmentum | T | 536,864.29 |
| 21 | Nucleus of medial longitudinal fascicle | nMLF | 30,611.07 |
| 22 | Oculomotor nerve nucleus | nIII | 77,251.04 |
| 23 | Torus longitudinalis | TL | 115,042.14 |
| 24 | Torus semicircularis | TS | 1,105,957.84 |
| 25 | Rhombomere 1 | R1 | 1,829,045.90 |
| 26 | Trochlear nerve motor nucleus | nIV | 50,476.46 |
| 27 | Interpeduncular nucleus | IPN | 196,849.62 |
| 28 | Superior Raphe | SR | 122,206.58 |
| 29 | Corpus cerebelli | CCe | 1,345,996.74 |
| 30 | Valvula cerebelli | Va | 282,478.01 |
| 31 | Eminentia granularis | EG | 270,844.81 |
| 32 | Lobus caudalis cerebelli | LCa | 32,741.70 |
| 33 | Medial octavolateralis nucleus | MON | 274,799.18 |
| 34 | Rhombomere 2 | R2 | 1,039,406.70 |
| 35 | Dorsal motor nucleus of trigeminal nerve | nVd | 36,112.51 |
| 36 | Rhombomere 3 | R3 | 897,971.87 |
| 37 | Ventral motor nucleus of trigeminal nerve | nVv | 16,845.30 |
| 38 | Inferior raphe | IR | 32,603.36 |
| 39 | Rhombomere 4 | R4 | 891,197.12 |
| 40 | Rhombomere 5 | R5 | 1,038,459.75 |
| 41 | Medial vestibular nucleus | MVN | 35,611.76 |
| 42 | Tangential vestibular nucleus | TVN | 30,629.58 |
| 43 | Rostral facial nerve motor nucleus and octavolateralis efferent neurons | nVIIr/OLe<br>r | 68,704.18 |
| 44 | Rostral motor nucleus of abducens nerve | nVIr | 6,896.53 |
| 45 | Rhombomere 6 | R6 | 1,048,429.95 |
| 46 | Caudal facial nerve motor nucleus and octavolateralis efferent neurons | nVIIc/OLe<br>c | 22,639.00 |
| 47 | Caudal motor nucleus of abducens nerve | nVIc | 14,460.40 |
| 48 | Rhombomere 7 | R7 | 930,447.61 |
| 49 | Rhombomere 8 | R8 | 1,964,286.62 |
| 50 | Glossopharyngeal/Vagal nerve motor nucleus | nIX/X | 188,459.61 |
| 51 | Vagal lobe | VLo | 15,311.87 |
| 52 | Inferior olive | IO | 37,314.70 |
| 53 | Outer nuclear layer | ONL | 1,935,701.58 |
| 54 | Outer plexiform layer | OPL | 218,800.10 |
| 55 | Inner nuclear layer | INL | 1,811,315.73 |
| 56 | Inner plexiform layer | IPL | 642,129.04 |
| 57 | Retinal ganglion cell layer | RGCL | 447,158.97 |
| 58 | Ciliary marginal zone | CMZ | 88,195.60 |
| 59 | Optic nerve | ON | 8,981.38 |

**Supplementary Table 4.**
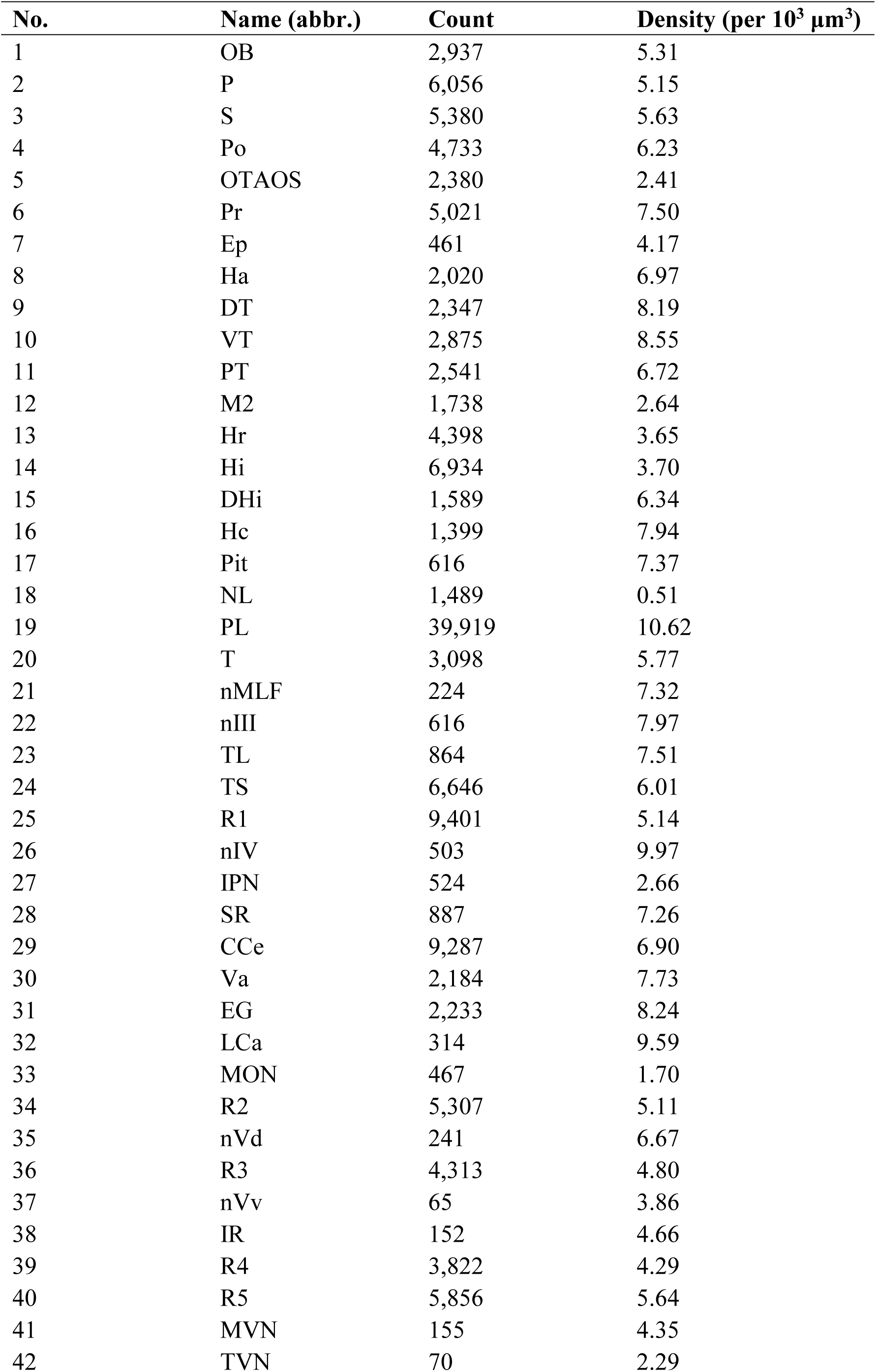
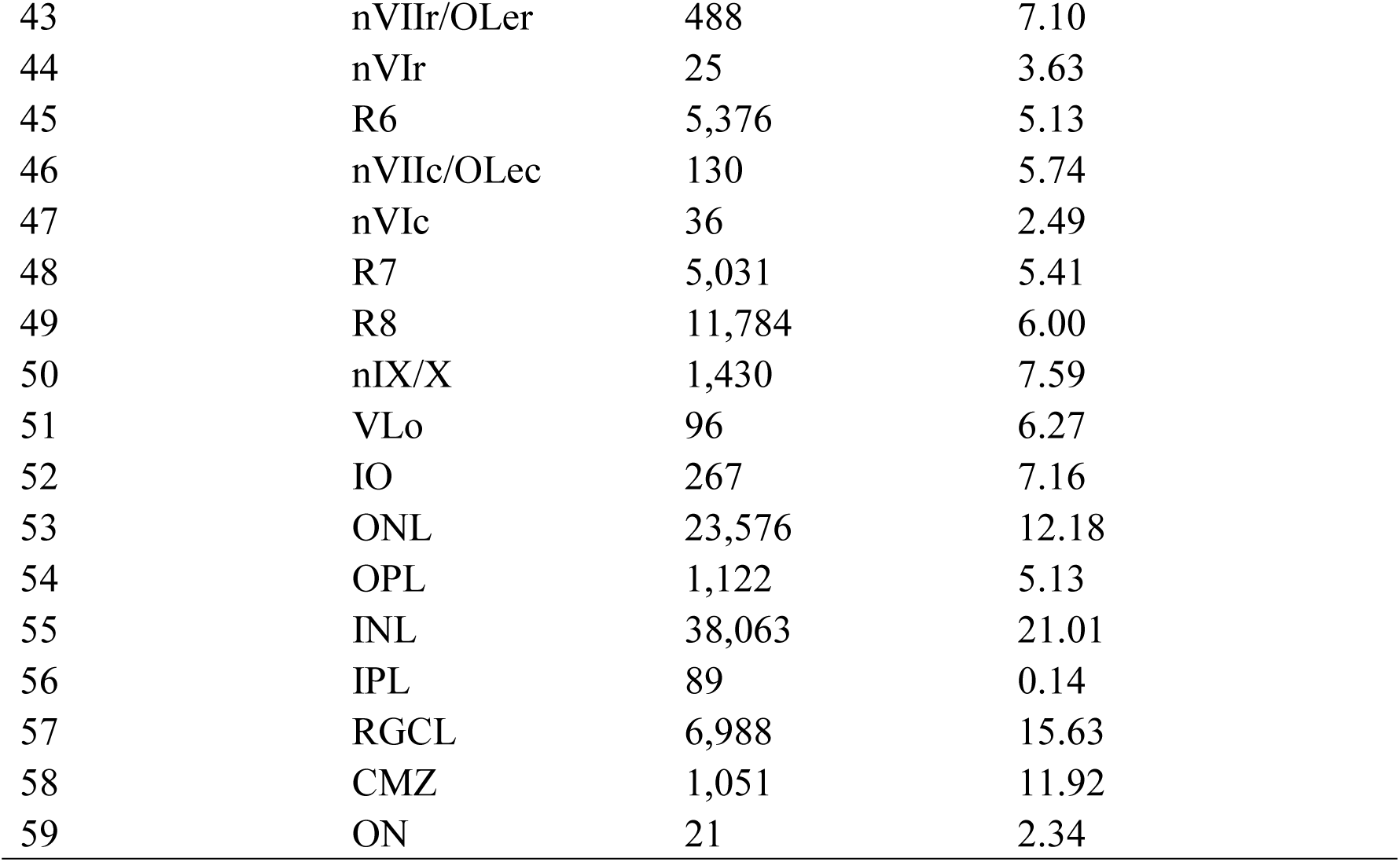
Nucleus count and density across brain regions.

**Supplementary Table 5.**
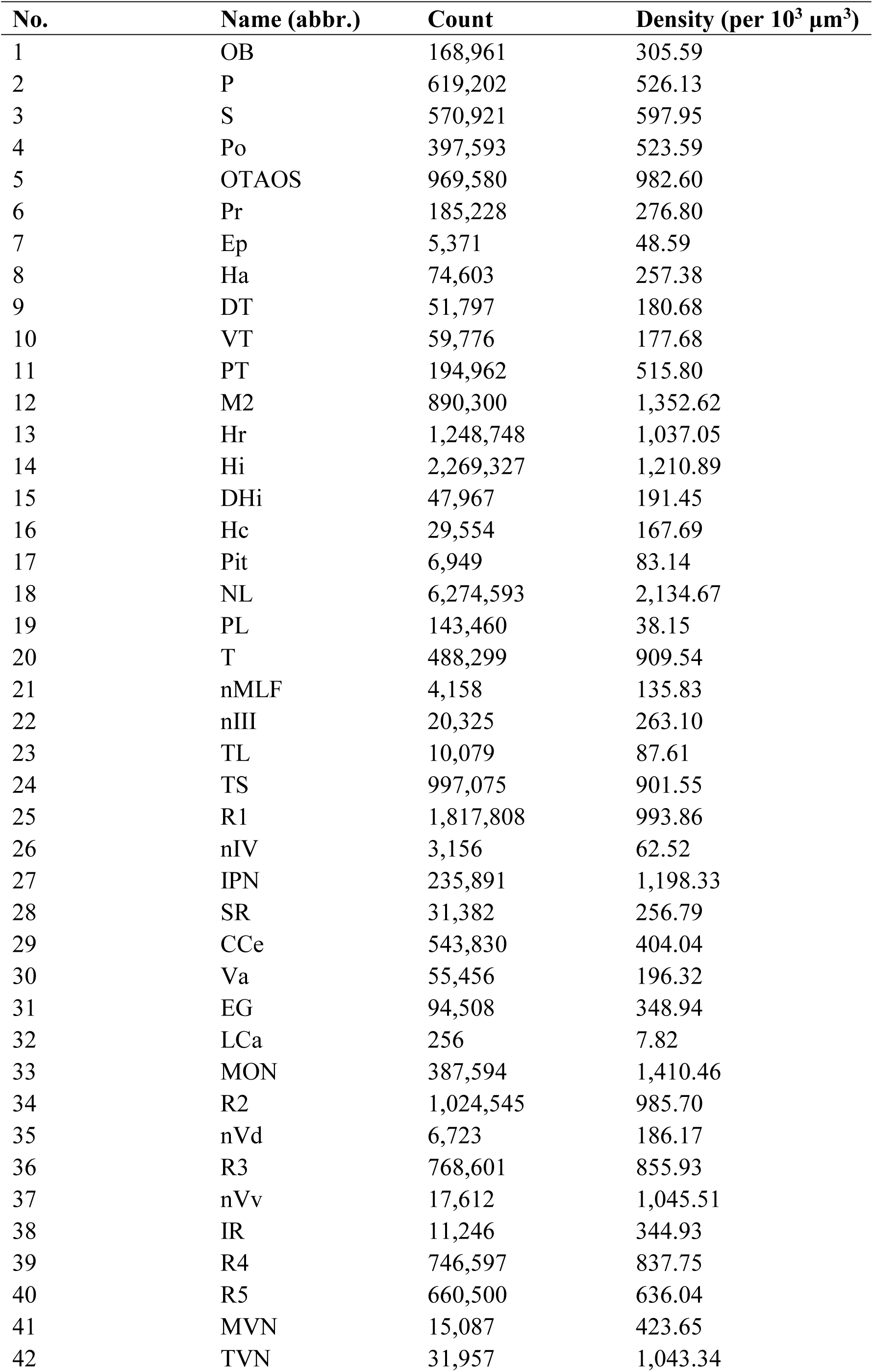
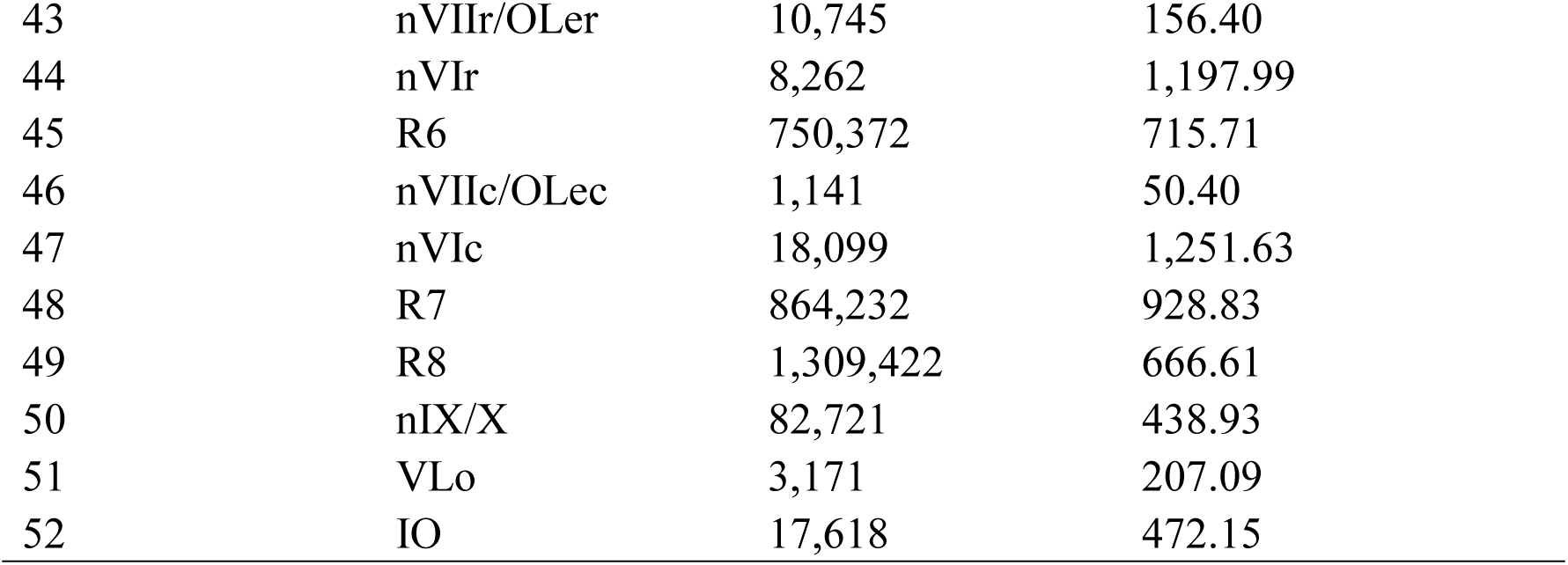
PSD count and density across brain regions.

**Supplementary Table 6.**
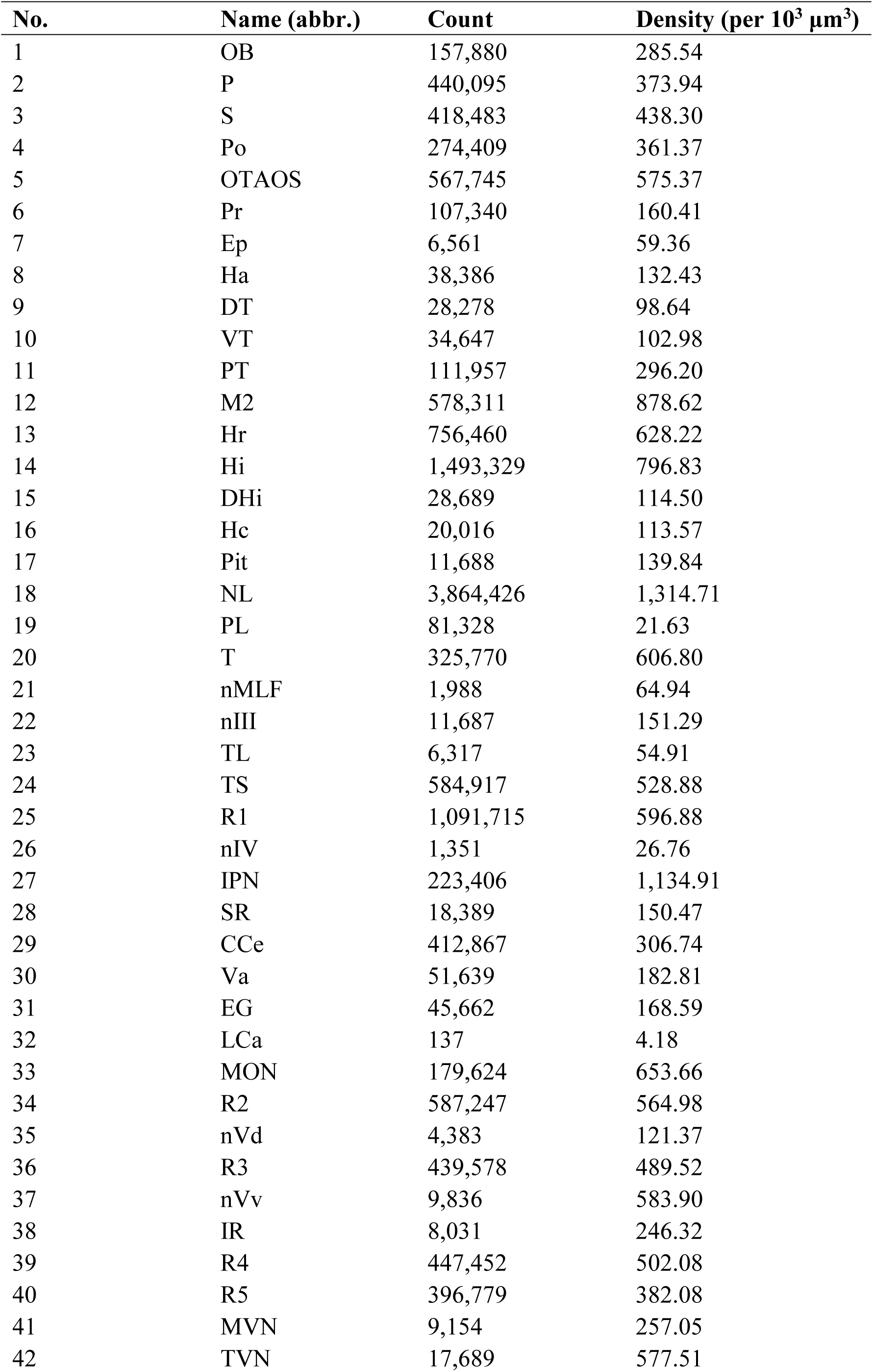
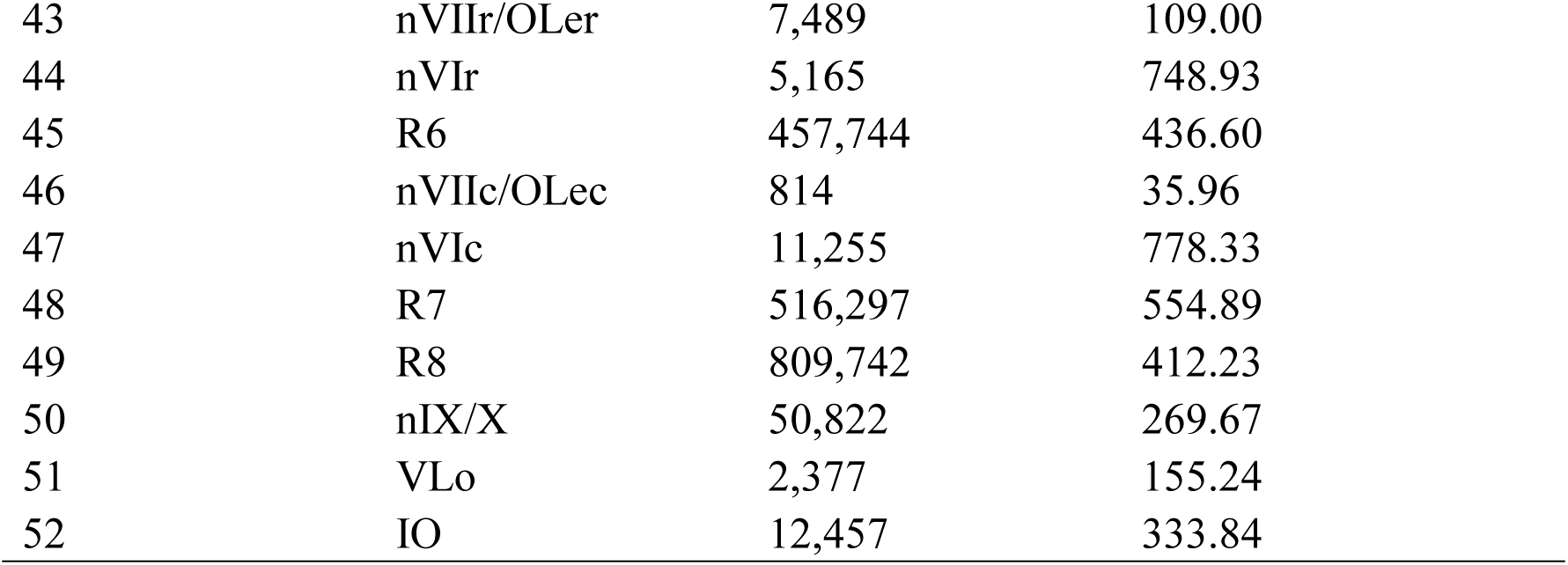
SVC count and density across brain regions.

**Supplementary Table 7.** Mito count and density across brain regions.

| No. | Name (abbr.) | Count | Density (per 10<sup>3</sup> μm<sup>3</sup>) |
| --- | --- | --- | --- |
| 1 | OB | 146,644 | 265.22 |
| 2 | P | 365,682 | 310.71 |
| 3 | S | 364,324 | 381.57 |
| 4 | Po | 247,663 | 326.15 |
| 5 | OTAOS | 535,443 | 542.63 |
| 6 | Pr | 133,626 | 199.69 |
| 7 | Ep | 11,706 | 105.90 |
| 8 | Ha | 58,557 | 202.02 |
| 9 | DT | 45,091 | 157.29 |
| 10 | VT | 47,103 | 140.01 |
| 11 | PT | 107,722 | 284.99 |
| 12 | M2 | 411,387 | 625.02 |
| 13 | Hr | 617,761 | 513.03 |
| 14 | Hi | 1,115,733 | 595.35 |
| 15 | DHi | 30,243 | 120.71 |
| 16 | Hc | 34,030 | 193.09 |
| 17 | Pit | 17,205 | 205.85 |
| 18 | NL | 2,582,241 | 878.50 |
| 19 | PL | 265,164 | 70.51 |
| 20 | T | 303,911 | 566.09 |
| 21 | nMLF | 3,942 | 128.78 |
| 22 | nIII | 18,408 | 238.29 |
| 23 | TL | 15,109 | 131.33 |
| 24 | TS | 509,154 | 460.37 |
| 25 | R1 | 905,849 | 495.26 |
| 26 | nIV | 4,828 | 95.65 |
| 27 | IPN | 217,883 | 1,106.85 |
| 28 | SR | 27,805 | 227.52 |
| 29 | CCe | 399,534 | 296.83 |
| 30 | Va | 65,467 | 231.76 |
| 31 | EG | 73,700 | 272.11 |
| 32 | LCa | 1,190 | 36.35 |
| 33 | MON | 251,781 | 916.24 |
| 34 | R2 | 490,509 | 471.91 |
| 35 | nVd | 6,031 | 167.01 |
| 36 | R3 | 402,701 | 448.46 |
| 37 | nVv | 6,147 | 364.91 |
| 38 | IR | 13,723 | 420.91 |
| 39 | R4 | 427,100 | 479.24 |
| 40 | R5 | 413,002 | 397.71 |
| 41 | MVN | 12,173 | 341.83 |
| 42 | TVN | 25,790 | 842.00 |
| 43 | nVIIr/OLer | 14,259 | 207.54 |
| 44 | nVIr | 4,869 | 706.01 |
| 45 | R6 | 497,382 | 474.41 |
| 46 | nVIIc/OLec | 5,806 | 256.46 |
| 47 | nVIc | 12,623 | 872.94 |
| 48 | R7 | 487,363 | 523.79 |
| 49 | R8 | 866,776 | 441.27 |
| 50 | nIX/X | 54,772 | 290.63 |
| 51 | VLo | 3,139 | 205.00 |
| 52 | IO | 15,150 | 406.01 |

**Supplementary Table 8.**
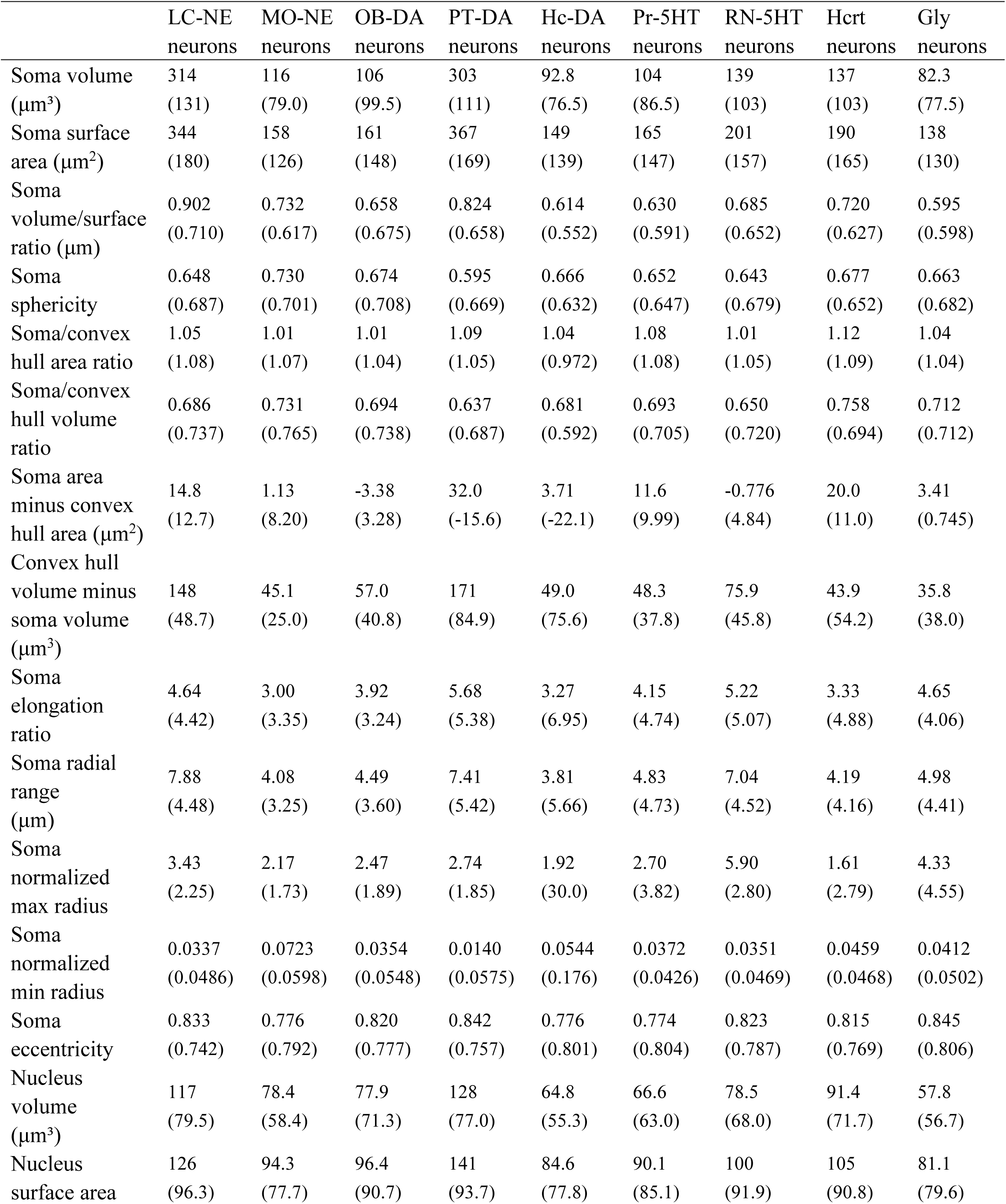
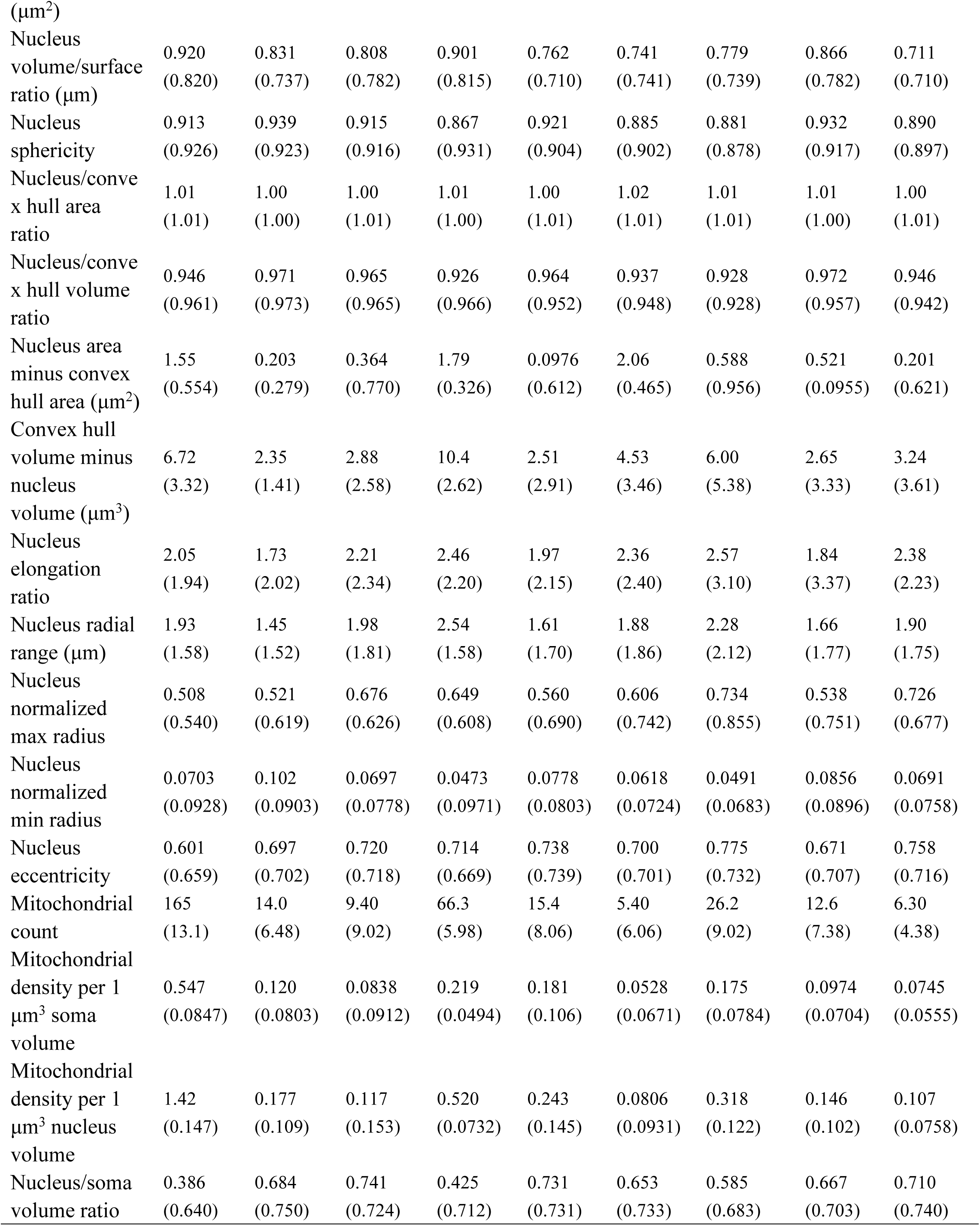
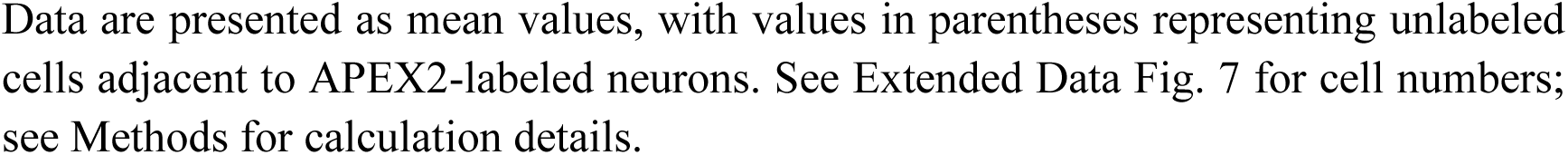
Perisomatic features of multiple APEX2-labeled neurons.

